# Regulation of Lung Immune Tone by the Gut-Lung Axis via Dietary Fiber, Gut Microbiota, and Short-Chain Fatty Acids

**DOI:** 10.1101/2023.08.24.552964

**Authors:** Daisuke Maruyama, Wen-I Liao, Xiaoli Tian, Marius Bredon, Johannes Knapp, Christine Tat, Thien N. M. Doan, Benoit Chassaing, Aditi Bhargava, Harry Sokol, Arun Prakash

**Affiliations:** Department of Anesthesia and Perioperative Care, University of California San Francisco and San Francisco General Hospital, San Francisco, CA

**Keywords:** Gut-lung Axis, SCFA, FFAR, gut microbiome, lung immune tone, lung injury, metabolites

## Abstract

Lung immune tone, i.e. the immune state of the lung, can vary between individuals and over a single individual’s lifetime, and its basis and regulation in the context of inflammatory responses to injury is poorly understood. The gut microbiome, through the gut-lung axis, can influence lung injury outcomes but how the diet and microbiota affect lung immune tone is also unclear. We hypothesized that lung immune tone would be influenced by the presence of fiber-fermenting short-chain fatty acid (SCFA)-producing gut bacteria. To test this hypothesis, we conducted a fiber diet intervention study followed by lung injury in mice and profiled gut microbiota using 16S sequencing, metabolomics, and lung immune tone. We also studied germ-free mice to evaluate lung immune tone in the absence of microbiota and performed *in vitro* mechanistic studies on immune tone and metabolic programming of alveolar macrophages exposed to the SCFA propionate (C3). Mice on high-fiber diet were protected from sterile lung injury compared to mice on a fiber-free diet. This protection strongly correlated with lower lung immune tone, elevated propionate levels and enrichment of specific fecal microbiota taxa; conversely, lower levels of SCFAs and an increase in other fatty acid metabolites and bacterial taxa correlated with increased lung immune tone and increased lung injury in the fiber-free group. *In vitro*, C3 reduced lung alveolar macrophage immune tone (through suppression of IL-1β and IL-18) and metabolically reprogrammed them (switching from glycolysis to oxidative phosphorylation after LPS challenge). Overall, our findings reveal that the gut-lung axis, through dietary fiber intake and enrichment of SCFA-producing gut bacteria, can regulate innate lung immune tone via IL-1β and IL-18 pathways. These results provide a rationale for the therapeutic development of dietary interventions to preserve or enhance specific aspects of host lung immunity.

## INTRODUCTION

The influence of the gut microbiome on human health and disease has been well established(Clemente et al., 2018; Erturk-Hasdemir & Kasper, 2013; Koh et al., 2016; Shukla et al., 2017), yet how gut microbiota control lung immune tone via the gut-lung axis is not well understood(Marsland et al., 2015; Samuelson et al., 2015; Shukla et al., 2017). Fiber-rich diets have been demonstrated to favor the expansion of fermenting gut bacteria resulting in the production of key metabolites, including short-chain fatty acids (SCFA). These SCFAs are recognized regulators of gut health and intestinal epithelial permeability and SCFAs have been implicated in modulating disease states throughout the body(Corrêa-Oliveira et al., 2016; Fusco et al., 2023; Koh et al., 2016). Previous reports have implicated SCFAs as having indirect or direct effects on asthma severity and bone health (Lucas et al., 2018; Trompette et al., 2014). More recent reports have implicated specific fiber sources as skewing immune responses towards type 2 inflammation (Arifuzzaman et al., 2022). However, the key mechanisms for how these changes in diet and gut microbiome tune lung immune tone largely remain to be fully defined as well as the specific effects of different fiber-sources and key bacterial species. Therefore, the identification of pivotal metabolically-active gut bacteria and bioactive metabolites could potentially lead to a better comprehension of inter-individual immune variability, as well as the exploration of potential therapeutic approaches and applications.

Lung immune tone refers to the level of immune priming or baseline immune set point present within the lung, which in turn dictates the degree of inflammatory responses following injury. Numerous lung diseases have implicated two NLRP3-dependent key cytokines, IL-1β and IL-18, as key regulators of downstream inflammatory responses and pathology. These include sterile injuries (Couillin et al., 2009; Gasse et al., 2007; Liu et al., 2021; Tian et al., 2017), infections (pneumonia, viruses, parasites, etc. refs), asthma responses (Abdel Fattah et al., 2015; Jordan et al., 2001; Trompette et al., 2014), etc. The secretion of IL-1β and IL-18 require a two-step process(Barnett et al., 2023), and we have earlier shown that the levels of these two cytokines’ mRNA can be used as a surrogate for measurement of lung immune tone(Liu et al., 2021). We also previously showed that gut-microbiome derived SCFA metabolites travel from the gut to the lung and can regulate the inflammatory output of alveolar macrophages (Liu et al., 2021; Tian et al., 2019).

Here, we used pectin-based high-fiber and fiber-free diets to study their effects on lung immune tone in sterile lung injury responses. We conducted targeted SCFA analyses in conjunction with inflammasome-focused lung transcriptomics and proteomics, observing both augmentation and suppression of IL-1β and IL-18 pathways by fiber-diet driven gut bacterial taxa and specific SCFA metabolites. The observed *in vivo* changes in IL-1β and IL-18 lung immune tone could be reproduced by solely stimulating alveolar macrophages with the SCFA propionate alone, thereby revealing key inflammatory and metabolic reprogramming mechanisms as a result of SCFA exposure. Overall, we demonstrate the importance of dietary fiber in influencing the gut microbiome composition and associated alterations in lung immune tone. SCFAs such as propionate play an important role in mediating the gut-lung immune axis.

## MATERIALS AND METHODS

### Animal care

All studies were approved by the institutional animal care and use committee at University of California, San Francisco (IACUC protocol# AN197325-01). All mice were purchased from the Jackson Laboratory (Bar Harbor, ME) or bred in the animal facility at University of California, San Francisco. Wild-type (WT) C57BL/6, and germ-free (GF aka gnotobiotic) mice were used in this study (10-15 week old GF or WT C57BL/6J adult male mice). Commercially purchased mice were allowed to acclimatize to their new housing for at least 1 week before any experiments were conducted on them.

Fiber diet intervention (schematized in **Supplemental Figure S2**): Mice received standard animal facility chow for at least 1 week after arriving at UCSF (**Supplemental Figure S11B**) after which for experiments with fiber diet intervention, mice either received 2 weeks of 0% fiber (**Supplemental Figure S11C**) or 2 weeks of 35% pectin fiber (**Supplemental Figure S11D**). Mouse stool quality and intestinal morphology was observed after IR surgery as well as weight changes over the course of the dietary intervention (**Supplemental Figure S11A**). A cross-over group was also included in which mice received 1 week of 0% fiber followed by 1 week of 35% pectin fiber prior to lung IR surgery (schematized in **Supplemental Figure S5A**).

GF mice (C57BL/6 background) were obtained from the UCSF Gnotobiotic Facility. In accordance with Biosafety Level 2 guidelines from the Centers for Disease Control and Prevention, GF mice were maintained in sterile gnotobiotic isolators under a strict 12h light cycle and were fed an autoclaved standard diet.

Left lung ischemia reperfusion (IR) surgery: A murine model of unilateral left pulmonary artery occlusion was used, as we have described previously (Prakash et al., 2012). Briefly, anesthetized mice (using IP tribromoethanol (Avertin®); Sigma-Aldrich) were orally intubated, given buprenorphine (IP; Harry Schein, Melville, NY), and placed on a rodent ventilator, using tidal volumes of 225uL (7.5cc/kg), and a respiratory rate of 180 breaths/min (assuming an average mouse weight of 30g). A left thoracotomy via the interspace between the 2nd and 3rd ribs was performed and the left pulmonary artery (PA) was identified and ligated using a slip knot suture with 7-0 or 8-0 prolene monofilament suture. The end of the suture externalized through a narrow bore (27g) needle to the anterior chest wall. Prior to closure of the thorax, the left lung was reinflated with positive end pressure. Local anesthetic (3-4 drops of 0.25% bupivacaine) was applied topically prior to skin closure. The total period of mechanical ventilation and surgery was approximately 20-25min. After skin closure, mice were extubated and allowed to recover from anesthesia. After 60min of ischemia, the ligature on the pulmonary artery was released and left lung reperfusion started. At the experimental end point times, mice were euthanized, and the blood, feces, and lungs were collected.

Blood was collected from anesthetized mice via cardiac and portal vein puncture using a heparinized syringe, centrifuged (14,000g, 5min) and the plasma separated, flash frozen in liquid nitrogen and stored at -80°C. Lower portions of the left lungs were excised and divided into two parts: one was placed in either Trizol® (Life Technologies, Carlsbad, CA) for RNA preparation and the second part was frozen at -80°C for homogenization for ELISA. Levels of cytokines and chemokines were quantified in plasma or homogenized lung tissue by ELISA or in lung tissue by qPCR.

All mice received equivalent durations of mechanical ventilation (20-25min), and were left spontaneously breathing during their recovery from anesthesia and the remainder of the ischemia period and subsequent reperfusion or equivalent periods in the sham mice.

Mice that did not survive the surgery or the reperfusion period due to technical complications in the surgical procedure (predominantly, left bronchus or left PA injury) were excluded from the study. Pre- and post-operative care was conducted consistent with ARRIVE guidelines.

### Reagents and cell lines

Propionate SCFA (catalog number P1880) and lipopolysaccharide (LPS, catalog number L4391, from Escherichia coli O111:B4) were purchased from Sigma-Aldrich (St. Louis, MO). Seahorse XF calibrant solution (part number 100840-000), Seahorse XF DMEM medium, pH7.4 (catalog number 103575-100), Seahorse XF RPMI medium, pH 7.4 (catalog number 103576-100), Seahorse XF1.0 M glucose solution (catalog number 103577-100), Seahorse XF 100mM pyruvate solution (catalog number 103578-100), Seahorse XF 200mM glutamine solution(part number 103579-100), and Seahorse XF Cell Mito Stress Test Kit (catalog number 103015-100, contains 1 each of oligomycin, FCCP, and rotenone/antimycin A.) were all purchased from Agilent Technologies, Inc. (Santa Clara, CA).

The following cell lines were used in this study: MLE 12 (WT FVB/N murine lung epithelial AT2 cell, catalog number CRL-2110), MH-S (wild-type BALB/c alveolar macrophage, catalog number CRL-2019) – all purchased from ATCC (Manassas, VA); all cells were incubated at 37°C under humidified 5% CO_2_ and then treated with 10-200ng/mL LPS for 24h (or other time points) as indicated.

### Short-chain fatty acid quantification

SCFA in mouse lung samples were quantified at the Metabolomics Core facility at University of Michigan. They performed sample extraction using aqueous extraction solvent containing 3% 1 M HCl (v/v) and isotope-labeled internal standards (d7-buytric acid and d11-hexanoic acid). Samples were then homogenized and centrifuged. Supernatants were transferred to new Eppendorf tubes for extraction by diethyl ether. After phase separation, the upper layer was transferred to an autosampler vial for gas chromatography-mass spectrometry (GC-MS) analysis. GC (Agilent 6890, Wilmington, DE) separation was performed using a ZB-Wax plus column, 0.25 μm × 0.25 mm × 30 m (Phenomenex, Torrance, CA). A single quadrupole mass spectrometer (Agilent, 5973 inert MSD) was used to identify and quantify SCFA using Agilent Masshunter software, version B.06 [14]. Absolute quantities of SCFA were normalized to sample mass.

### Seahorse^TM^ Extracellular Flux Metabolism Analysis

MH-S cells were cultured in RPMI-1640 growth medium (Gibco, 72400047) supplemented with 10% FBS (Gibco, 10438026) and 1% Penicillin/streptomycin (Gibco, 15140122) at 37°C in a 5% CO2 atmosphere. To examine the metabolic response in MH-S alveolar macrophage (AM) cell lines exposed to LPS, LTA, and SCFAs overnight before metabolic analysis, we utilized the Agilent Seahorse XFe24 analyzer platform (Agilent Technologies, Inc.) following the manufacturer’s instructions. The Seahorse XF Cell Mito Stress Test was used to measure mitochondrial function by analyzing the oxygen consumption rate (OCR) of cells. This plate-based live cell assay enables real-time monitoring of OCR.

The assay consists of two steps: the day prior to the assay and the day of the assay. On the day prior to the assay, cultured MH-S cells were seeded into the Seahorse XFe24 assay plate at an optimized density of 40,000 cells per well. After the cells fully attached to the plate bottom, they were treated with LPS (10 ng/mL), LTA (100 μg/mL), and SCFA propionate (C3, 0.1 mM or 5 mM) contained in RPMI 1640 full culture medium, and were left overnight. A sensor cartridge was hydrated with XF calibrant at 37°C in a non-CO2 incubator overnight. The experiment was designed in the Ware software on the day prior to the assay as well.

On the day of the assay, an assay medium was prepared by supplementing XF RPMI medium with 1 mM pyruvate, 2 mM glutamine, and 10 mM glucose, ensuring a pH of 7.4, it’s crucial to maintain this pH. All compounds of the Mito Stress Test (Oligomycin, FCCP, and Rot/AA) were resuspended and diluted into appropriate working concentrations for the assay, optimized concentrations of 1.5μM for Oligomycin, 1μM for FCCP, and 0.5μM for Rot/AA were applied. The assay plate containing MH-S cells needed to be carefully washed with fresh and warm assay medium, and then placed in a non-CO2 incubator for 1 hour before assay. All ports of the sensor cartridge were properly loaded with the three compounds, and the cartridge was placed into the analyzer platform for calibration. Before beginning the assay, the Seahorse XF assay plate was removed from the 37°C non-CO2 incubator, the cells were examined under a microscope to confirm confluence, and then the cell assay plate was loaded to initiate the assay.

### Reverse transcription quantitative real-time polymerase chain reaction (RT-qPCR)

TaqMan-specific inventoried gene primers for glyceraldehyde 3-phosphate dehydrogenase (GAPDH), beta actin (Actb), ribosomal protein L19 (Rpl19), interleukin (IL)-6, IL-1β, IL-10, IL-18, NOD-like receptor family, pyrin domain containing (NLRP) 3, chemokine (C-X-C motif) ligand (CXCL) 1, CXCL2 (Life Technologies, Carlsbad, CA) were used to measure the mRNA levels of these human or mouse genes in cells or lung tissue.

Lung tissue was homogenized (Tissue-Tearor - Biospec Products, Bartlesville, OK) and total RNA isolated using Trizol^®^ (Invitrogen – Thermo Fisher Scientific, Waltham MA) and RNAeasy Mini Kit (Qiagen) to purify RNA. We used the High-Capacity RNA-to-cDNA reverse transcription Kit (Life Technologies) using 1 μg messenger RNA per reaction. Quantitative real-time polymerase chain reaction was performed using TaqMan fast advance master mix and TaqMan gene expression assay (reagents) and QuantStudio^TM^ 6 and 7 Flex Real-time PCR systems (device). Run method: Polymerase chain reaction activation at 95°C for 20s was followed by 40 cycles of 1s at 95°C and 20s at 60°C.

The average threshold cycle count (Ct) value of 2–3 technical replicates was used in all calculations. The average Ct values of the internal controls (GAPDH, beta actin, L19) was used to calculate ΔCt values for the array samples. Data analysis was performed using the 2^-ΔΔCt^ method, data presented as relative quantification (RQ), and the data were corrected for statistical analysis using log transformation, mean centering, and autoscalingClick or tap here to enter text.(Livak & Schmittgen, 2001; Schmittgen & Livak, 2008)Click or tap here to enter text..

### Lung immune tone and Lung inflammatory marker measurements

Based on our prior published work, we define lung immune tone by the levels of IL-1β and IL-18 mRNA as quantified by qPCR(Liu et al., 2021). After lung IR injury, early markers of inflammation are used as indicators for severity of injury response; these markers include IL-6, CXCL-1, CXCL-2 (schematized in **Figure S1**) as we have previously reported(Prakash et al., 2015; Tian et al., 2017, 2019).

### Sandwich enzyme-linked immunosorbant assay (ELISA)

Concentrations of cytokines and chemokines in cell culture supernatant, plasma and lung homogenate were determined using the mouse DuoSet ELISA kit (R&D Systems, Minneapolis, MN). All assays were performed according to the manufacturer’s supplied protocol. Standard curves were generated and used to determine the concentrations of cytokines and chemokines in the samples. Results are presented as Mean ± SD.

### Correlation Analyses (16S sequencing data, metabolomic data, immune/inflammatory markers)

The size of the sequenced pair-end libraries ranges from 5,976bp to 285,864bp, representing a total of over 15 million 151bp reads. Reads were processed through Qiime2 (version 2020.8.0)(Bolyen et al., 2019):low-quality reads and sequencing adaptors were removed using Cutadapt(Martin, n.d.), and sequencing errors were corrected with Dada2(Callahan et al., 2016) using custom parameters (--p-trunc-len-f 150 --p-trunc-len-r 140). Taxonomic assignation of resulted ASVs was done using SILVA trained database (version 138-99)(Quast et al., 2013) based on scikit-learn’s naïve Bayes algorithm(Pedregosa FABIANPEDREGOSA et al., 2011). For the subsequent analyses (except for alpha-diversity calculations), the abundance of each taxon present in a sample was normalized using the relative method to allow sample-to-sample comparison. Results were deep analyzed with the Phyloseq package (version 1.34.0)Click or tap here to enter text.(McMurdie & Holmes, 2013)Click or tap here to enter text. as for the analysis of taxonomic and alpha diversity. Statistical analyses were performed using rstatix (version 0.7)(Kassambara, 2021) and figures were plotted using the ggplot2 package (version 3.3.5)(Wickham, 2016). Principal coordinate analyses (PCoA) were carried out with Vegan package (version 2.5-7)(Dixon, 2003) on the Bray-Curtis dissimilarity matrices constructed from the relative abundance of ASVs. Communities that emerged were verified using a PERMANOVA test and the confidence interval were plotted at 95% and 97% confidence limits, using the standard deviation method. Spearman correlation analyses were conducted to establish associations between metabolites and microbiota species using the R package energy (version 1.7-8).

### Statistical Analysis

Data in the figures are expressed as mean +/- SD. Data from *in vivo* studies comparing two conditions were analyzed using 2-tailed nonparametric Mann–Whitney analyses. Aseesa’s Stars (www.aseesa.com) analysis tool was used to analyze and generate principal analysis donut and scatter plots, correlation, volcano plots, and heat maps. Data from *in vitro* studies comparing two conditions were analyzed using standard Student’s t-test with equal SD to generate *P* values. GraphPad Prism was used for statistical analyses (GraphPad Software, La Jolla, CA). P values < 0.05 were considered significant. P values are represented as follows in the figures: *< 0.05; **< 0.01; ***< 0.001; ****< 0.0001. Experiments were repeated 2 or more times, as indicated in the figure legends. One cage was initially excluded from beta-diversity analysis, Mantel and Procrustes analyses due to the presence of *E. coli/Shigella* in the fecal microbiome composition but included in all other analyses (as described in the text below).

## RESULTS

### High Fiber (HiFib) diet results in reduced sterile lung injury responses

Having previously reported that direct lung administration of SCFA led to reduced lung immune responses and altering the gut microbiome leading to an enrichment in SCFA-producing *lachnospiraceae* species resulted in reduced sterile lung injury (Tian et al., 2019), we asked the question whether creating a dietary environment to enrich for bacteria that ferment fiber and thus produce SCFAs might result in a similar reduction in lung injury responses. Additionally, we sought to better understand the link between the gut microbiome and the IL-1β priming (aka lung immune tone) upstream of NLRP3-dependent lung IR injury responses. We hypothesized that gut bacteria that produced SCFAs might contribute to the regulation of IL-1β mRNA levels in lung tissue and thereby control lung immune tone to early inflammatory injury (as Signal 1, schematized in **Supplemental Figure S1**).

We tested this hypothesis by providing fiber-rich diets containing 35% pectin, as based on prior published work (Lucas et al., 2018; Trompette et al., 2014), to mice for 2 weeks. To elucidate the effects of fiber on the microbiome and lung injury responses, we chose to use a fiber-free diet as our control (**Supplemental Figure S2**). After a 2 week dietary intervention, we performed lung IR injury (1h left lung ischemia, 1h reperfusion) and subsequently examined immediate early lung injury responses after 1h reperfusion (**Figure 1A**). This early time point post injury has been validated from our previous work to investigate first wave cytokine and chemokine release prior to the histopathological signs of lung injury, which include edema, hyalin casts, and neutrophil infiltration, occurring 3h following reperfusion (Prakash et al., 2012). As shown in **Figure 1B**, high-fiber diet intervention prior to IR injury led to a marked reduction of systemic IL-6 secretion as well as tissue IL-6 levels. Furthermore, there was a significant reduction in other inflammatory markers including in the neutrophil recruiting chemokines, CXCL-1 and CXCL-2.

**Figure 1.**
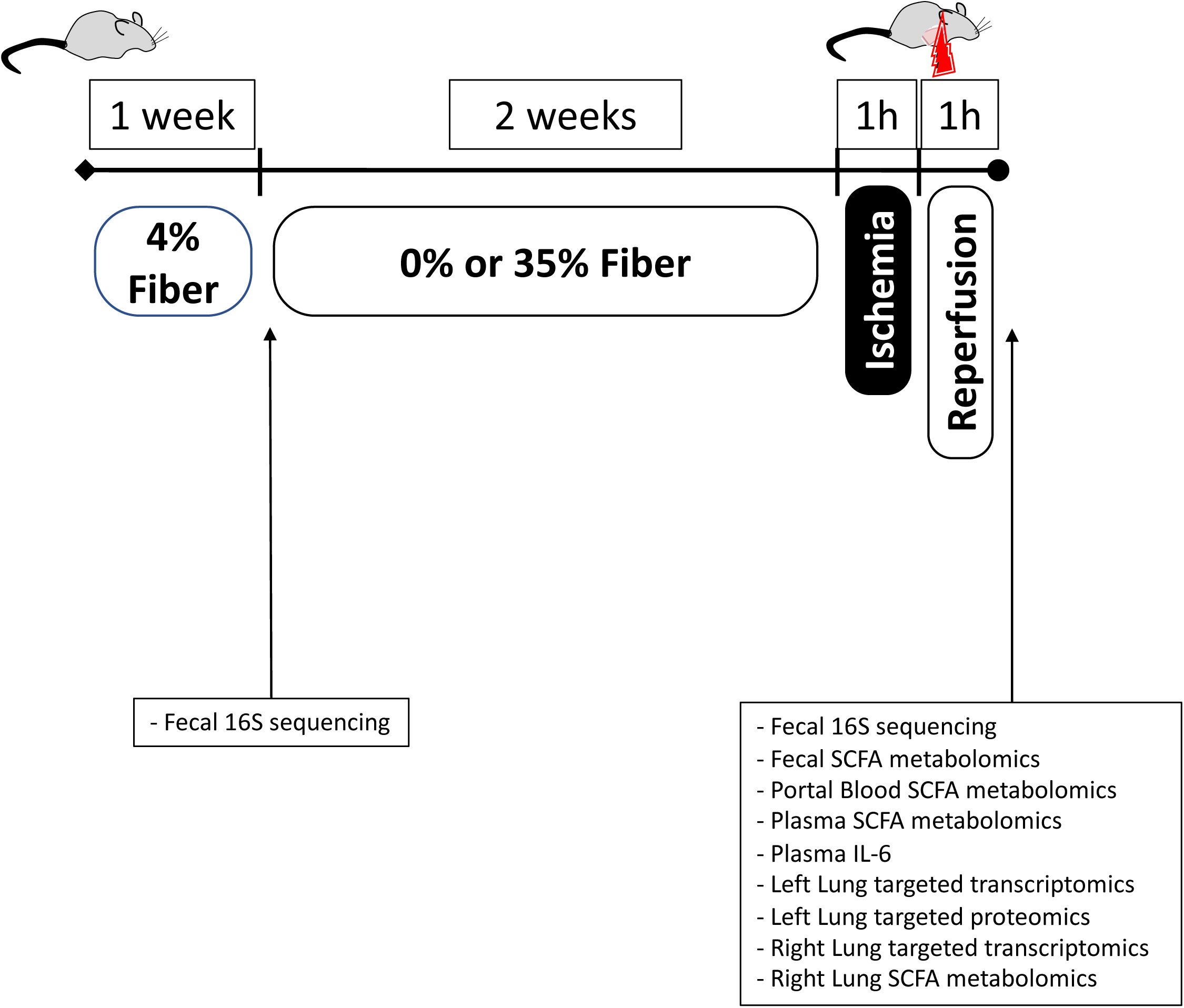
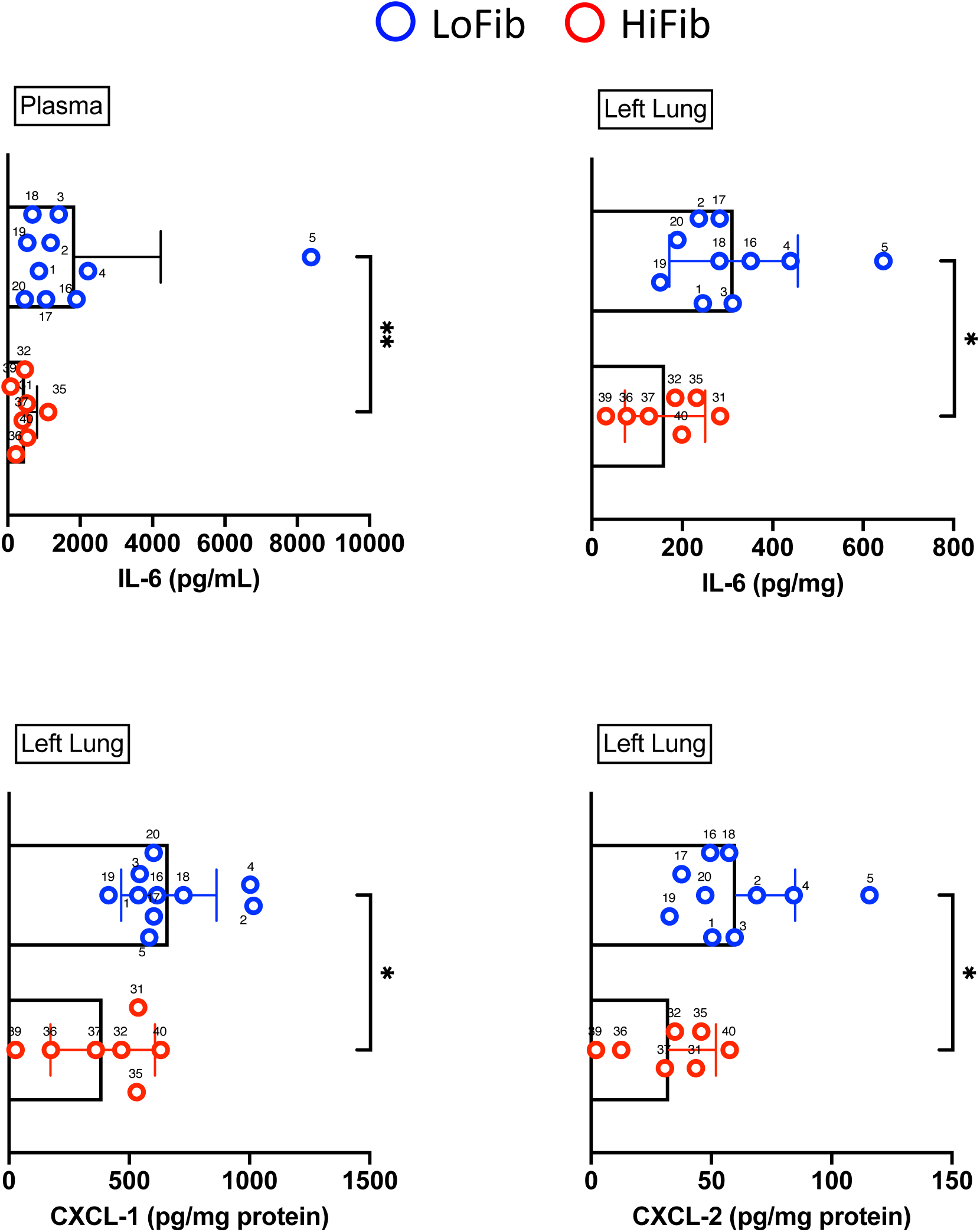
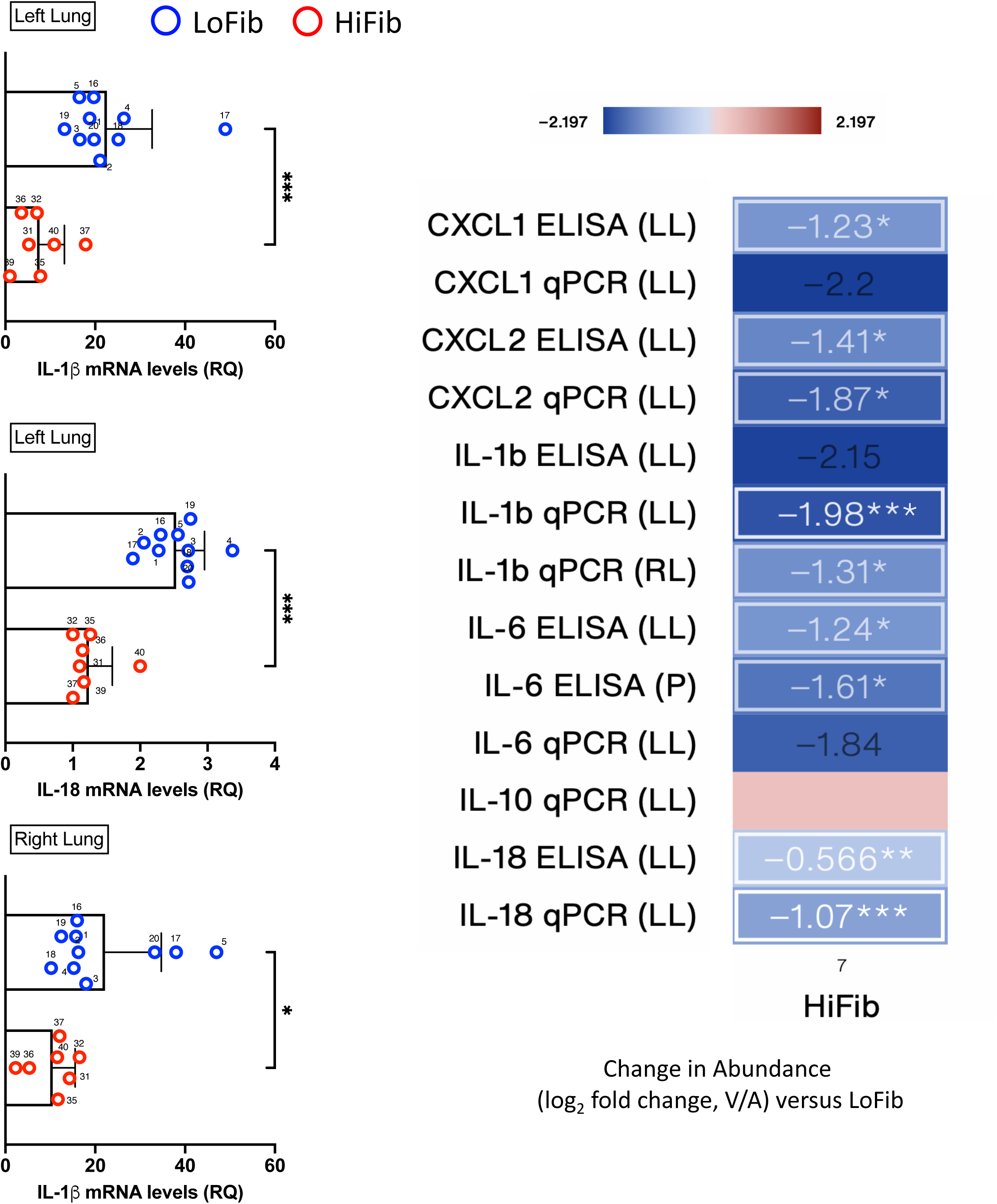
High pectin fiber diet reduces lung IR injury related inflammatory responses and lowers lung immune tone. (A) Experimental outline for low fiber (0% fiber/LoFib) and high fiber (35% fiber/HiFib) dietary intervention, followed by lung ischemia reperfusion (IR) injury two weeks later. (B) Post-lung IR injury inflammatory markers: plasma IL-6 protein levels as well as left lung tissue IL-6, CXCL-1, and CXCL-2 expression in LoFib and HiFib groups after lung IR. (C) Markers of lung immune tone: relative mRNA (RQ) levels of IL-1β and IL-18 from injured left lung tissue (left top and middle), IL-1β from uninjured right lung tissue (left bottom). Heatmaps showing log2 fold changes in various chemokine/cytokines levels in left lung (LL), right lung (RL) and plasma (P) as measured with ELISA and qPCR. Significantly changed measures are in white boxes with ***, ** and * denoting p < 0.001, 0.01 and 0.05, respectively by Welch’s t-test. log2 fold change versus low fiber diet (LoFib) is shown. Color scale: red denotes increased levels and blue denotes decreased levels versus control group.

### HiFib diet reduces lung immune tone

The importance of IL-1β and NLRP3 inflammasome in lung IR injury responses has been reported(Tian et al., 2019). We observed significantly lower levels of IL-1β and IL-18 mRNA in the left lungs of our HiFib group indicating reduced lung immune tone (**Figure 1C, left top and middle**). Interestingly, even in the right lung that was not directly subjected to injury, there was a similar reduction in IL-1β mRNA levels (**Figure 1C, left bottom**). Overall, compared to the low fiber control group, we observed that high fiber diet resulted in reduced lung immune tone and inflammatory marker production and release in the injured left lung and plasma, as well as in the uninjured right lung (summarized in **Figure 1C, right**).

### HiFib vs. LoFib diet changes gut microbiome taxa, SCFA production and transition along the gut-lung axis

Next, we examined the impact of the dietary fiber interventions on gut microbiota. Following the 2-week diet change, we were able to observe significant alternations in the 16S composition of the gut microbiome with HiFib diet with an expansion of *Bacteriodetes* phyla and a reduction of the *Firmicutes:Bacterioides* ratio (**Figure 2A**) and enrichment for specific taxa (**Supplemental Figure S3**). Using beta-diversity analysis, we observed clear clustering of gut microbiota from all mice at baseline prior to diet change and distinct separation of the HiFib group microbiota away from the LoFib group (**Figure 2B**). Enriched and depleted taxa in LoFib and HiFib groups compared to baseline microbiome composition were also observed with specific and significant enrichment of *Bacteriodes* and *Parasutterella* in the HiFib group (**Supplemental Figure S4A**). We also observed a reduction in alpha diversity with HiFib diet (**Supplemental Figure S4B**). This reduction in richness and diversity has also been observed in other fiber diet intervention studies (Trompette et al., 2018) and very likely reflecting the strong effect of pectin fiber in driving the expansion of pectin-fermenting *Bacterioides* genera. Targeted SCFA metabolomics of the feces, portal blood, and circulating plasma reflected specific changes in SCFAs as well as medium and branch-chain fatty acids (MCFA, BCFA) with acetate (C2) and propionate (C3) significantly enriched in portal blood and plasma in the HiFib group in contrast to the LoFib group (**Figure 2C**). Interestingly, the LoFib group exhibited elevated levels of MCFA and BCFA (heptanoate, caproate, valerate) in their feces and specific enrichment of these metabolites in lung tissue (**Figure 2C**).

**Figure 2.**
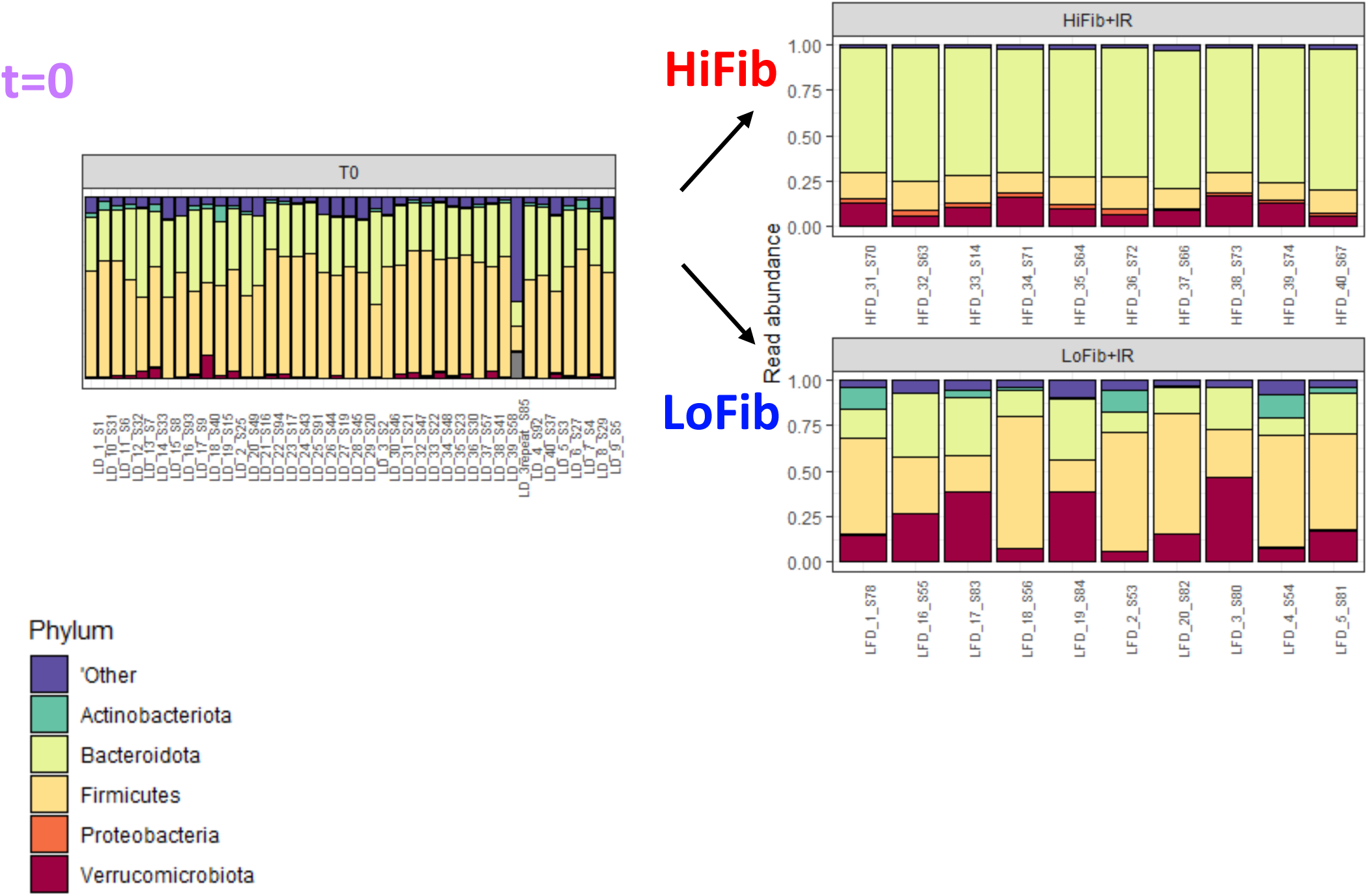
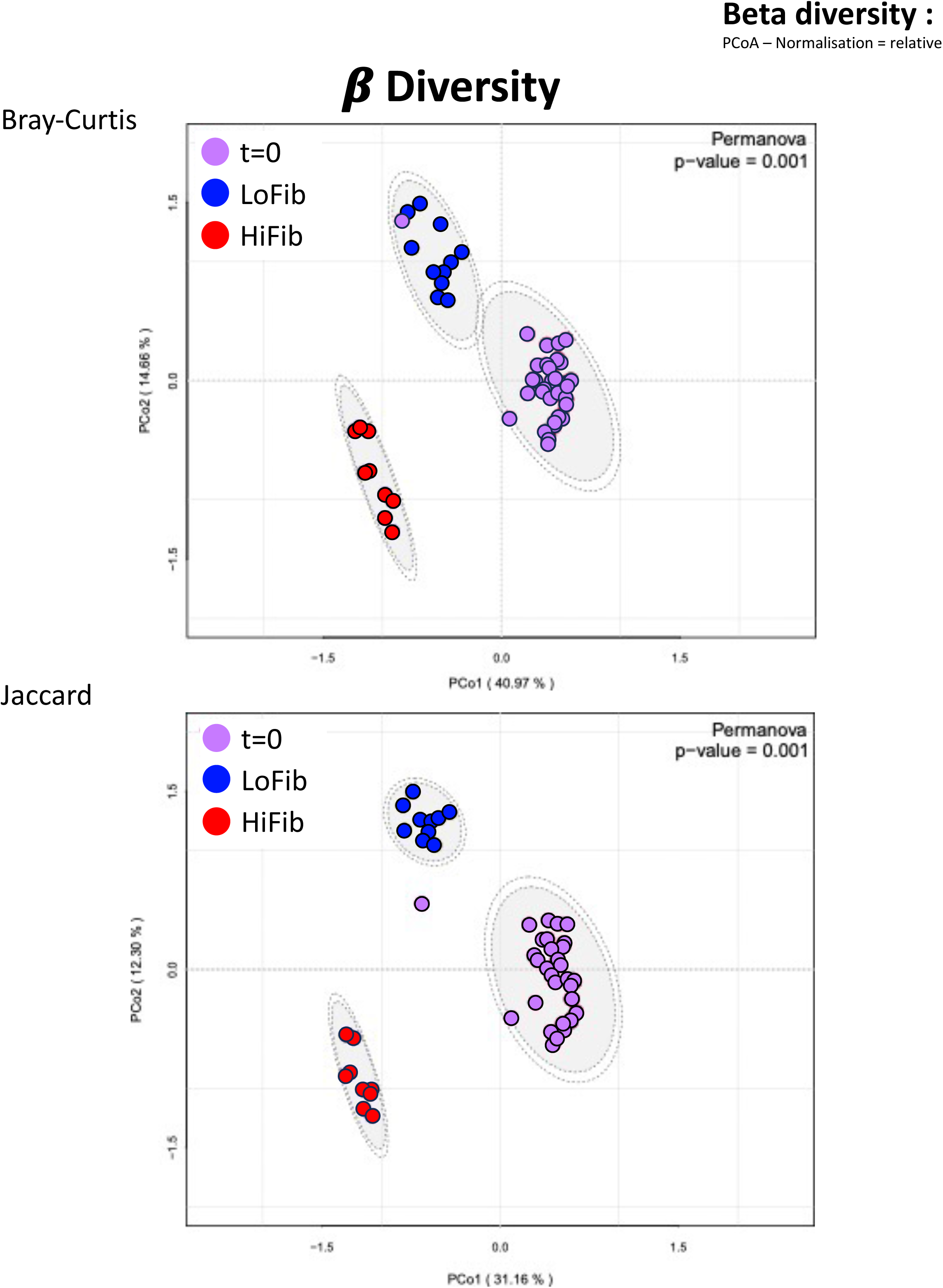
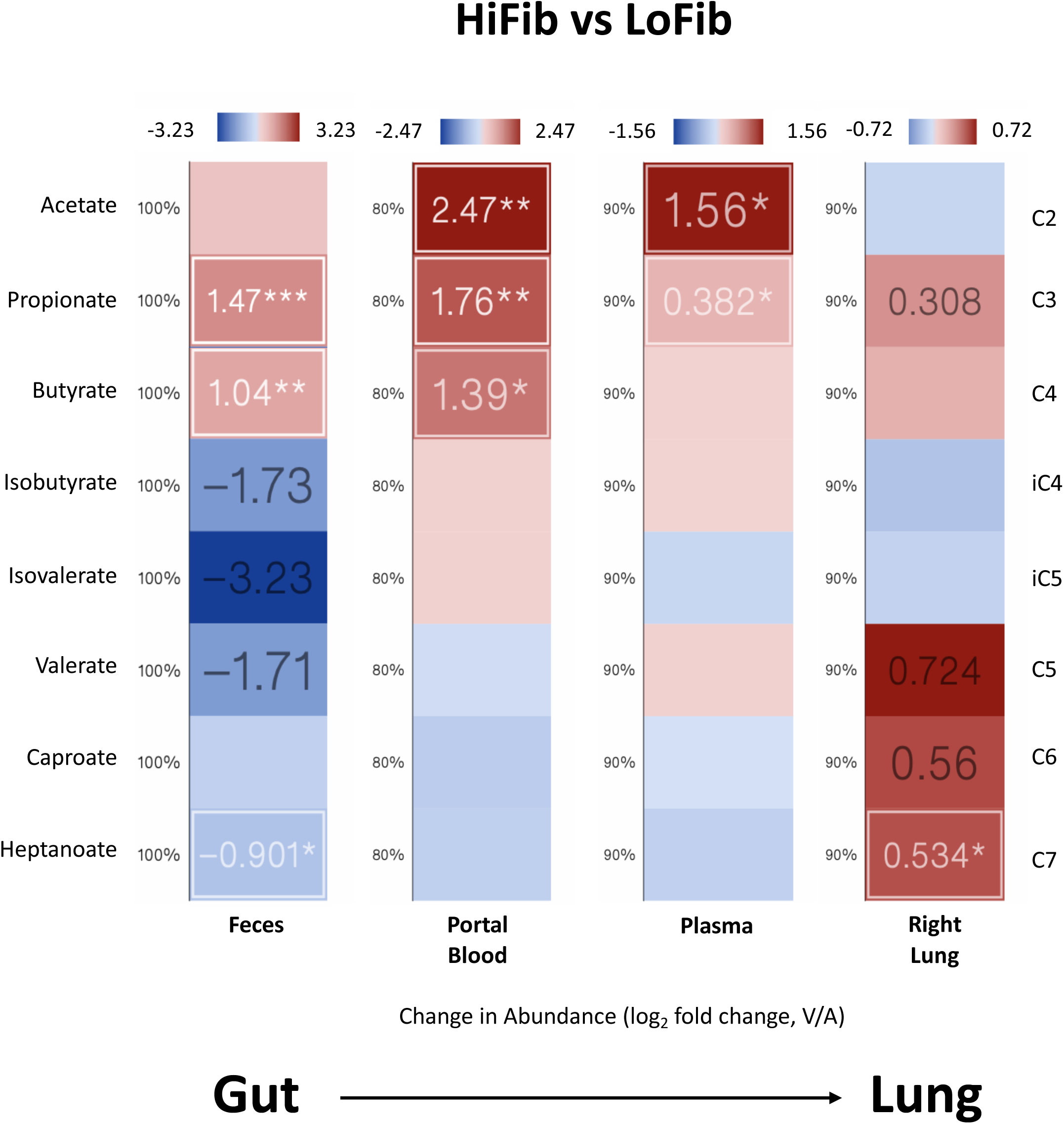
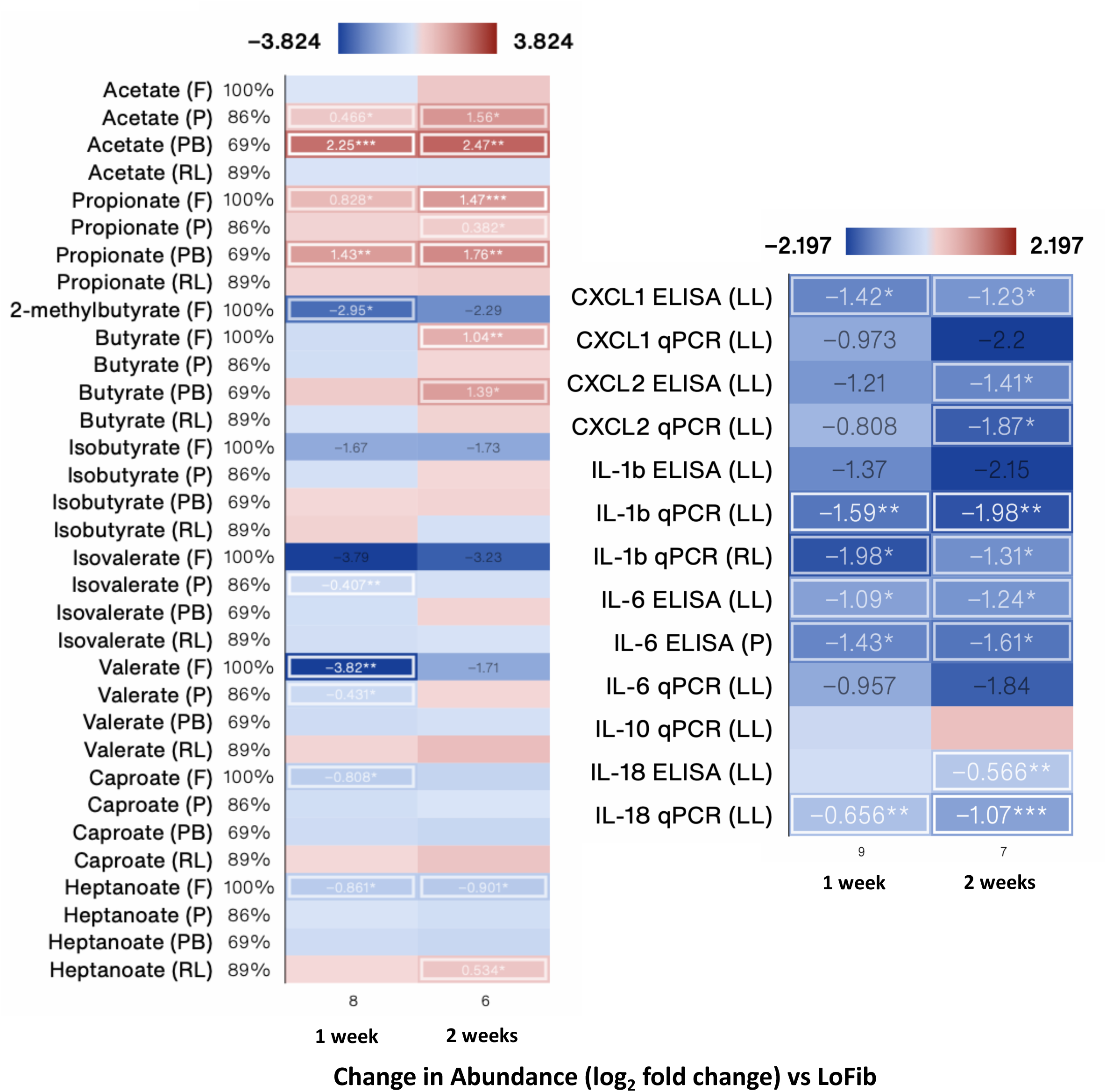
Fiber diet results in changes in gut microbiome and gut metabolome. (A) Phylum level changes in gut microbiome comparing mice on normal (4% fiber) chow (t=0, prior to dietary intervention), LoFib and HiFib groups (after 2 weeks of dietary intervention). (B) Beta diversity clustering of mouse groups (Bray-curtis top, Jaccard bottom). (C) Heatmaps showing log2 fold changes in SCFA levels in feces, portal blood, plasma, and right lung tissue as measured with mass spectrometry. (D) Heatmaps showing log2 fold changes in SCFA levels (left) and immune tone/inflammatory markers (right) between 1-week and 2-week HiFib groups. F=feces, PB=portal blood, P=plasma, LL=left lung, RL=right lung. Significantly changed measures are in white boxes with ***, ** and * denoting p < 0.001, 0.01 and 0.05, respectively by Welch’s t-test. log2 fold change versus low fiber diet (LoFib) is shown. Color scale: red denotes increased levels and blue denotes decreased levels versus control group.

### 1-week vs 2-week HiFib diet have similar effects on lung injury responses, lung immune tone, gut microbiota composition, and SCFA production

To test whether one week of HiFib diet was sufficient to alter lung immune tone and if the effect of high fiber could dominate over low fiber or vice versa, we subjected mice to 1 week of LoFib diet, followed by 1 week of HiFib diet and performed the same analyses as described above (schematically shown in **Supplemental Figure S5A**). We found that 1 week of HiFib diet even after prior LoFib diet exposure was able to generate very similar SCFA profiles compared to 2 weeks of HiFib diet (**Figure 2D, left and Supplemental Figure S6**) as well as result in similarly reduced lung injury responses and lung immune tone (**Figure 2D, right and Supplemental Figure S6**). Furthermore, this 1-week HiFib group’s gut microbiota was more similar to the 2-week HiFib group in terms of alpha and beta diversity, with significant overlap in enriched taxa (data not shown). Interestingly, one cage from this group (5 mice) did not show an enrichment for the same taxa as the rest of the HiFib exposed mice, and instead by 16S sequencing appeared to have a *E. coli/Shigella* enrichment (data not shown). We included this group for later analyses since we were interested in uncovering drivers of both elevated and reduced lung immune tone.

### Specific bacterial taxa, associated metabolic pathways, and metabolites strongly correlate with lung immune tone regulation

Further analysis of our dataset consisting of multiple measurements (16S bacterial sequences, metabolomics, transcriptomics, proteomics) from individual mice provided the opportunity to uncover key correlations between measured factors. These analyses could provide the basis for generating hypotheses regarding the mechanistic control of the biological variation observed in lung immune tone. Principal component analysis (PCA) was performed on the 2-week HiFib data set focusing on metabolites (from all sample sources), protein and transcript data (from lung and plasma) revealed five modules or principal components that explained 90% of the variation in the combined 2-week HiFib data set (**Figure 3A**). Modules combining SCFAs and inflammatory markers were observed suggesting potential relationships between these factors in co-regulating the gut-lung axis. In the 2-week HiFib group (compared to the LoFib group), acetate and propionate were the most elevated in the portal blood, feces and plasma, while lung IL-18 and IL-1β mRNA were the most significantly diminished (**Figure 3B**). Performing the same PCA and volcano plot analyses on the 1-week HiFib group revealed similar modules (7 modules explaining 90+% of the variation in the 1-week HiFib dataset) (**Supplemental Figure 7A**) and metabolite/transcript enrichment/depletion along with a more pronounced difference in the levels of fecal valerate, isovalerate and 2-methyl butyrate between 1-week HiFib and the LoFib control group (**Supplemental Figure 7B**). Comparing the 1-week and 2-week HiFib diet groups, we found that 11 measured factors changed significantly in the same direction (including key SCFAs and lung immune tone and lung injury markers of interest), while 7 were significant only for the 2-week HiFib group and 5 only for the 1-week HiFib group (**Figure 3C**). Notably, none changed in opposite directions between these two groups. To further elucidate the interrelationships among these datasets (now including the 16S bacterial taxa data) we employed Mantel and Procrustes tests (**Figure 3D, top and middle tables**). These analyses revealed significant correlations between the gut microbiome 16S genomic dataset and the SCFA metabolomic datasets. These were expected given the fact that gut microbiota are the predominant source of SCFA production in the body. However, we also noted highly significant correlations between the 16S dataset and the NLRP3-targeted lung transcriptomic dataset as well as significant correlation with lung tissue/plasma inflammatory proteins. Visual representation of the Procrustes analysis highlighted close relationships between 16S data and protein, transcript and metabolite data, which were particularly evident in the HiFib groups (shorter line distances on the left of each plot in **Figure 3D, bottom**).

**Figure 3.**
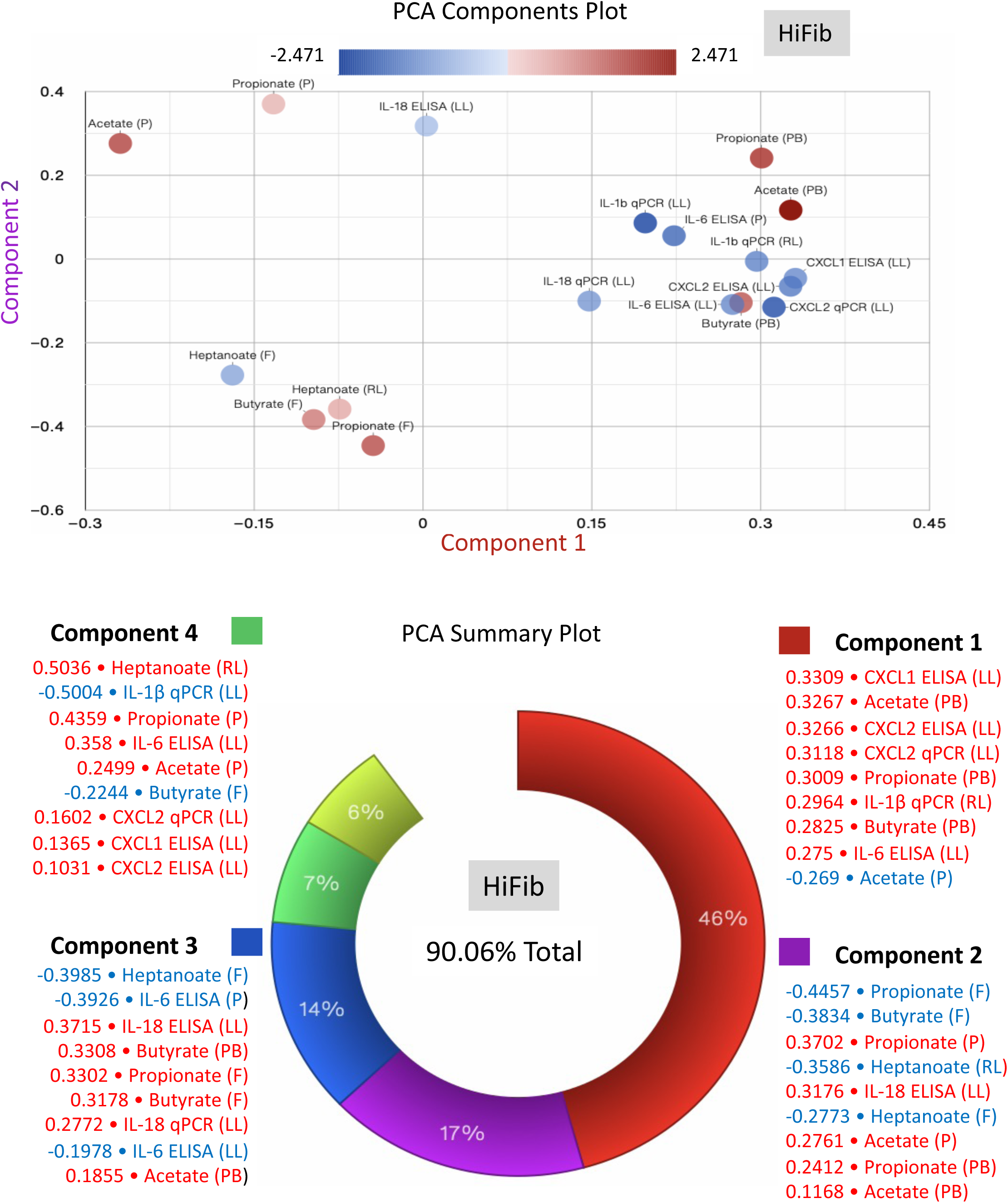
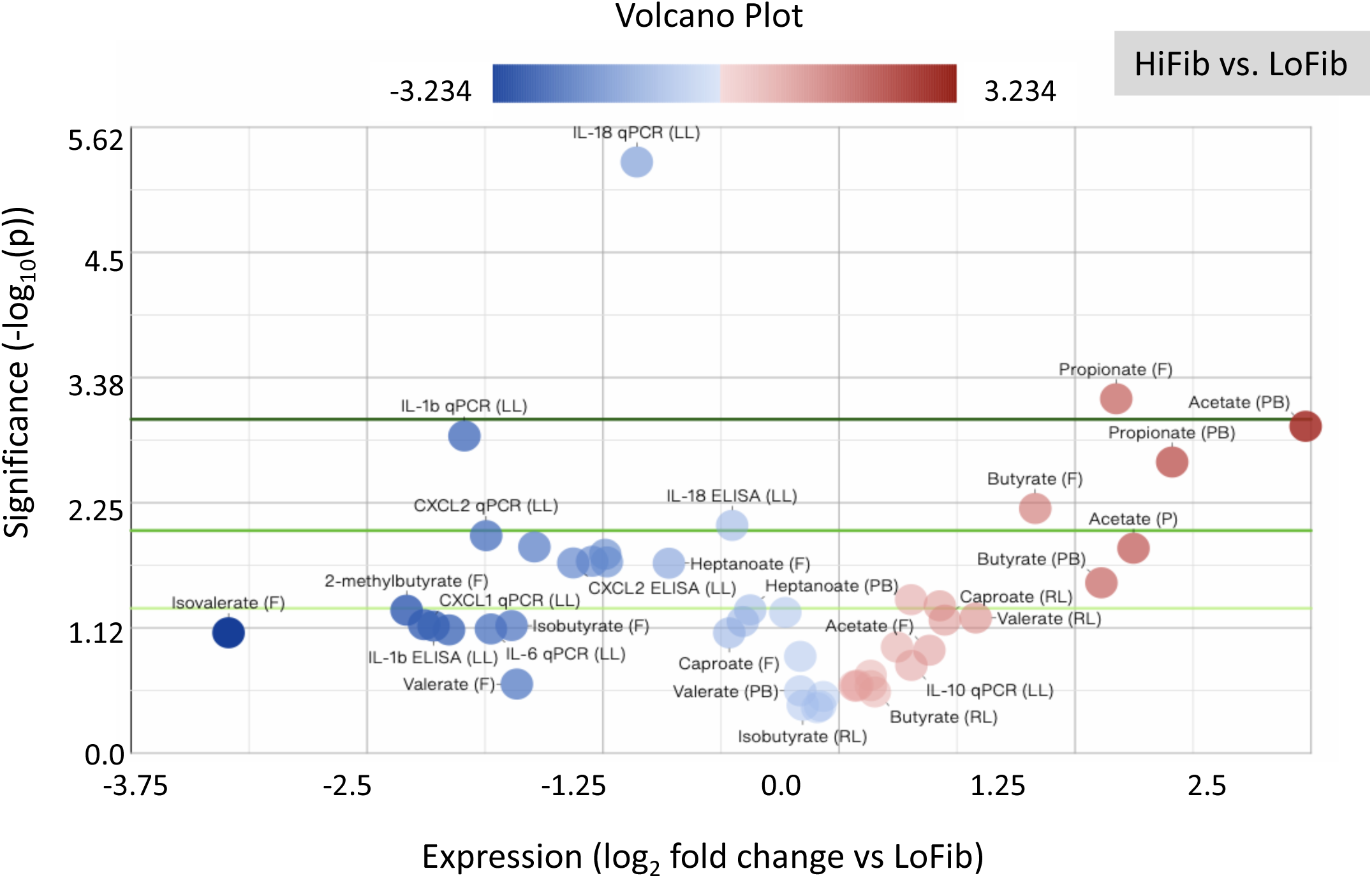
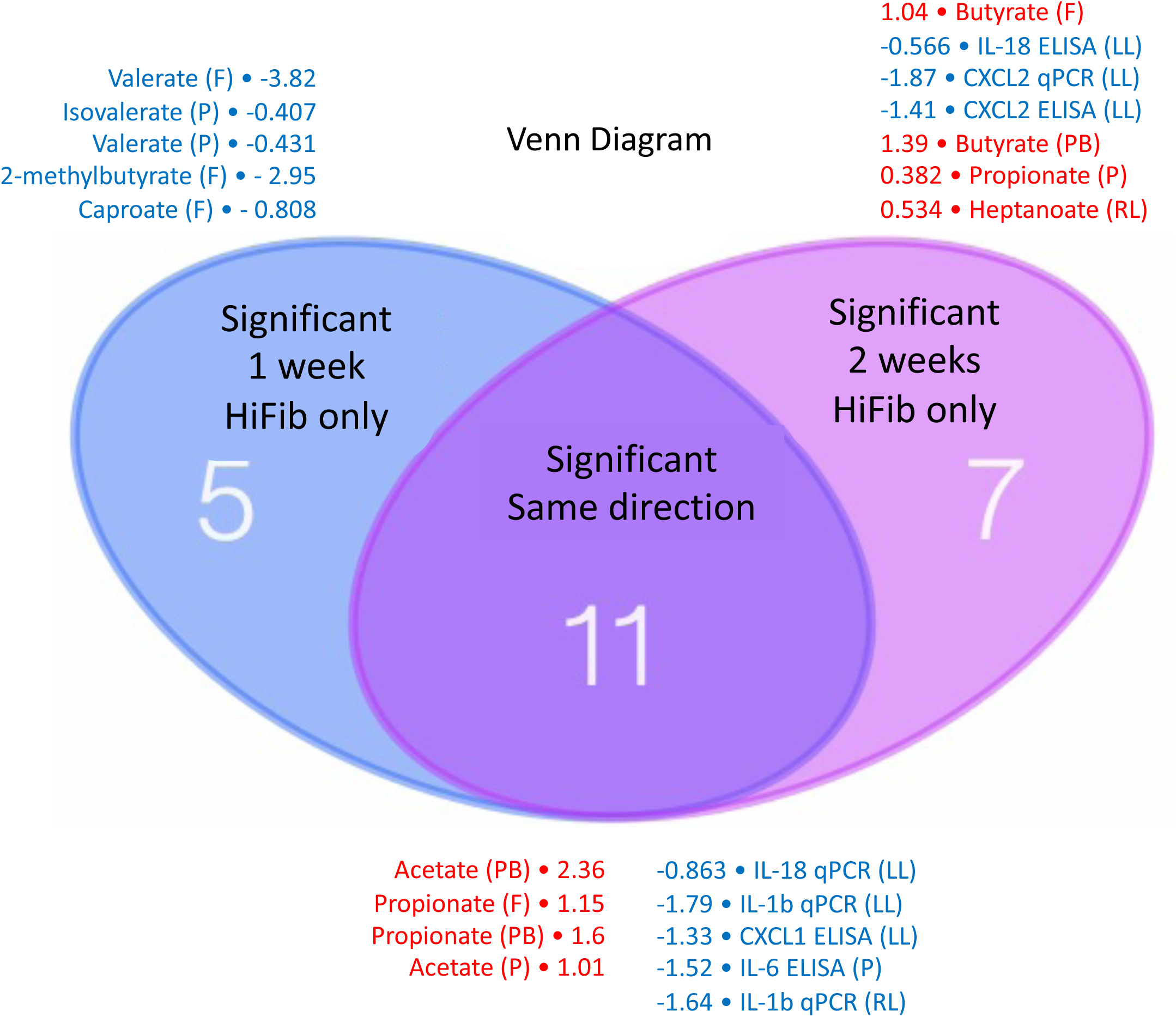
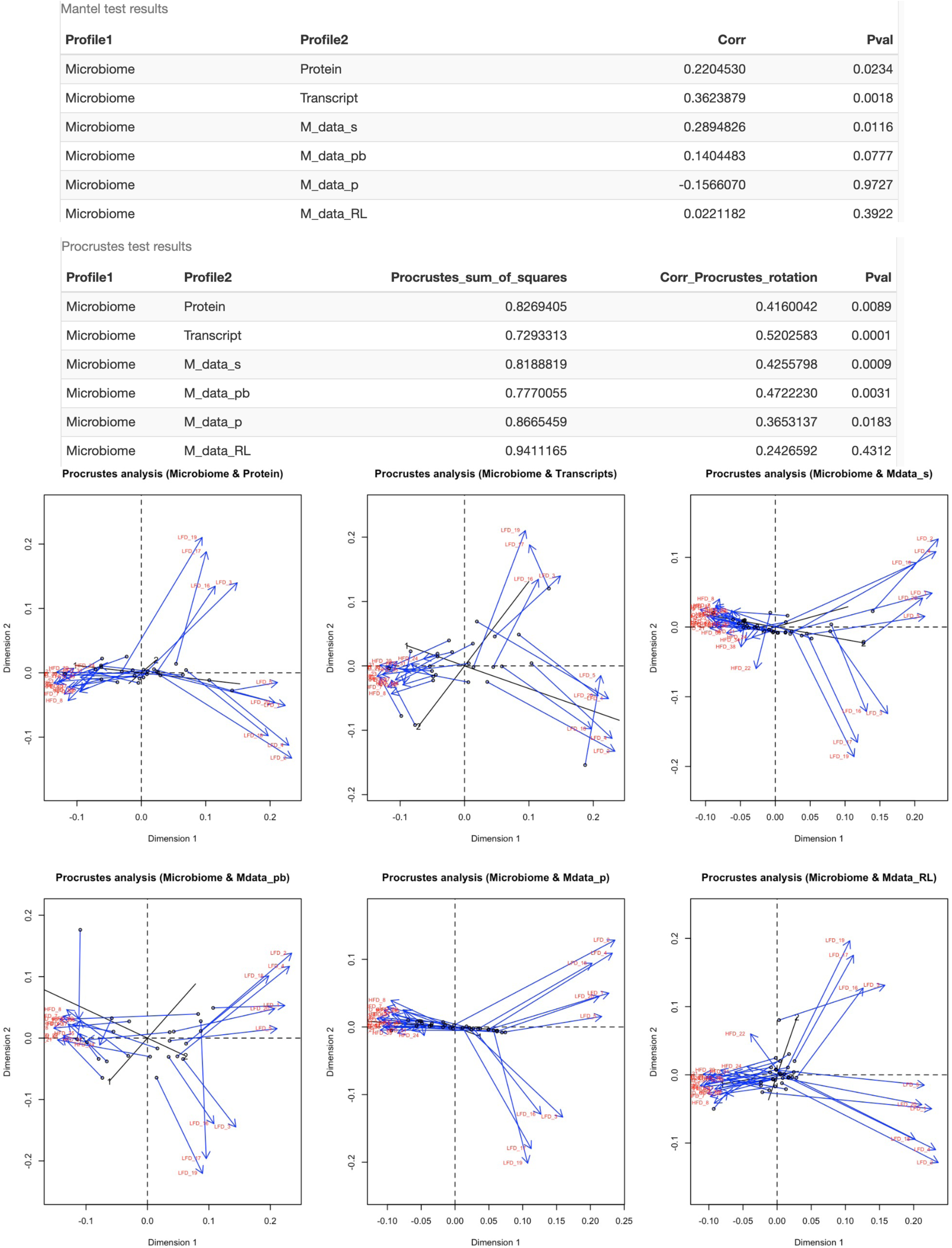
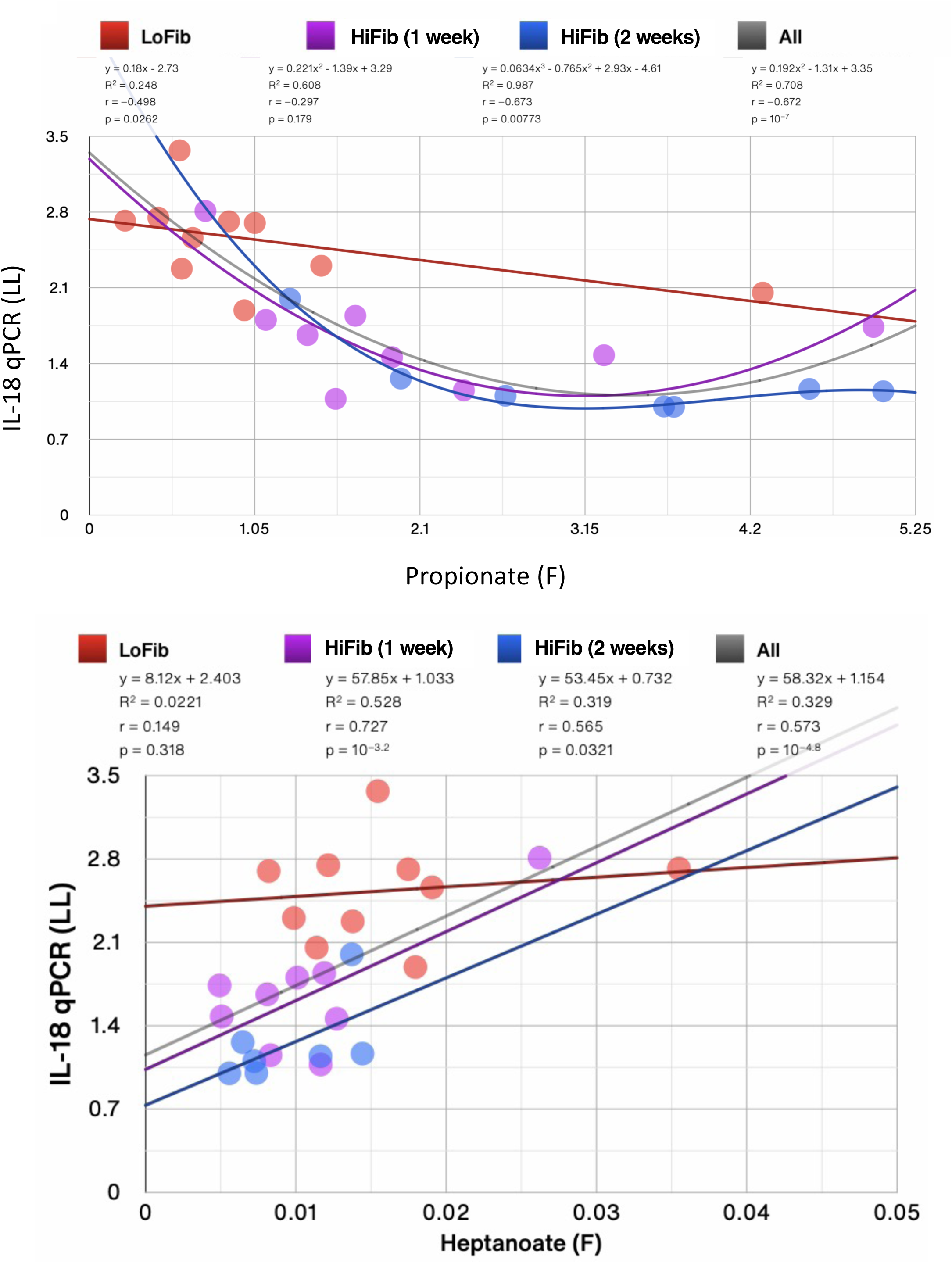
Enrichment of small SCFAs and reduction of lung immune tone markers after HiFib diet. (A) PCA scatter plots visualizing the first 2 principal components with the intensity of points denoting log2 fold change versus LoFib (top). Donut chart showing contributions of the five principal components necessary to explain at least 90% of the dataset’s variance in HiFib diet group and the 9 most correlated measures for each of the first 4 principal components are listed (bottom). Measures in red/blue text are positively/negatively correlated with the respective component. (B) Volcano plot showing increased and decreased metabolites, cytokine/chemokine in all tissues examined in HiFib group versus LoFib. SCFA Acetate in portal blood was the most increased measure and isovalerate in feces was the most decreased measure in HiFib diet group. Green lines denote significance cut off levels (lighter to darker, p<0.05, p<0.01, and p<0.001). (C) Venn diagram depicting significantly changed immunological measures in 1-week and 2-week HiFib groups, respectively, versus LoFib group. Of the total 47 measures, 11 were changed significantly in both groups, 7 were specific to the 2-week HiFib and 5 to the 1-week HiFib group. In the 2-week HiFib group, 18/47 (∼38%) measures were significantly changed. Of the changed measures, 8/18 (∼44%) increased and 10/18 (∼66%) decreased. In the 1-week HiFib group, 16/47 (∼34%) measures were significantly changed. Of the changed measures, 4/16 (∼25%) increased and 12/16 (∼75%) decreased. Measures in red text denote increased and in blue text decreased log2 fold change versus control (LoFib group). (D) Mantel (top) and Procrustes (middle) analyses to correlate 16S fecal genomic sequencing dataset with inflammatory transcriptomic/proteomic and SCFA metabolomic datasets. Visual representation of Procrustes analysis shown at bottom. (E) IL-18 levels in left lung as measured using qPCR correlated with propionate (top) and heptanoate (bottom) levels in feces. Individual r and p values for each group are shown. Linear trendlines, or an nth-degree polynomial trendline if its goodness-of-fit is either 50% greater than, or if it explains at least half of the variance not explained by the (n – 1)th-degree polynomial, are shown. R^2^, r and p denote goodness-of-fit, Pearson’s correlation coefficient and significance of the correlation, respectively.

Next, we combined all datasets from the three diet intervention groups and were able to observe significant correlations between measured metabolites, 16S taxa, and inflammatory markers. The most striking findings were a strong negative correlation between lung IL-18 mRNA levels and fecal C3 (propionate) levels (p=10^-7^) and a positive correlation with fecal C7 (heptanoate) levels (p=10^-4.8^) (**Figure 3E**). Additional significant correlations with bacterial taxa were observed (**Supplemental Table S1**) with IL-1β mRNA levels negatively correlating with fecal proteobacteria levels (p=10^-6.4^) and IL-18 mRNA with fecal firmicutes levels (p=10^-10.1^) (**Supplemental Figure S8A**). We also noted strong correlations between chemokines and cytokines with each other (IL-1β with CXCL-1, IL-6, CXCL-2,; IL-6 with CXCL-2; IL-1β in the left (LL) and right lungs (RL)) as well as SCFA metabolites with each other (caproate with heptanoate, acetate with propionate, butyrate with propionate, and propionate with heptanoate), and IL-1β mRNA with heptanoate (**Supplemental Figure S8B**).

We performed Spearman correlation analysis to uncover significant correlations between metabolites and microbiota taxa across our various datasets (combining all 3 groups together). By using a liberal FDR p value of <0.1 and even the more stringent FDR p<0.05 and P<0.01, we were able to identify species-level significant correlations (*Parasutterella, Bacteriodes, and Lactococcus*) with SCFA production and both reduced and increased lung immune tone markers (IL-1β and IL-18 mRNA) (**Figure 4A**). At the amplicon sequence variant (ASV) level, we were able to identify unique ASV taxa that simultaneously correlated at FDR p<0.05, p<0.01 and even p<0.001 with fecal and portal blood SCFAs (C2 and C3) as well as levels of IL-1β and IL-18 mRNA in lung tissue (**Figure 4B**). Some unique ASVs correlated with increases in SCFAs and reduced lung immune tone while others with decreased SCFAs, increased BCFAs and MCFAs, and increased lung immune tone. The ASV level findings in relation to SCFA metabolites and lung immune tone markers are summarized in **Supplemental Table S2**.

**Figure 4.**
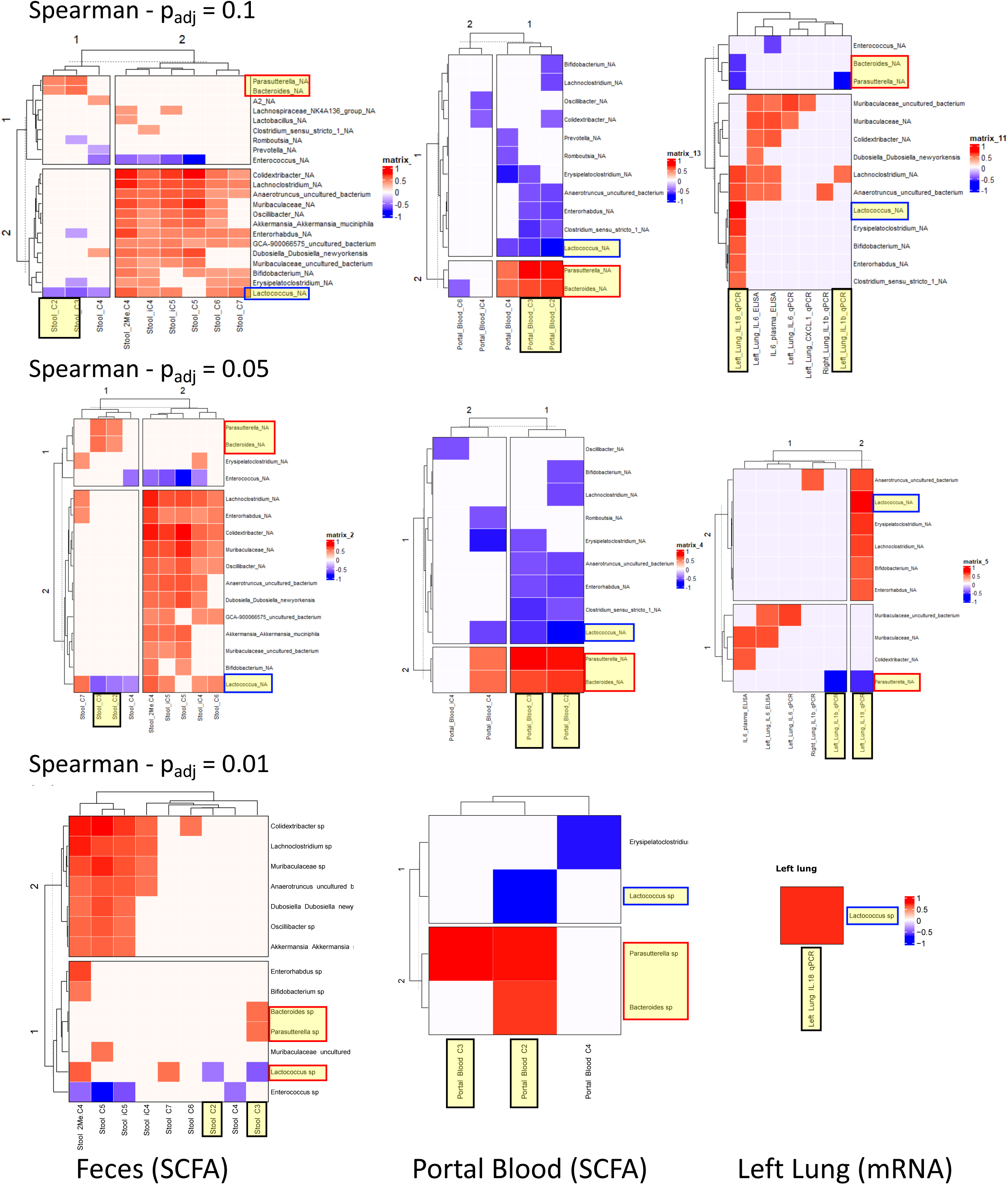
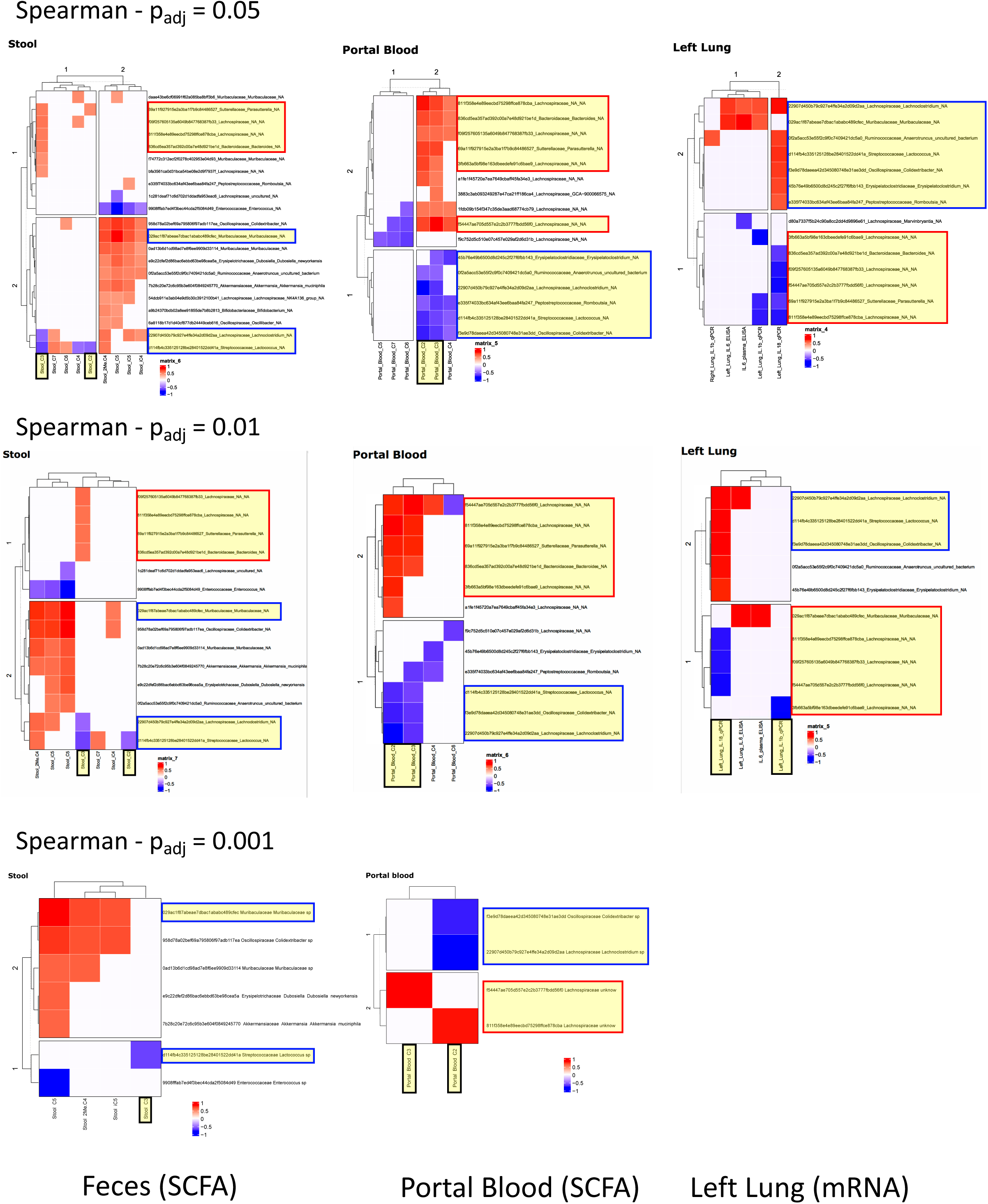
Gut bacteria correlations with SCFA metabolites and with lung immune tone and inflammatory markers. (A) Species level taxa correlations observed with fecal and portal blood SCFA levels and with lung IL-1β, IL-18, IL-6, and CXCL-1 (all three groups combined). Spearman adjusted FDR p value of 0.1 (top), 0.05 (middle) and 0.01 (bottom). Significant correlations of interest are highlighted in yellow: Stool and portal blood C2 and C3, left lung IL-1β and IL-18 mRNA/qPCR (black boxes), Parasutterella and Bacteriodes species (red boxes) and Lactococcus (blue boxes). (B) ASV (amplicon sequence variant) level taxa correlations observed with fecal and portal blood SCFA levels and with lung IL-1β, IL-18, IL-6, and CXCL-1 (all three groups combined). Spearman adjusted FDR p value of 0.05 (top), 0.01 (middle) and 0.001 (bottom). Significant correlations of interest are highlighted in yellow: Stool and portal blood C2 and C3, left lung IL-1β and IL-18 mRNA/qPCR (black boxes), recurring taxa correlating with reduced lung immune tone (red boxes) and increased lung immune tone (blue boxes).

Using our 16S-based microbiota composition analysis, we next applied the PICRUST2 algorithm to infer biological enzymatic pathways and compare our three fiber diet groups. This approach revealed that, similarly to 16S-based microbiota composition analysis, HiFib groups 2 and 3 closely resembled each other and were distinct from the LoFib group 1 (**Figure 5, top**). Delving deeper into specific pathways, we observed that, as expected, pectinesterase was enriched in our HiFib groups (groups 2 and 3; except for the 5 mice in group 2 that were enriched for *E. coli/Shigella* as mentioned earlier) (**Figure 5, bottom**). We also examined KEGG modules that correlate with the K pathways enriched in the HiFib groups vs. the LoFib group and noted specific enriched modules: SCFA production, carbohydrate metabolism, microbial metabolism in diverse environments, oxidative phosphorylation, ABC transporters, etc. (**Figure S9**).

**Figure 5.**
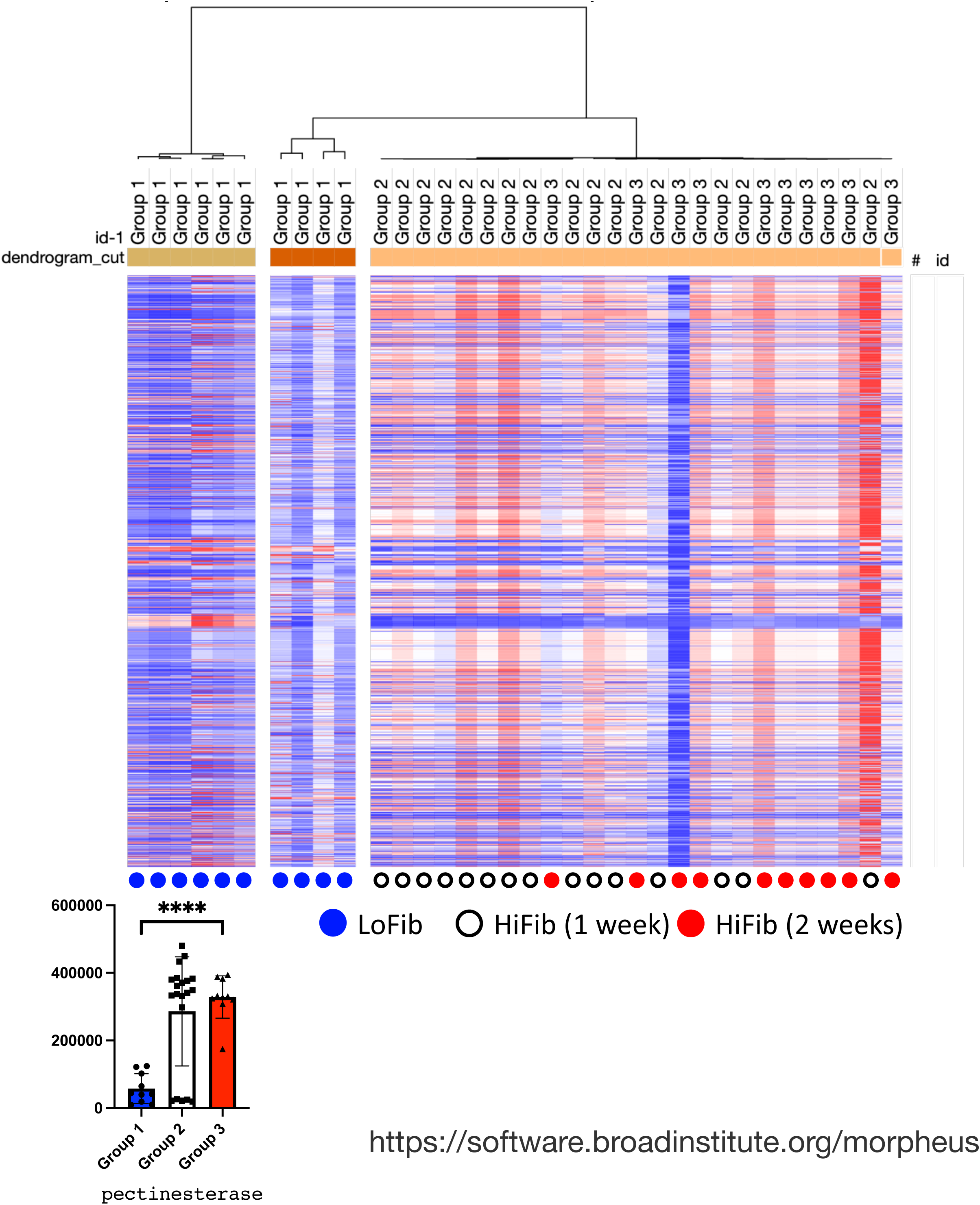
Metabolic profile of low fiber group is distinct from high fiber groups. Using PICRUST2, the metabolic pathways calculated from the 16S profiles of the three fiber groups were compared using web-based Morpheus tool to generate a heatmap (https://software.broadinstitute.org/morpheus) (top). Individual rows represent K value reference pathways/enzymes obtained from https://www.genome.jp/kegg/mapper/ used to identify metabolic pathways and metabolites. Pectinesterase pathway differences between the low fiber and the two high fiber groups (bottom). P values are represented as follows in the figures: *< 0.05; **< 0.01; ***< 0.001; ****< 0.0001.

### Germ-free mice have reduced lung immune tone and lung injury responses

To further understand the role of the microbiome in establishing lung immune tone, we examined the levels of IL-1β and IL-18 mRNA in lungs from steady state uninjured germ-free (GF) mice and studied their lung IR injury inflammatory responses. Baseline lung IL-6 expression levels were unchanged reflecting the absence of systemic inflammation in lungs from both GF and SPF mice (data not shown). We observed that GF mice had reduced IL-1β but not IL-18 mRNA (**Figure 6A**) and significantly reduced lung IR injury responses (**Figure 6B**). Thus, it would appear that gut microbiota are required to establish IL-1β (but not IL-18) lung immune tone while the specific composition of gut microbiota can affect the levels of both IL-1β and IL-18 lung immune tone.

**Figure 6.**
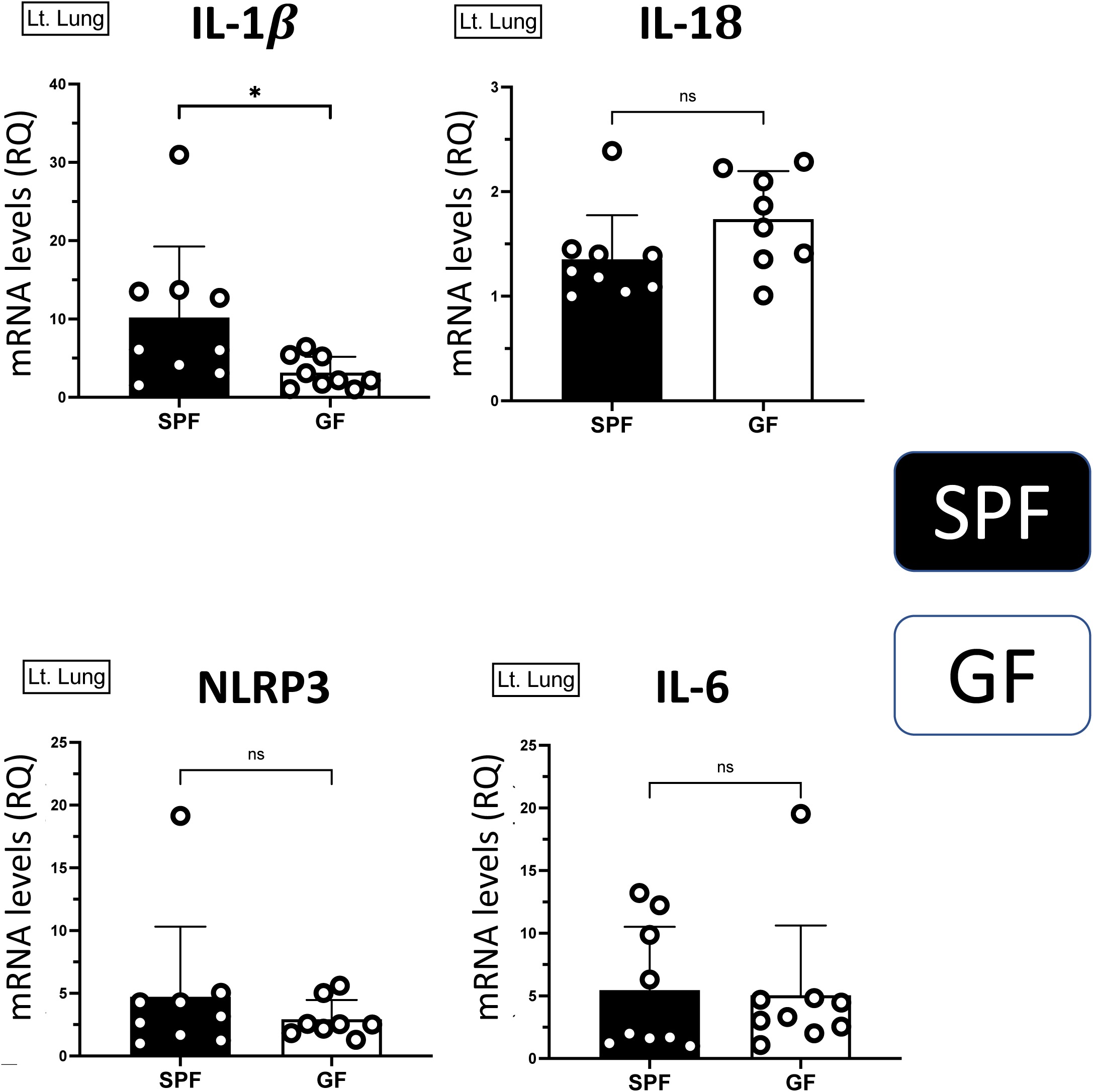
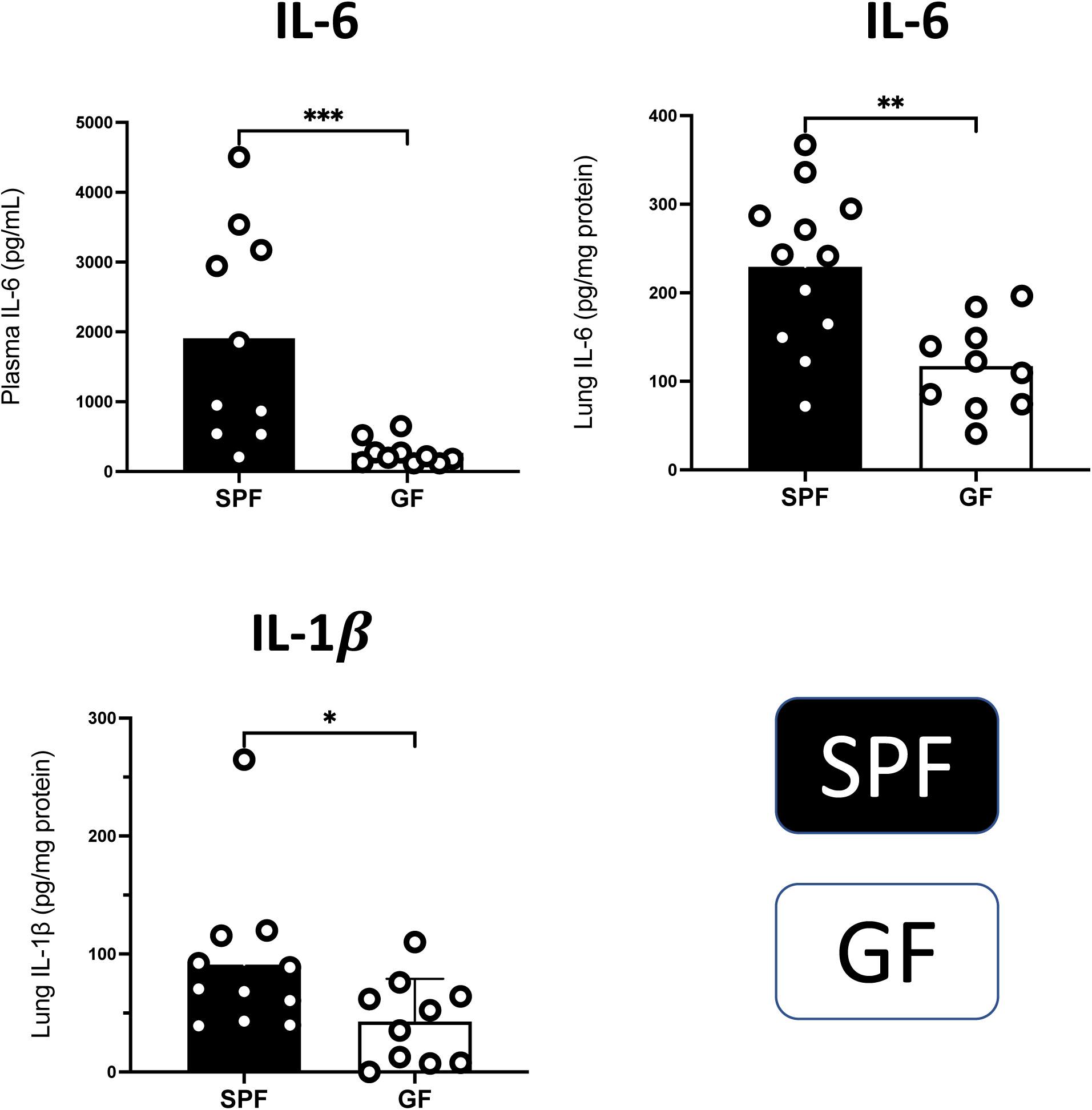
Germ-free (GF) mice have reduced IL-1β lung immune tone and reduced inflammation after IR injury. (A) lung immune tone markers (IL-1β, IL-18, NLRP3 mRNA levels) were measured in uninjured SPF or GF mice. IL-6 levels were measured to examine baseline differences in lung inflammation if any. (B) lung inflammation markers after IR injury were measured in plasma (IL-6) and injured left lung tissue (IL-6 and IL-1β protein levels). P values are represented as follows in the figures: *< 0.05; **< 0.01; ***< 0.001; ****< 0.0001.

### SCFA regulate alveolar macrophage lung immune tone and metabolic programming

To further explore the mechanistic basis for SCFA regulation of lung immune tone, we treated alveolar macrophages (AMs) with propionate (C3), i.e. the SCFA that we found to be highly correlated with lung immune tone in our *in vivo* experiments. We noted that C3 pre-treatment prior to LPS challenge resulted in a dose dependent regulation of the immune tone markers, IL-1β and IL-18, while NLRP3 levels were unchanged (**Figure 7A**). Exposure of AMs to C3 during LPS challenge was also able to specifically reduce inflammatory cytokine (IL-6) output, maintain growth factor (GM-CSF) levels and strikingly increasing anti-inflammatory IL-10 levels (**Figure 7B**). These data support our *in vivo* findings that C3 can regulate lung immune tone and are consistent with our previously published work that alveolar macrophages help establish baseline lung immune tone *in vivo* by expressing IL-1β and IL-18 mRNA and initiate and integrate lung IR injury responses(Liu et al., 2021; Prakash et al., 2012; Tian et al., 2019).

**Figure 7.**
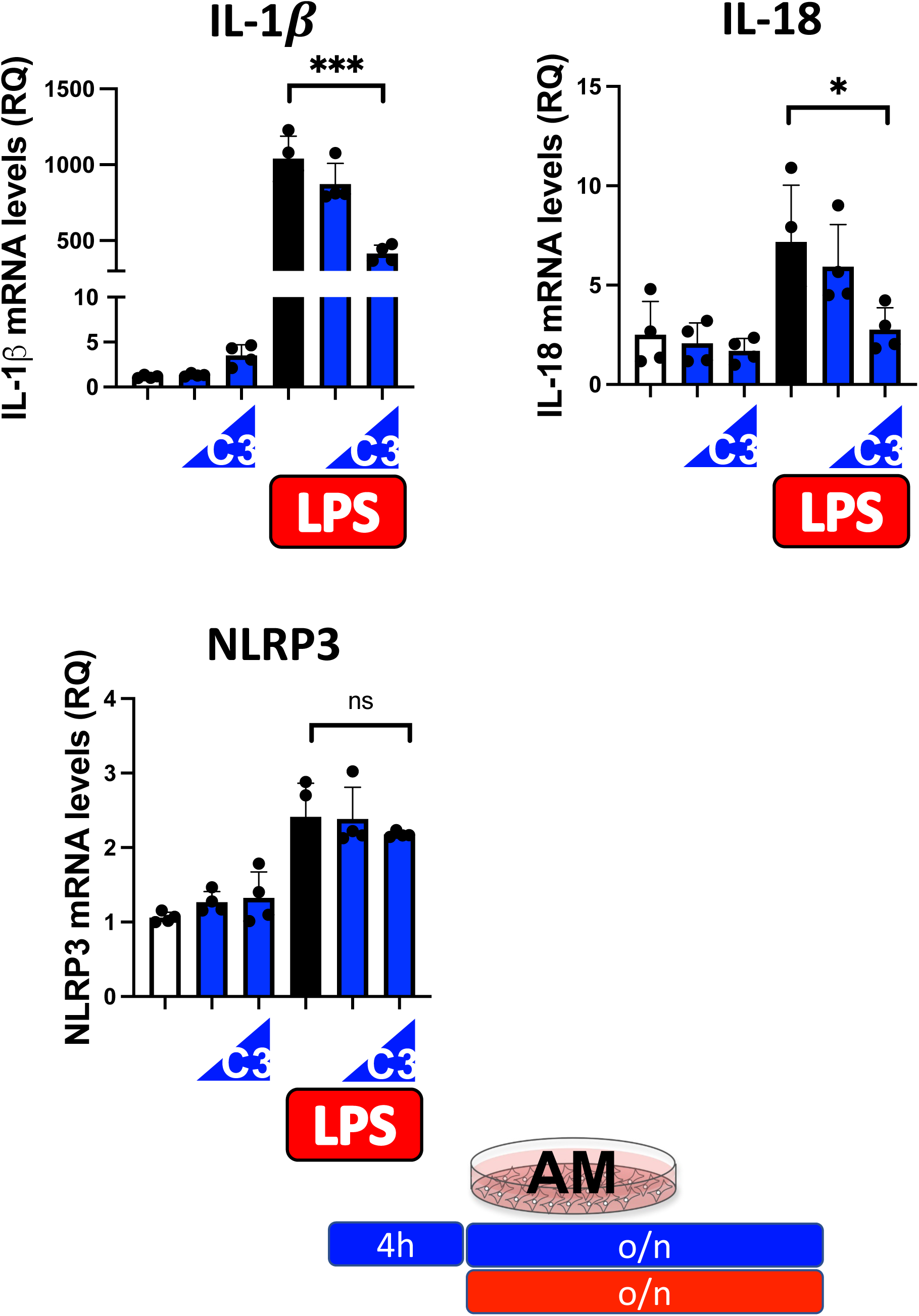
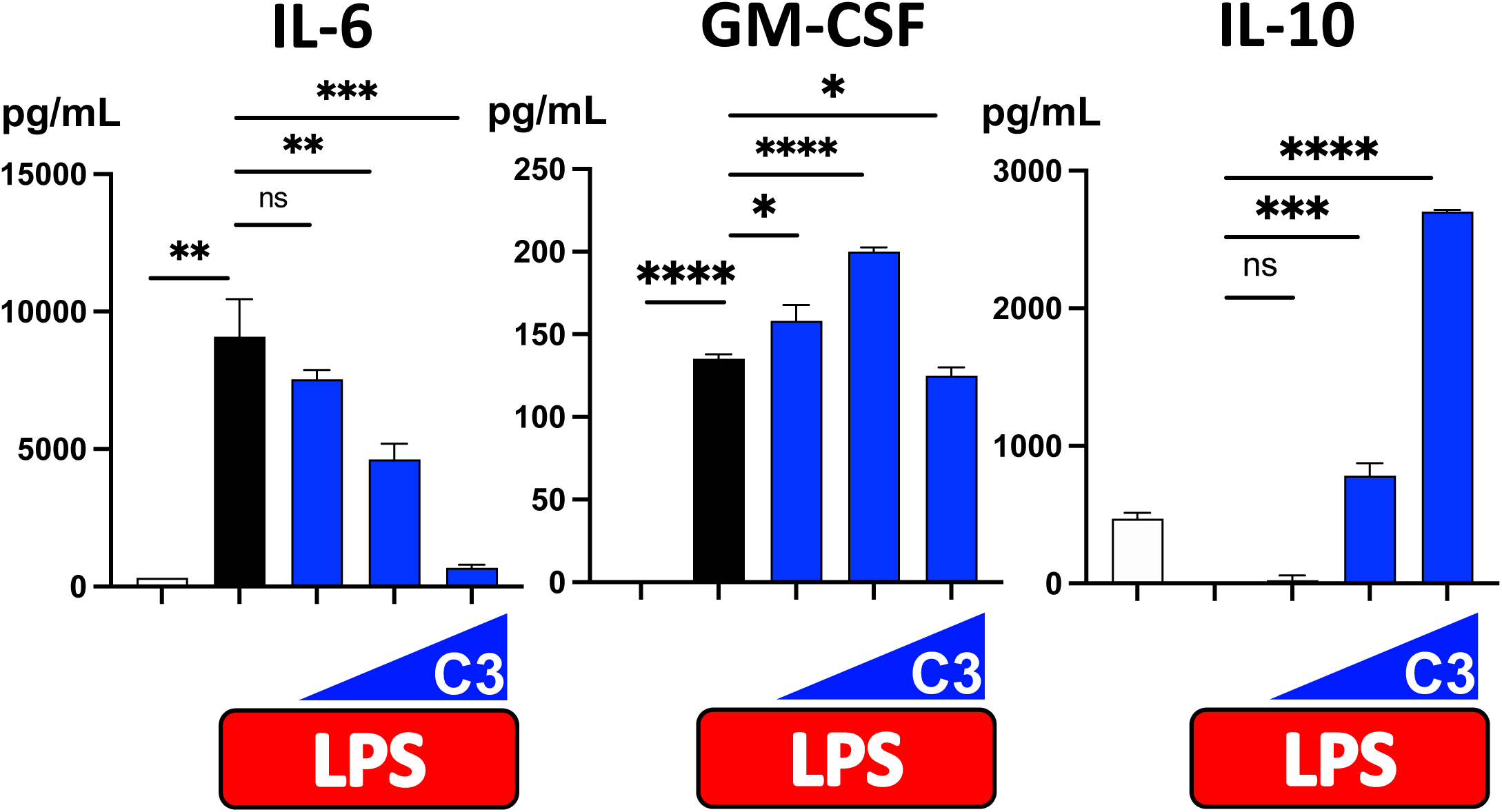
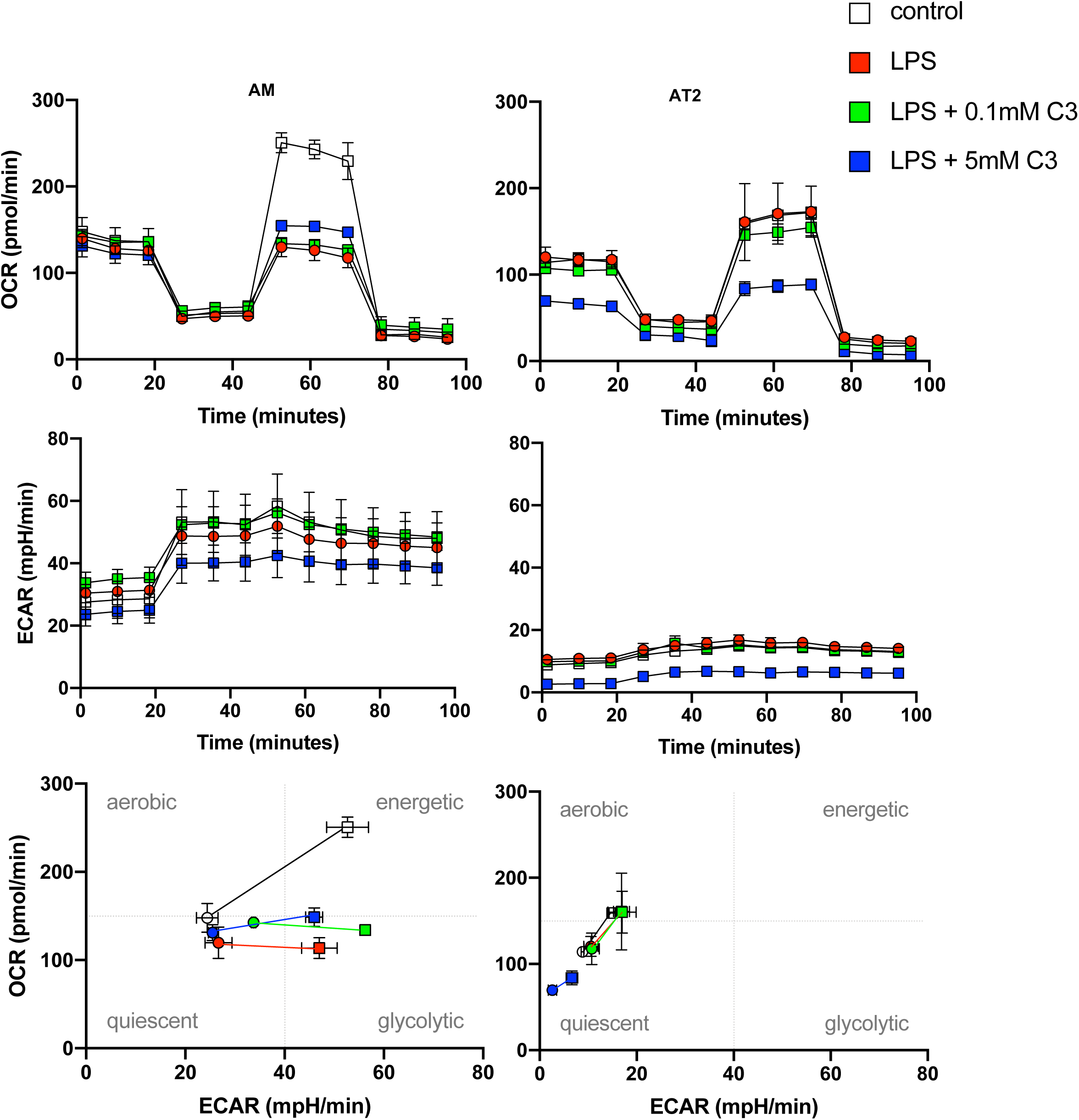
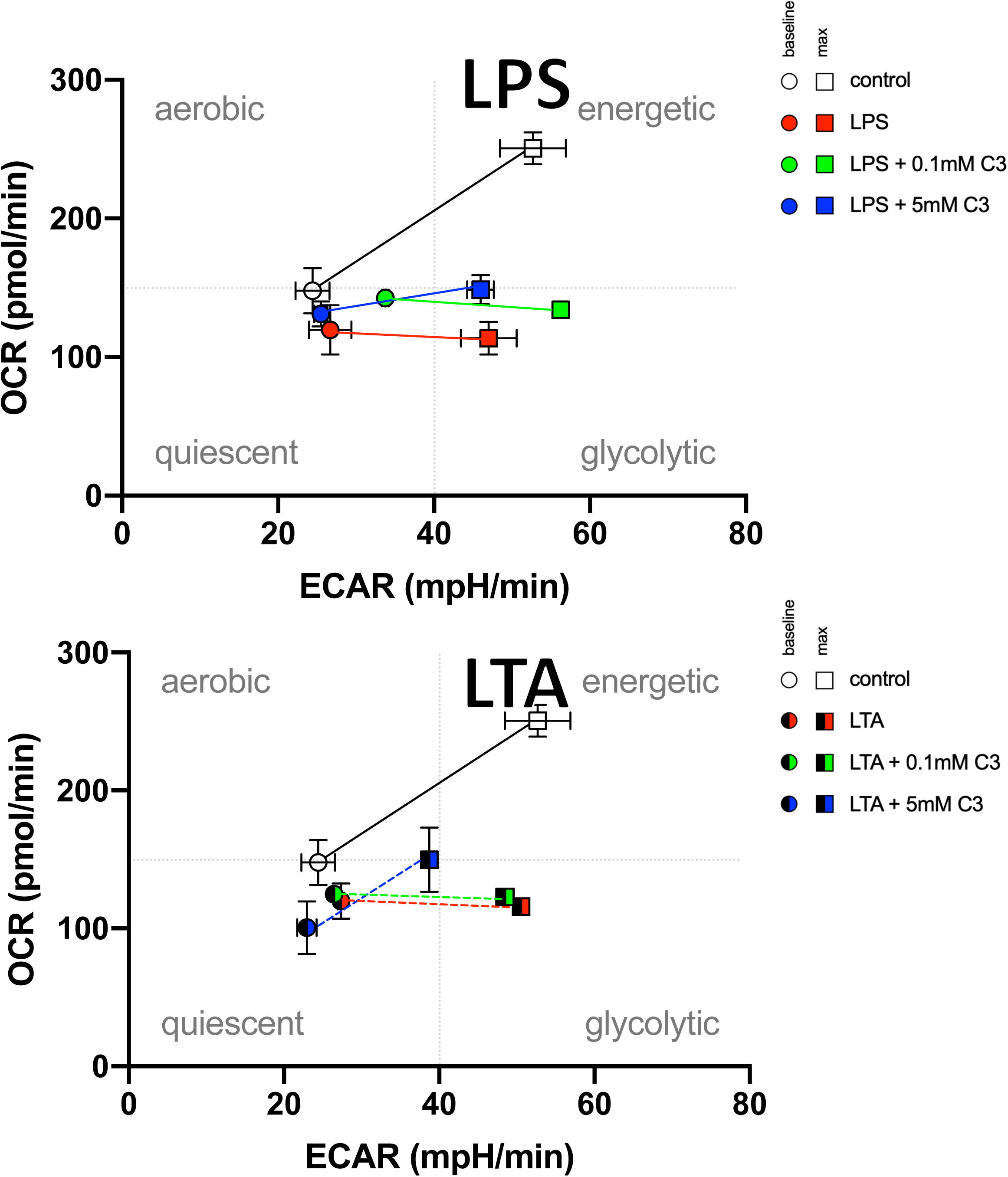
Propionate (C3) can regulate alveolar macrophage immune tone, inflammatory injury output, and metabolic programming *in vitro*. (A) The alveolar macrophage cell line (MH-S) was pre-treated with different doses of C3 (0.1mM and 1mM) in the presence of LPS and markers of immune tone were measured (IL-1β, IL-18, NLRP3 mRNA). (B) Effect of increasing amounts of C3 on LPS inflammatory responses in alveolar macrophages *in vitro*. IL-6, GM-CSF and IL-10 protein levels were measured at 24h, 24h and 48h, respectively. (C) Seahorse metabolic profiling of alveolar macrophages treated *in vitro* with LPS (10ng/mL) and C3 (0.1mM and 5mM). OCR is a measure of oxidative phosphorylation and ECAR a measure of glycolysis. (D) Comparison on the metabolic reprogramming of alveolar macrophages by LPS (10ng/mL) and LTA (100ug/mL) and the effects of co-exposure to C3 (0.1mM and 5mM). P values are represented as follows in the figures: *< 0.05; **< 0.01; ***< 0.001; ****< 0.0001.

Given that metabolism is inextricably tied to immune responses, we examined the possible metabolic alveolar reprogramming that might occur in AM and type 2 alveolar epithelial cells (AT2) in the presence of low and high C3 in the context of background low LPS stimulation (Signal 1 of the inflammasome). We observed that AM and AT2 were metabolically programmed very differently at baseline and responded distinctly to the presence of LPS and C3. AMs exposed to LPS favored glycolysis and this was reversed back towards balanced oxidative phosphorylation (energetic state) in the presence of high C3 (**Figure 7C**). Specifically, metabolic potential and spare respiratory capacity were restored in AMs in the presence of high C3, while AT2 had reduced basal and maximal respiration, reduced baseline and stressed OCR and ECAR, reduced non-mitochondrial oxygen consumption and reduced ATP-coupled respiration (**Supplemental Figure S10**). To verify that gram-positive pathogen-associated molecular patterns (PAMP) could function similarly to LPS, we challenged AMs with lipoteichoic acid (LTA) and observed similar if not greater restoration of AM metabolic potential in the context of high C3 (**Figure 7D**).

### Model for fiber regulation of microbiome, metabolites and lung immune tone

Our data supports the strong influence of the gut-lung axis on lung immune tone and response to lung injury. Based on the results we have presented here, we propose, in **Figure 8**, a model for regulation of lung immune tone based on the relative presence of fiber-fermenting, SCFA-producing gut bacterial taxa that transmit these (and likely other) metabolite signals to the lung where they reprogram alveolar macrophages and potentially other resident lung immune cells.

**Figure 8.**
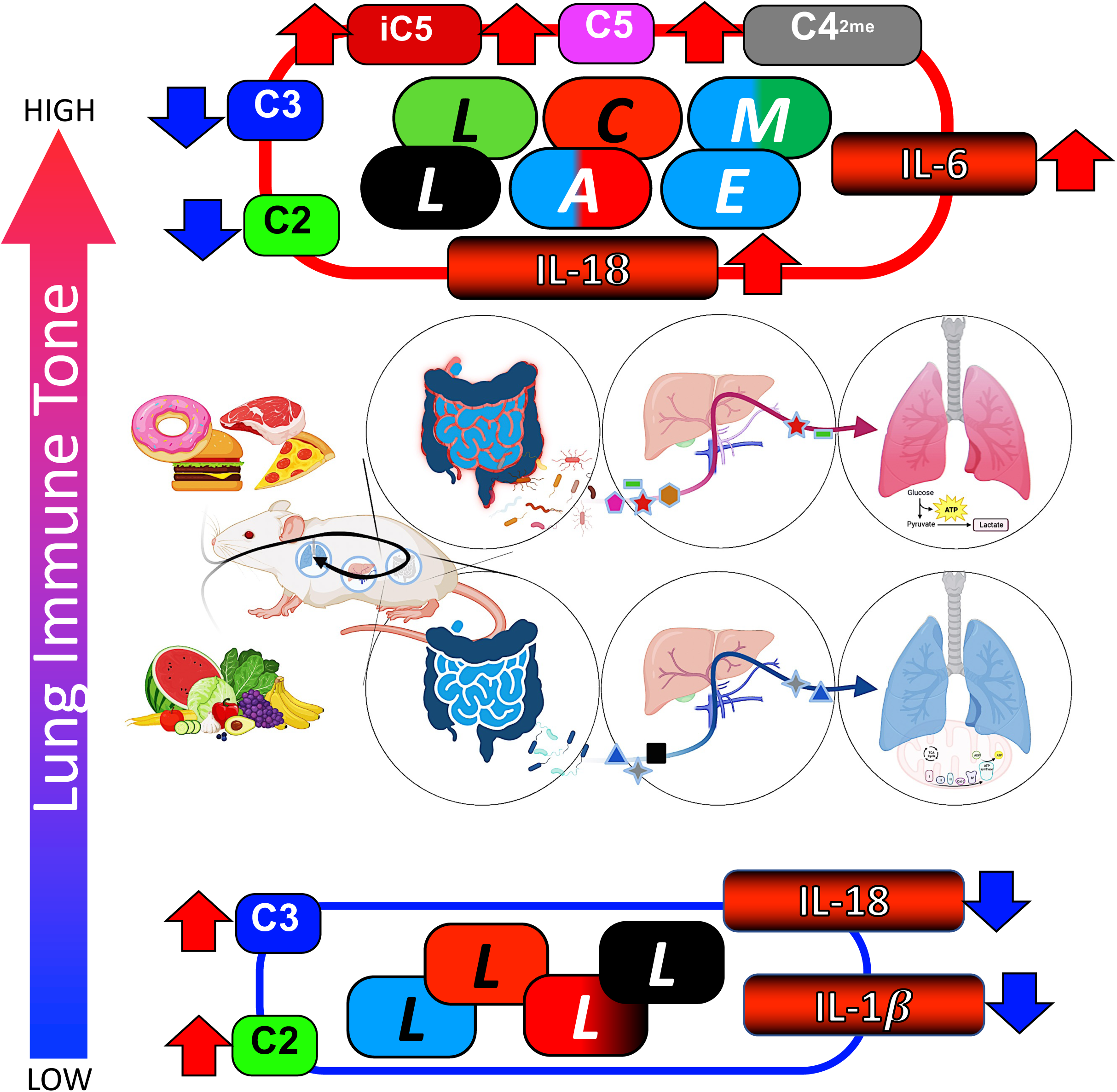
Dietary fiber Gut-Lung Axis Model. Low fiber diets enrich for medium-chain and branch-chain fatty acids (MCFA and BCFA) producing gut bacteria and are depleted for SCFA producers. This correlates with the transit of these pro-inflammatory metabolites to the lung and the establishment of higher lung immune tone. Conversely, high fiber diet promotes the enrichment of fiber-fermenting species that produce acetate and propionate that are transported systemically and possibly consumed in the lung where they might help establish reduced lung immune tone. Overall, these level of lung immune tone inform the downstream inflammatory response to injuries such as ischemia reperfusion.

## DISCUSSION

The major conclusions of the current study are that dietary pectin fiber can alter lung immune tone and sterile lung injury early inflammatory responses and that specific bacterial taxa and SCFA metabolites can mediate gut-lung axis regulation of lung immune tone. These conclusions are based on the following experimental evidence. First, we demonstrated that early inflammatory cytokine and chemokine production following sterile lung IR injury was reduced in mice exposed to high fiber diets for 2 weeks. These mice had reduced lung immune tone as measured by IL-1β and IL-18 mRNA levels and these levels correlated strongly with the abundance of specific fermenting bacterial taxa and the SCFAs propionate and acetate in feces, portal blood, and circulating plasma. Next, we provided strong evidence that germ-free mice lacking microbiota had reduced IL-1β lung immune tone as well as reduced lung IR injury inflammatory responses. Finally, we demonstrated that propionate, at higher levels, could directly reduce alveolar macrophages’ immune tone and inflammatory responses as well as alter their metabolic programming. Collectively, this experimental evidence strongly supports the concept that diet and the gut-lung axis play important roles in regulating lung immune tone and lung injury responses.

Our prior research demonstrated that lung immune tone could be measured by the presence of IL-1β mRNA levels at baseline and that LPS and SCFAs could transit from the gut to the lung (Liu et al., 2021). Furthermore, our *in vitro* investigations illustrated that different levels of propionate could either prime or suppress alveolar macrophage inflammatory responses and high levels of propionate producing bacteria coincided with reduced lung IR inflammation (Tian et al., 2019). The significance of lung immune tone was also demonstrated by the fact that interrupting NLRP3 inflammasome mediated IL-1β production or sensing was associated with reduced lung IR inflammation and worse bacterial clearance as part of a superimposed pneumonia(Tian et al., 2017). Other researchers have demonstrated the importance of the gut microbiota in combating lung infections [(Galvão et al., 2018; Sencio et al., 2020) and reviewed in (Cruz et al., 2021)] and we demonstrated that LPS sensing via TLR4 was also required for full lung IR inflammatory responses(Prakash et al., 2012). Two other groups used high pectin fiber and observed altered lung immunity and bone remodeling with changes in the gut microbiome (Lucas et al., 2018; Trompette et al., 2014). Importantly, both noted increases in SCFAs in circulation and identified direct or indirect effects of SCFAs on the physiology and pathology of target organs of interest.

Recent work has reported an alteration in type 2 inflammation following high fiber diet with worse allergic lung responses and improved parasite clearance in mice (Arifuzzaman et al., 2022). This paper implicated ILC2s, IL-33, cholic acid, and the farnesoid X receptor (FXR) as driving the eosinophilia observed in their system. Interestingly, this group used inulin as their fiber source but did not detect significant elevations in fecal propionate and butyrate levels in their study. Therefore, this study was not able to examine the effects of increased SCFA-producing fermenting gut bacterial taxa on lung immune tone and injury responses, in addition to other differences in their lung injury models. Work from Andrew Gewirtz and Benoit Chassaing (contributor to this study) has shown that different fiber sources (inulin vs. pectin vs. psyllium) may have distinct effects on gut immunity with some fiber sources being beneficial and others deleterious depending on the injury model (Zou et al., 2018). We used pectin as the fiber source based on prior work that demonstrated strong effects on the lung immune responses and bone pathophysiology (Lucas et al., 2018; Trompette et al., 2014) and with pectin we were able to observe elevated SCFA levels in feces, portal blood and circulating plasma (Figure 2C). Unlike other published work, we looked for and found strong correlations between SCFA producing bacterial taxa and acute lung injury immune tone markers, namely IL-1β and IL-18 mRNA. Interestingly, the regulation of IL-1β and IL-18 appeared to be distinct as well, with only IL-1β being dependent on the presence of microbiota while both IL-1β and IL-18 appeared to be strongly modulated by the presence of acetate, propionate, heptanoate, and possibly other fatty acid metabolites. Our focus on alveolar macrophages as the key cell expressing basal IL-1β mRNA and regulating lung immune tone in our model is based largely on our prior work (Liu et al., 2021) but is also supported by work from other groups focusing on alveolar macrophages in IR lung injury (McCourtie, Farivar, Woolley, Merry, Wolf, Mackinnon-Patterson, Keech, Fitzsullivan, & Mulligan, 2008; McCourtie, Farivar, Woolley, Merry, Wolf, Mackinnon-Patterson, Keech, Fitzsullivan, Mulligan, et al., 2008; Shi et al., 2012) and very recent work from on alveolar macrophages as being a key cell type in the gut-lung axis from Andrew Gewertz’s group which showed that gut microbiota can program alveolar macrophages and affect lung injury responses (private communication, manuscript in submission).

Our study begins to fill in important gaps in knowledge on how the gut microbiome influences lung injury responses via the gut-lung axis. Our findings that specific bacterial taxa may use SCFAs to regulate lung immune tone has important implications for human health, disease treatment, and prevention strategies. Now, we may begin to probe the effects of specific bacteria or consortia to alter lung immune tone and downstream injury responses. Moreover, compared to fecal microbiome transplantation (FMT), the ability of fiber-rich diets to shape the landscape of the gut microbiome elevates dietary interventions as a potential powerful and simple tool to modulate lung immune tone. The direct administration of SCFAs to alter lung immune tone and other organ responses has been already demonstrated by us and others(Lucas et al., 2018; Tian et al., 2019) and the widespread therapeutics applicability of SCFAs as supplements or the ingestion of SCFA-rich foods needs to be further investigated. Overall, compared to FMT and direct SCFA administration (via pure compounds or foods rich in them), the use of dietary fiber to enrich for fermenting gut bacteria may be holistic, simpler, and perhaps preferrable. Specific diets (not just based on fiber) could be envisioned to shape organ health, resilience, and immunity both in health, prior to injury, and in disease.

The reduction of inflammatory responses downstream of sterile lung IR injury may be advantageous in clinical scenarios such as organ transplantation, reperfusion after pulmonary embolus, or even hypovolemic trauma. However, much remains to be studied on the effects of high fiber and reduced lung immune tone vis-à-vis other common lung injuries and diseases – such as pneumonia, asthma, fibrosis, etc. While it is possible that suppressing inflammation could be detrimental in situations of infection, our data shows that propionate does not indiscriminately suppress all immune responses and may instead create an overall healthy state for alveolar macrophages. Other groups have reported increased viral and bacterial killing by macrophages in the presence of specific SCFAs(Antunes et al., 2019; Niu et al., 2023).

A limitation of our study is the use of a single source of fermentable fiber, namely pectin. This fiber source from fruits may act differently compared to other sources and these differences need to be further investigated and understood. Inulin was shown to promote type 2 immunity and help resolve influenza infections while restricting immune pathology. It remains to be studied if pectin can similarly improve lung immune response to bacterial and viral infection.

A very short duration of fiber diet exposure was shown to be effective. One or two weeks appeared to be sufficient to reprogram lung immune tone *in vivo*. More work needs to be done to understand how plastic this immune tone and alveolar macrophage programming is. One week of low fiber diet was not sufficient to prevent the subsequent high fiber exposure from reducing lung immune tone; it would be of great interest to understand the duration of these effects once the fiber source is removed, i.e. the half-life of this immunometabolic programming in the lung.

Other metabolites besides SCFAs, MCFAs, and BCFAs are likely important as well. Our study is limited by its measurement of these and not other larger, different metabolites from the other classes of metabolites (primary and secondary bile acids, tryptophan derivatives, lipids, vitamins, etc.). It is very plausible that SCFAs would combine with any of the other classes of gut-derived and diet-derived metabolites and factors to influence lung immune tone. These interactions need to be studied in a systematic unbiased fashion. Our future work involved direct metatranscriptomic analysis of the fecal contents to better understand the metabolic pathways enriched in our groups and thereby target other metabolite pathways that might also be regulated by fiber diets.

The effect of fiber diets on human physiology and disease conceptually should not be limited to just the lung. Other groups have investigated the gut-brain, gut-liver, gut-bone, and other axes, and it is very likely that the enrichment of fiber-fermenting gut bacteria will have profound effects on the physiology and immune tone of a variety of organ systems in the body.

The major conclusions of this study are that dietary fiber can regulate lung immune tone and downstream lung injury responses. Specifically, to the field of sterile lung injury, these findings have broad implications on the use of dietary interventions prior to general surgery, lung transplantation and other anticipated events involving lung damage. Further understanding of specific bacteria that interact with and manipulate lung immune tone could permit the identification of resilient patient populations as well as at risk populations that might benefit from fecal microbiota transplantation (FMT) with lung-protective taxa. The use of specific metabolites to directly manipulate the system may also have a place in emergency situations when short-term manipulation may be in the patient’s best interests to boost or suppress lung immune responses. Overall, the use of dietary interventions to wholistically modulate lung and other organ function and immune tone is of great general health and therapeutic interest.

## ACKNOWLEDGMENTS

We would like to acknowledge the following individuals for assistance with providing lung tissue, reagents, mice, advice, helpful discussions, and critical reading and editing of the manuscript: Susan Lynch (UCSF), Judith Hellman (UCSF), Michael Matthay (UCSF), Mervyn Maze (UCSF). We also acknowledge the contributions of the BCMM and NORC (Gnotobiotic) core facilities at UCSF. The authors would also like to acknowledge the continuing support of THFC (COYS) on these studies.

