## Supplemental Figures and Tables Legend for "Regulation of Lung Immune Tone by the Gut-Lung Axis via Dietary Fiber, Gut Microbiota, and Short-Chain Fatty Acids"

**Supplemental Figure S1. Diet intervention schematic.** Mice purchased from The Jackson Laboratory were fed normal chow (4% fiber) for 1 week. Prior to the diet switch, feces were collected and stored for 16S sequencing. The diets were then changed to either 0% fiber (LoFib) or 35% pectin (HiFib) fiber for 2 weeks. Then mice underwent left lung ischemia reperfusion injury (1h ischemia, 1h reperfusion) after which the feces, portal blood, circulating plasma, left and right lung tissue were collected for analyses (as indicated).

**Supplemental Figure S2. Lung immune tone and lung injury inflammation measurements.** Schematic representing NLRP3 regulation of IL-1b and IL-18 production in the context of low lung immune tone (top) and high lung immune tone (bottom). IL-1b and IL-18 mRNA and latent forms of their proteins (pro-IL-1b and pro-IL-18) are first produced in response to Signal 1 which sets up the state of lung immune tone. The level of lung immune tone in our studies is quantified by the mRNA levels of IL-1b and IL-18. After injury (e.g. IR injury), which serves as Signal 2, the NLRP3 inflammasome complex is activated and results in the processing of IL-1b and IL-18 into their active forms and eventual release. Measurements of IL-1b and IL-18 along with other downstream early inflammatory cytokines and chemokines (IL-6, CXCL-1, CXCL-2) are used in our studies to quantify lung injury.

**Supplemental Figure S3**. Changes in gut microbiota composition after fiber diet intervention at the phylum, family and genus levels.

**Supplemental Figure S4**. Enriched and depleted taxa in LoFib and HiFib groups (compared to t=0).

**Supplemental Figure S5**. Alpha-diversity comparisons of mice on normal (4% fiber) chow (t=0), LoFib and HiFib groups (after 2 weeks of diet change).

**Supplemental Figure S6**. **Similar significant changes in SCFAs and immune tone markers in 1 week fiber group (LoFib->HiFib) compared to 2 week group (HiFib).** (A) Experimental outline comparing LoFib group (top) with 1 week vs 2 week HiFib groups (middle and bottom). (B) PCA scatter plots visualizing the first 2 principal components with the intensity of points denoting log2 fold change versus LoFib (top) and donut chart showing contributions of the seven principal components necessary to explain at least 90% of the dataset’s variance in 1 week HiFib diet group and the 9 most correlated measures for each of the first 4 principal components are listed. Measures in red/blue text are positively/negatively correlated with the respective component. (bottom). (C) Volcano plot showing increased and decreased (p < 0.1) metabolites, cytokine/chemokine in all tissues examined in 1 week HiFib group versus LoFib. SCFA Acetate in portal blood was the most increased measure and isovalerate and valerate in feces were the most decreased measure in 1 week HiFib as also seen for 2 week HiFib diet group. Green lines denote significance cut off levels (lighter to darker, p<0.05, p<0.01, and p<0.001).

**Supplemental Table 1**. **Significant correlations between cytokine/metabolite data and 16S bacterial taxa.** Correlation table (SIVLA) for measured analytes (all three groups included: LoFib and the 2 high fiber groups) with phylum level 16S data and other analytes (top) and curated list with significantly correlated bacterial species and metabolites (bottom). ***, ** and * denote p < 0.001, 0.01 and 0.05, respectively (also denoted by darker to lighter blue shading). Pearson’s correlation coefficient shown for each significant correlation.

**Supplemental Figure S8**. **Additional PICRUST2 comparison of metabolic pathways showing differences between LoFib and HiFib groups.** Other specific key pathways examined for differences between the three fiber groups. K value reference pathways/enzymes obtained from <https://www.genome.jp/kegg/mapper/> used to identify metabolic pathways and metabolites. P values are represented as follows in the figures: *< 0.05; **< 0.01; ***< 0.001; ****< 0.0001.

**Supplemental Figure S7**. **Selected correlations between cytokine/metabolite data and 16S bacterial taxa.**

(A) IL-1b and IL-18 levels in left lung as measured using qPCR correlated with protobacteria (top) and firmicutes (bottom) levels in feces, respectively. (B) Other significant correlations between lung immune tone and post-lung injury inflammatory marker levels including the contributions of individual dietary groups and all groups combined. Individual r and p values for each group are shown. Linear trendlines, or an nth-degree polynomial trendline if its goodness-of-fit is either 50% greater than, or if it explains at least half of the variance not explained by the (n – 1)th-degree polynomial, are shown. R^2^, r and p denote goodness-of-fit, Pearson’s correlation coefficient and significance of the correlation, respectively.

**Supplemental Table 2**. Summary of ASV level linear correlations with SCFA metabolites and lung immune tone/inflammatory markers. Upward/downward arrows indicate increased/decreased levels and outline of arrow represent Spearman adjusted p values: 0.05 (black), 0.01 (red), 0.001 (green).

**Supplemental Table S3. Comparison of metabolism measurements in alveolar macrophages (AM/MH-S) and type 2 alveolar epithelial cells (AT2/MLE-12).** Seahorse metabolic profiling of alveolar macrophages treated in vitro with LPS (100ng/mL) and C3 (0.1mM and 5mM). P values are represented as follows in the figures: *< 0.05; **< 0.01; ***< 0.001; ****< 0.0001.

**Supplemental Figure 9.** (A) Stool and colon morphology (top and middle), and weight gain/loss comparisons for LoFib vs. HiFib groups (bottom). Composition of t=0 starting diet (B), low fiber (C), and high fiber (D) diets.
