## Supplemental Figures and Tables for "Regulation of Lung Immune Tone by the Gut-Lung Axis via Dietary Fiber, Gut Microbiota, and Short-Chain Fatty Acids"

### Figure S1: Lung Immune Tone and Lung Injury Inflammation Measurements

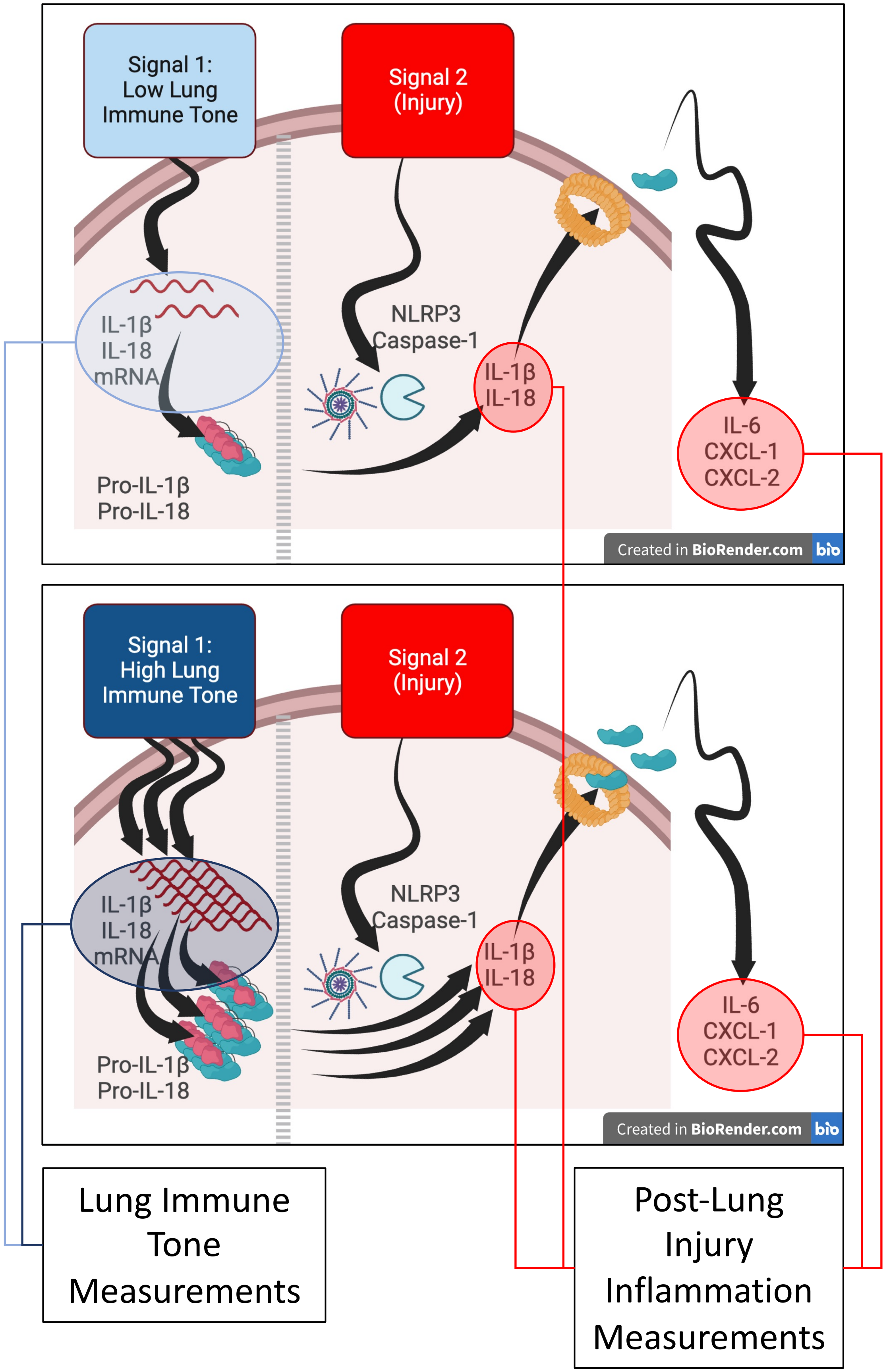

### Figure S2: Diet intervention Schematic

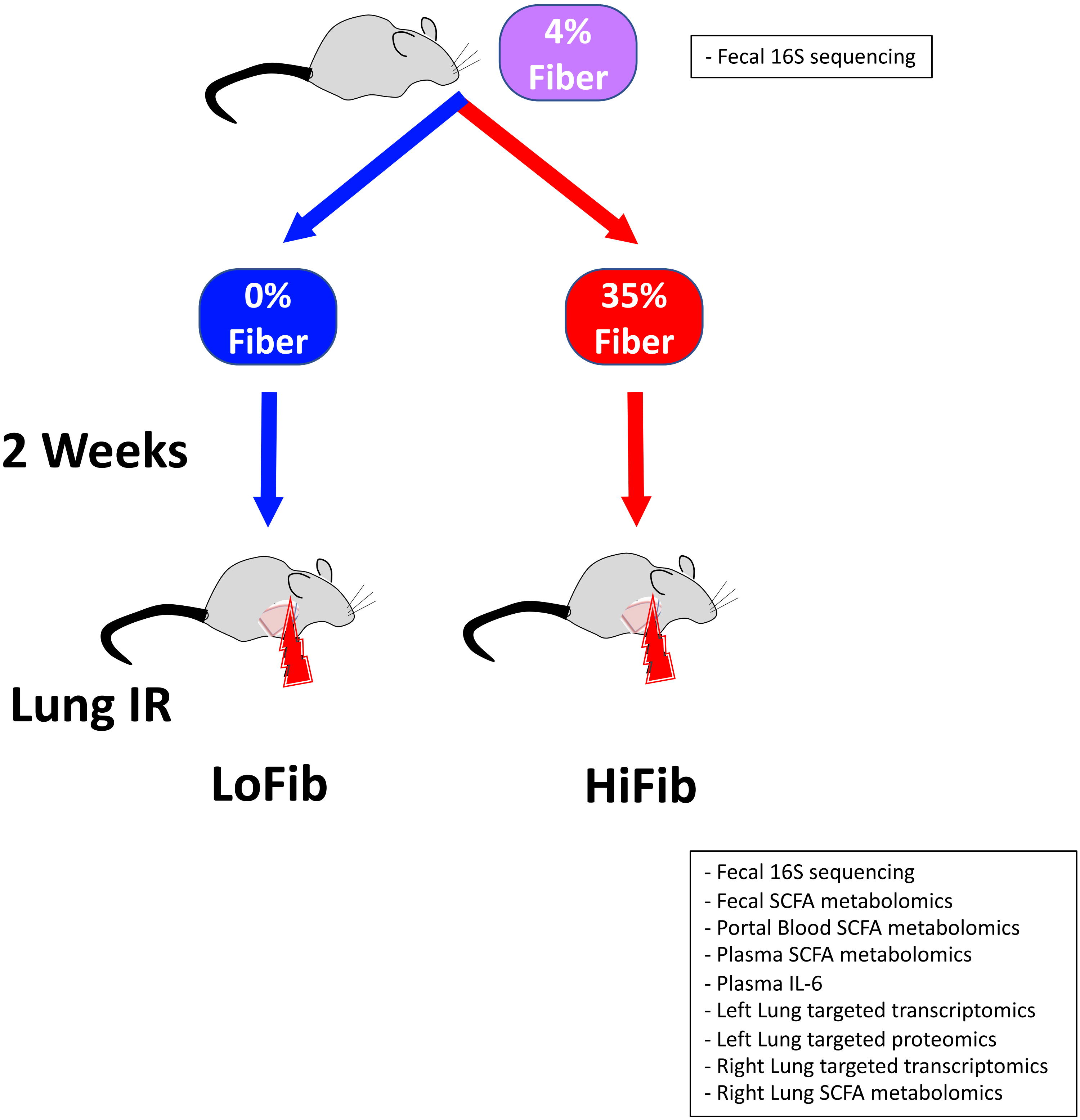

### Figure S3: 16S phylum, genus, and family level changes with fiber diet intervention

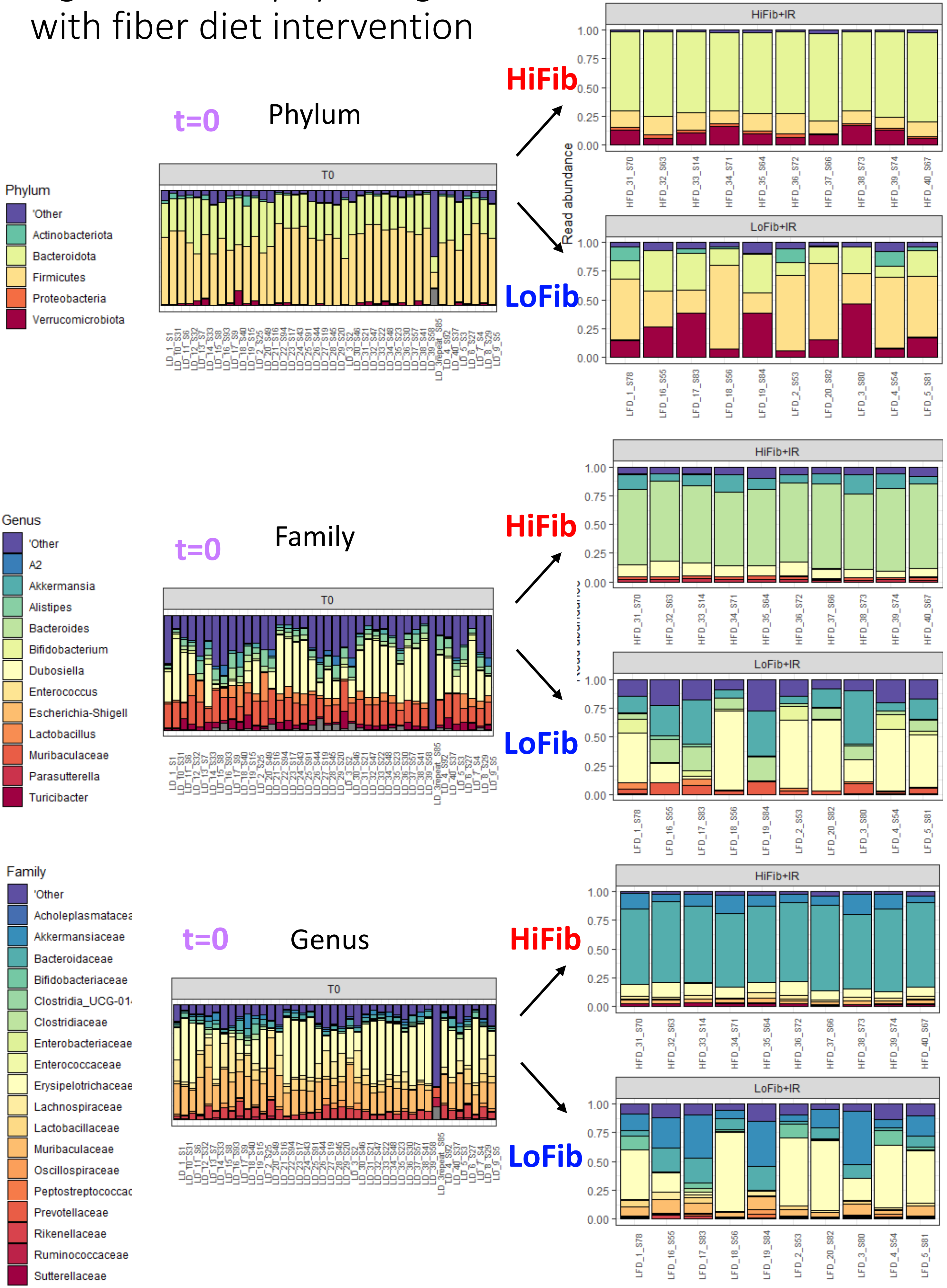

### Figure S4A: Enriched and depleted species in LoFib and HiFib groups (vs t=0)

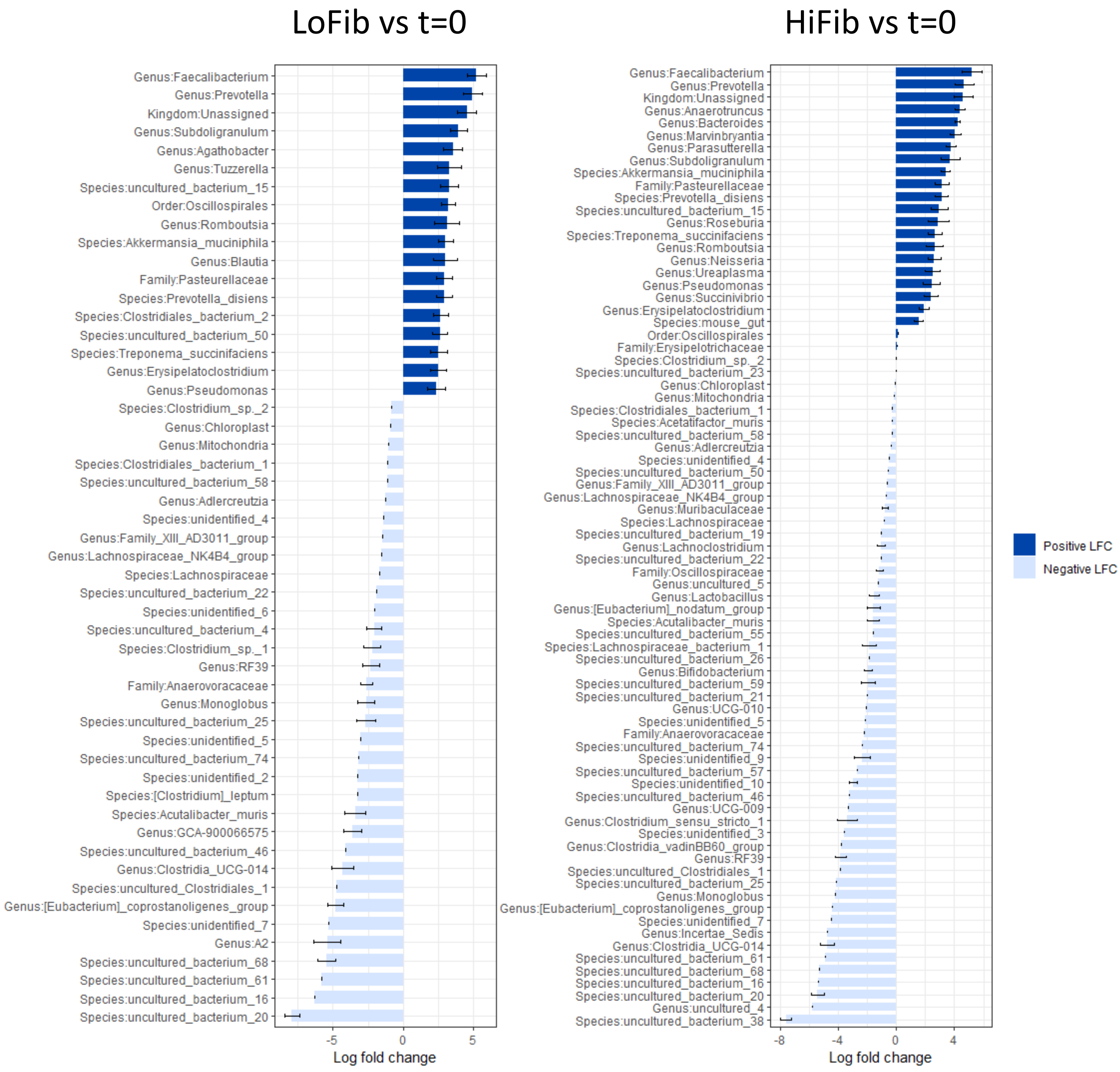

### Figure S4B: alpha diversity

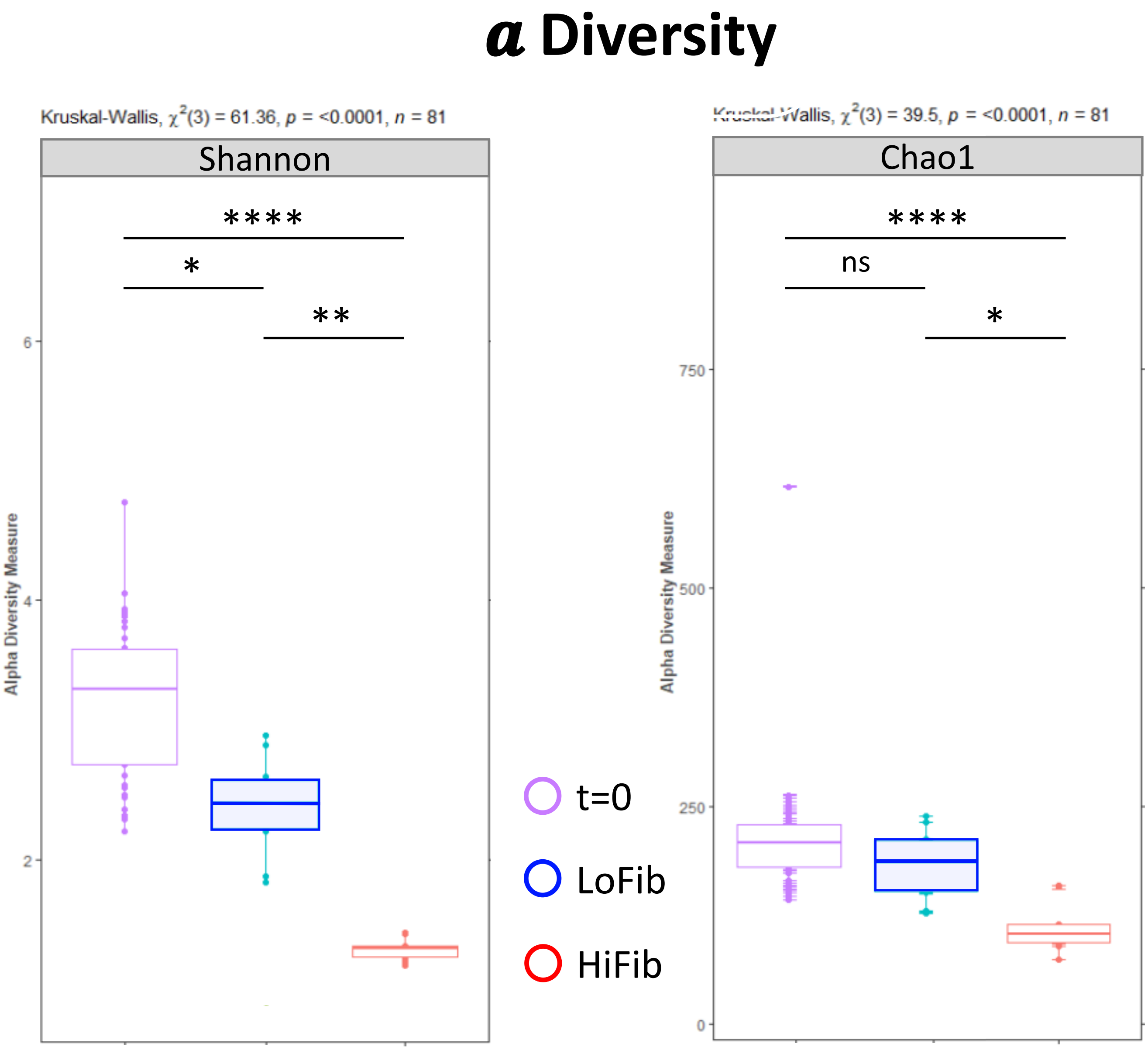

Figure S5: Schematic for HiFib diet (1-week vs 2-week) and LoFib diet

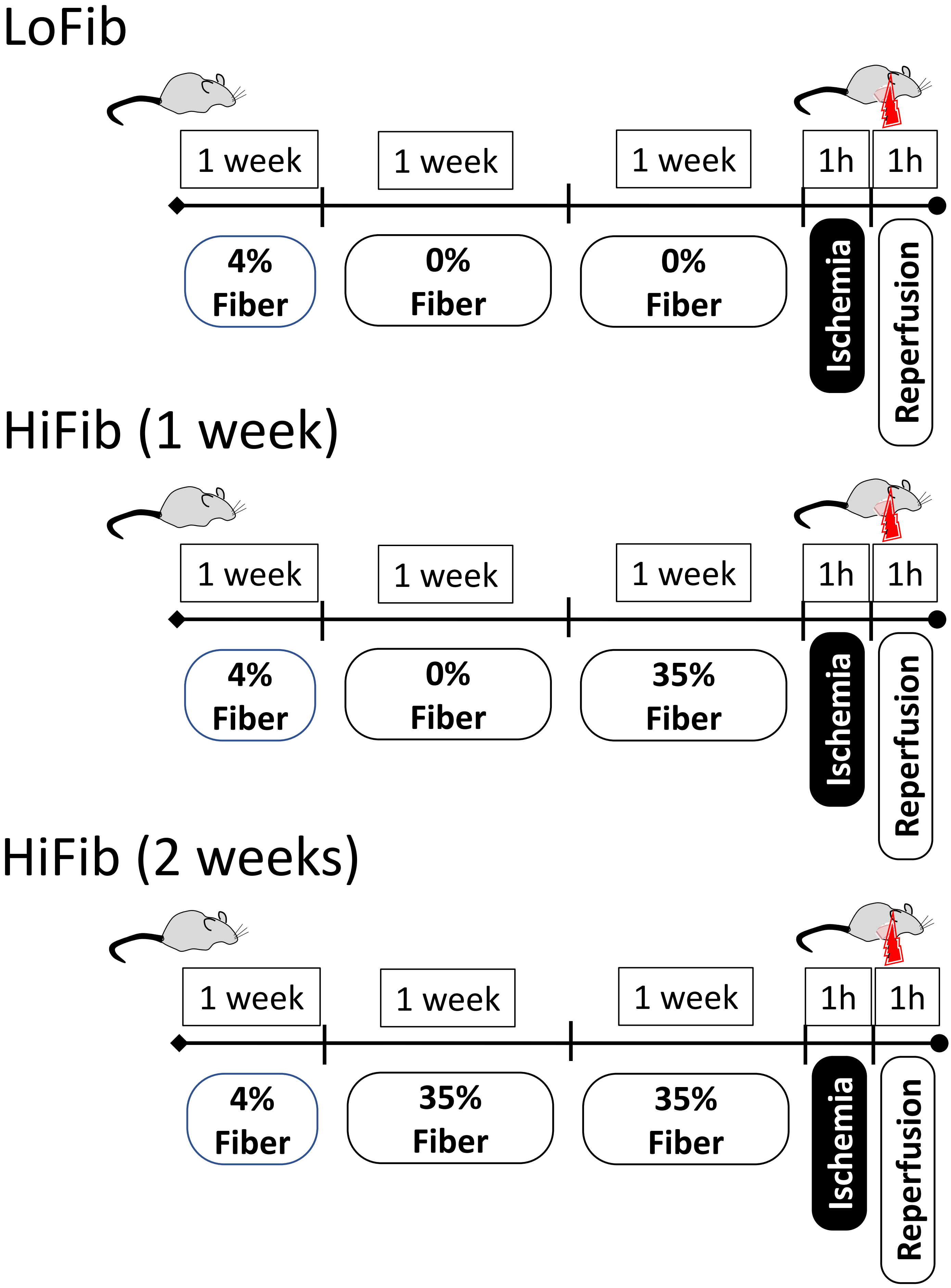

### Figure S6A: Summary of all measurements for all 3 groups

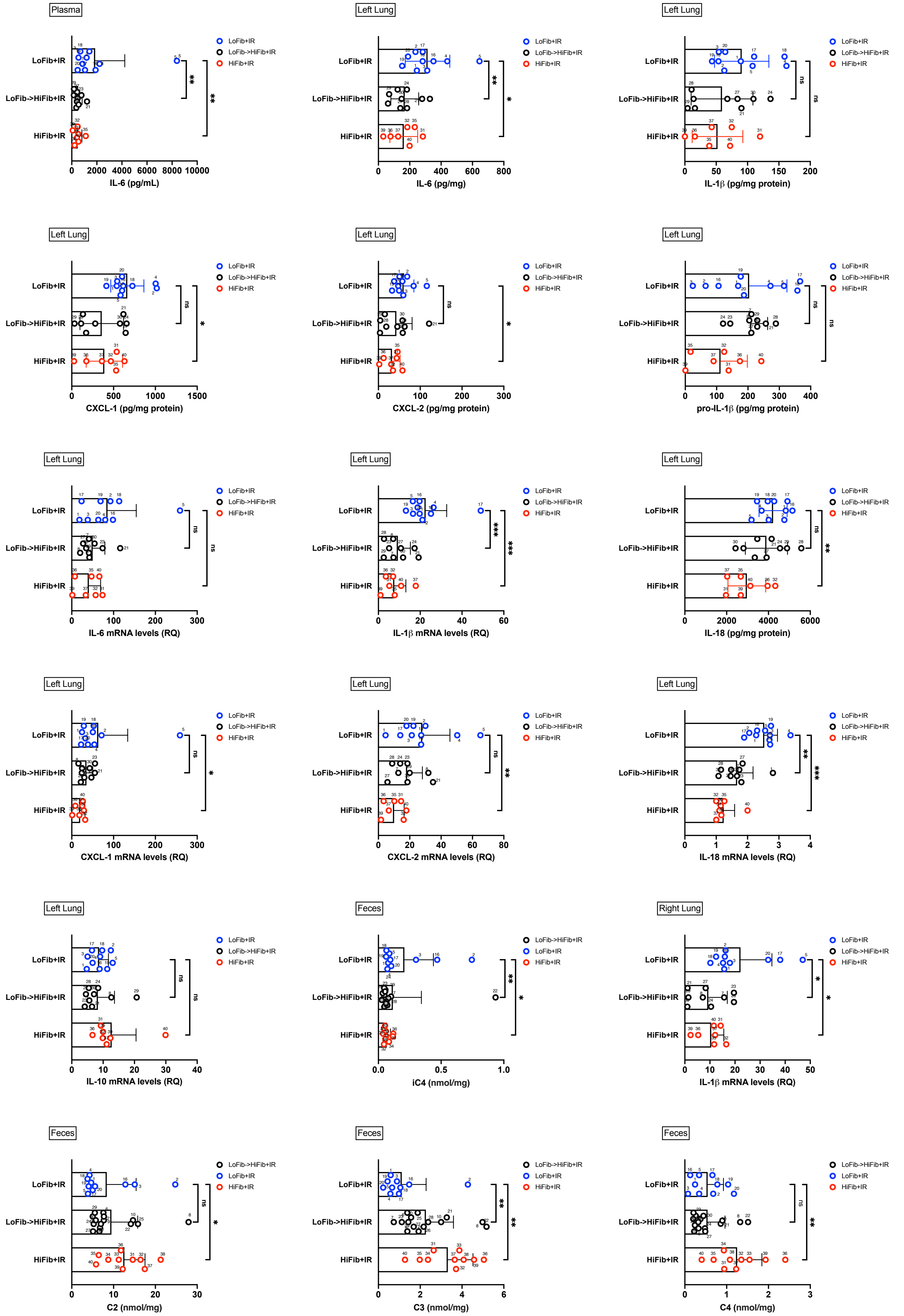

### Figure S6B: Summary of all measurements for all 3 groups

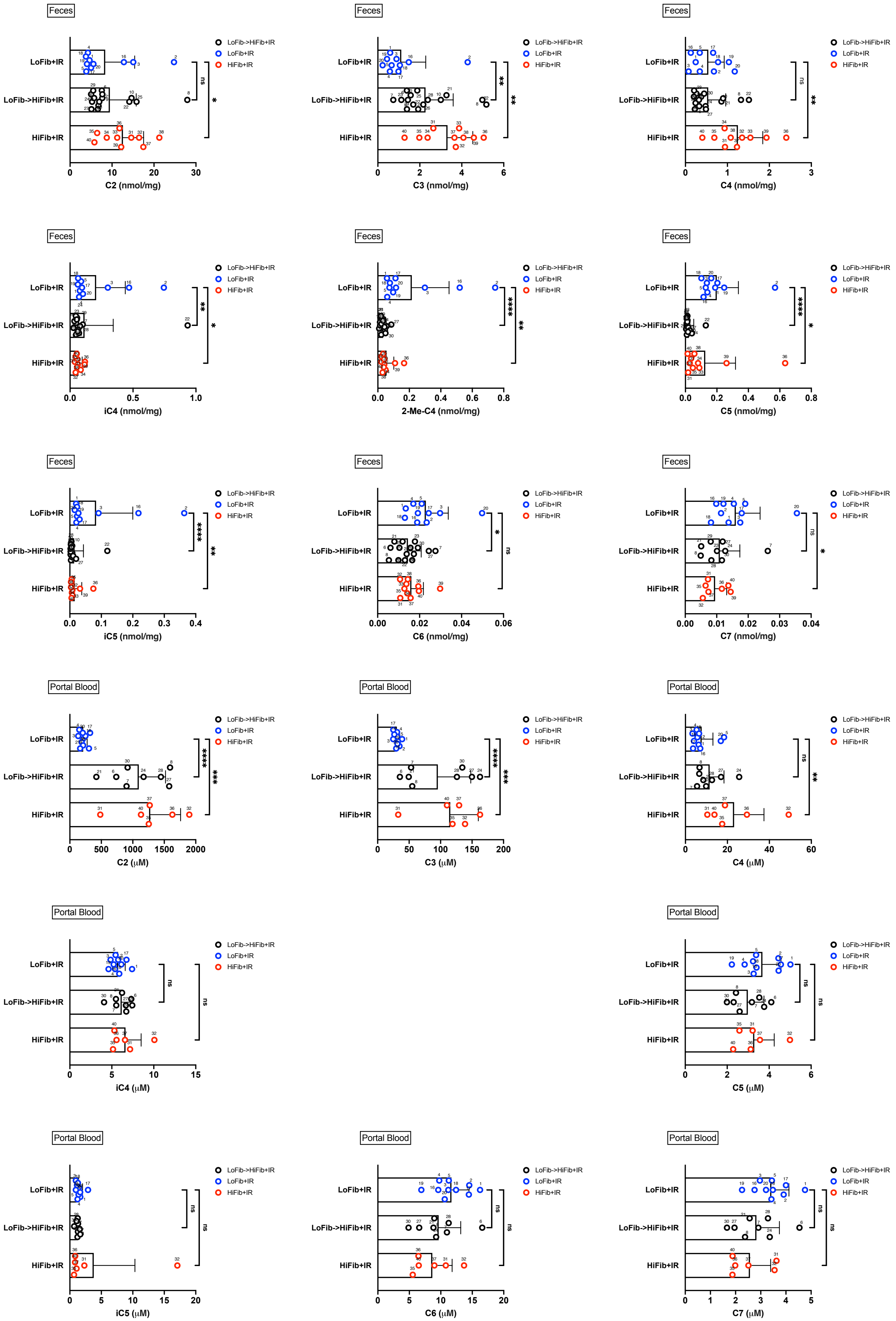

### Figure S6C: Summary of all measurements for all 3 groups

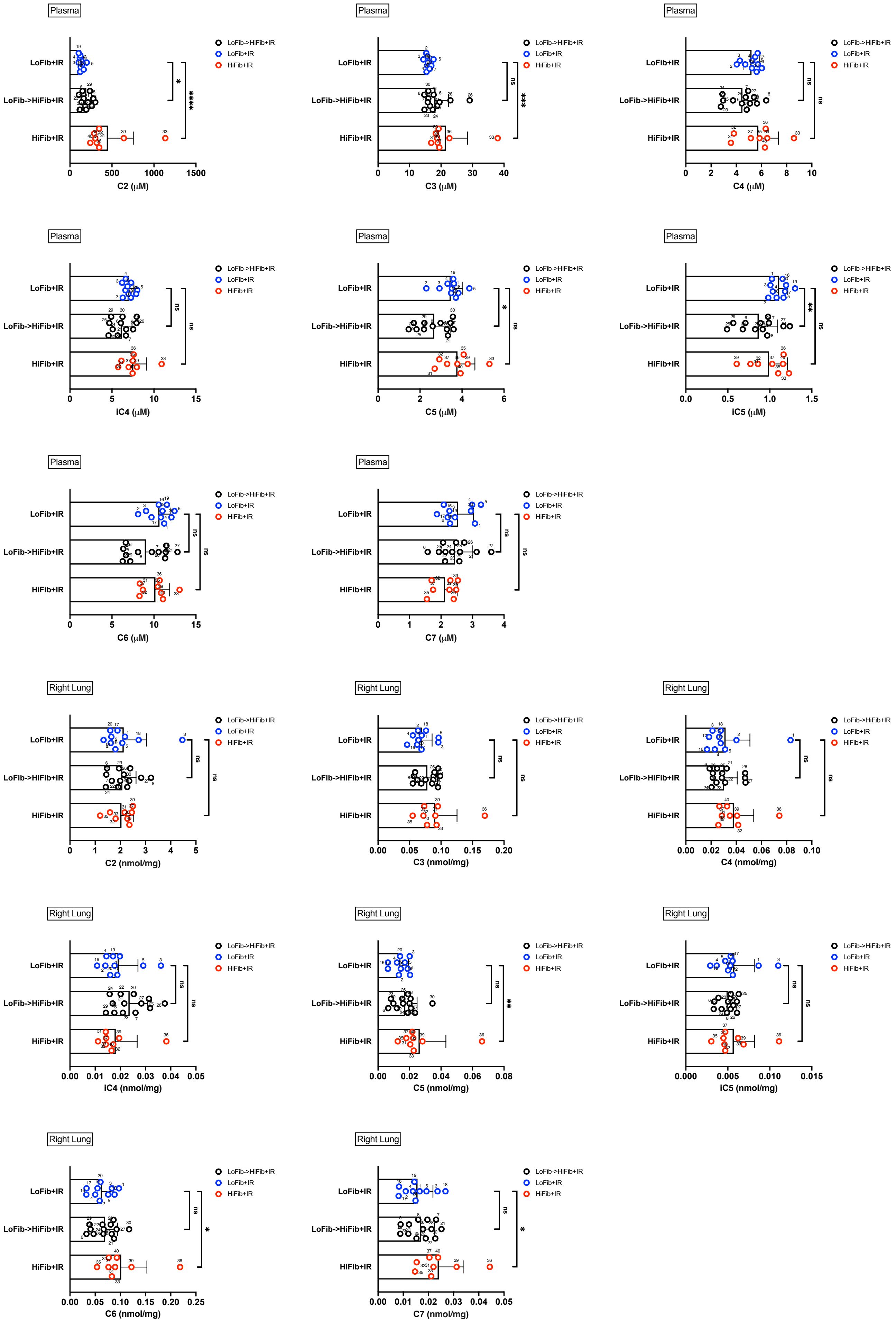

### Figure S7A: PCA plot of HiFib (1 week): Cytokines and Metabolites

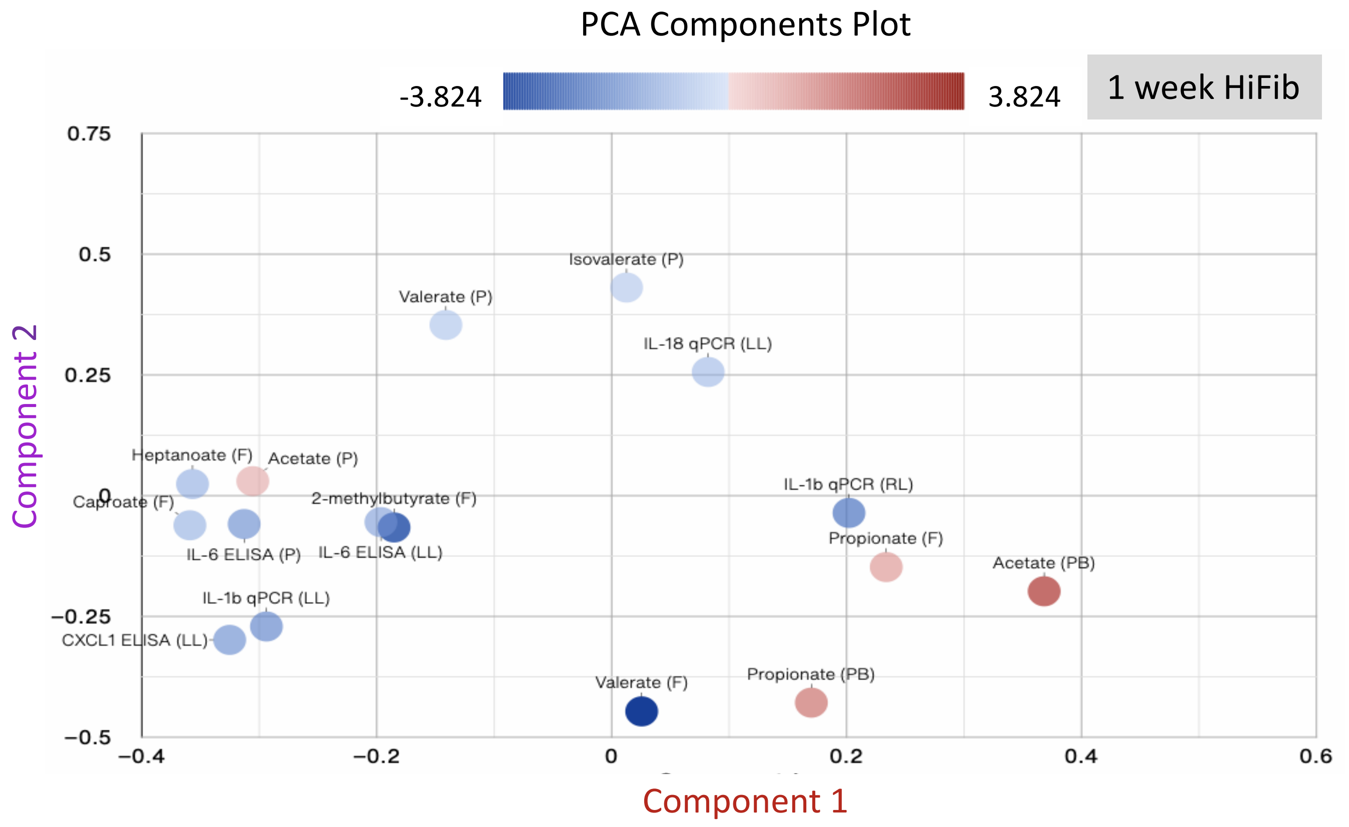


**Component 4**

0.5354 • 2-methylbutyrate (F)
-0.4795 • IL-6 ELISA (LL)
-0.3875 • IL-6 ELISA (P)
0.3107 • Caproate (F)
-0.2885 • Propionate (F)
-0.248 • IL-1β qPCR (LL)
0.1602 • Propionate (PB)
-0.1467 • IL-1β qPCR (RL)
0.1044 • Heptanoate (F)

**Component 3**

-0.4436 • Propionate (F)
-0.4209 • Valerate (F)
-0.3824 • Valerate (P)
-0.3453 • Isovalerate (P)
-0.2495 • IL-1β qPCR (LL)
-0.2443 • CXCL1 ELISA (LL)
-0.2415 • IL-18 qPCR (LL)
0.2372 • IL-6 ELISA (P)
0.2335 • IL-1β qPCR (RL)

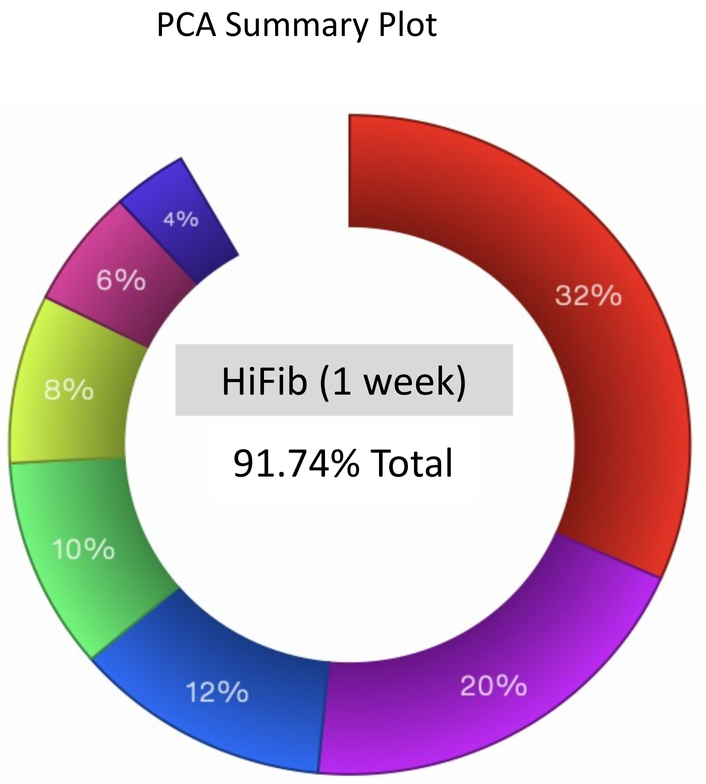


**Component 1**

0.3682 • Acetate (PB)
-0.3591 • Caproate (F)
-0.3568 • Heptanoate (F)
-0.3252 • CXCL1 ELISA (LL)
-0.3129 • IL-6 ELISA (P)
-0.3055 • Acetate (P)
-0.2939 • IL-1β qPCR (LL)
0.2338 • Propionate (F)
0.2021 • IL-1β qPCR (RL)

**Component 2**

-0.4466 • Valerate (F)
0.4309 • Isovalerate (P)
-0.4285 • Propionate (PB)
0.3534 • Valerate (P)
-0.2986 • CXCL1 ELISA (LL)
-0.2708 • IL-1β qPCR (LL)
0.2561 • IL-18 qPCR (LL)
-0.1979 • Acetate (PB)
-0.1478 • Propionate (F)

### Figure S7B: Volcano plot of HiFib (1 week) vs. LoFib : Cytokines and Metabolites

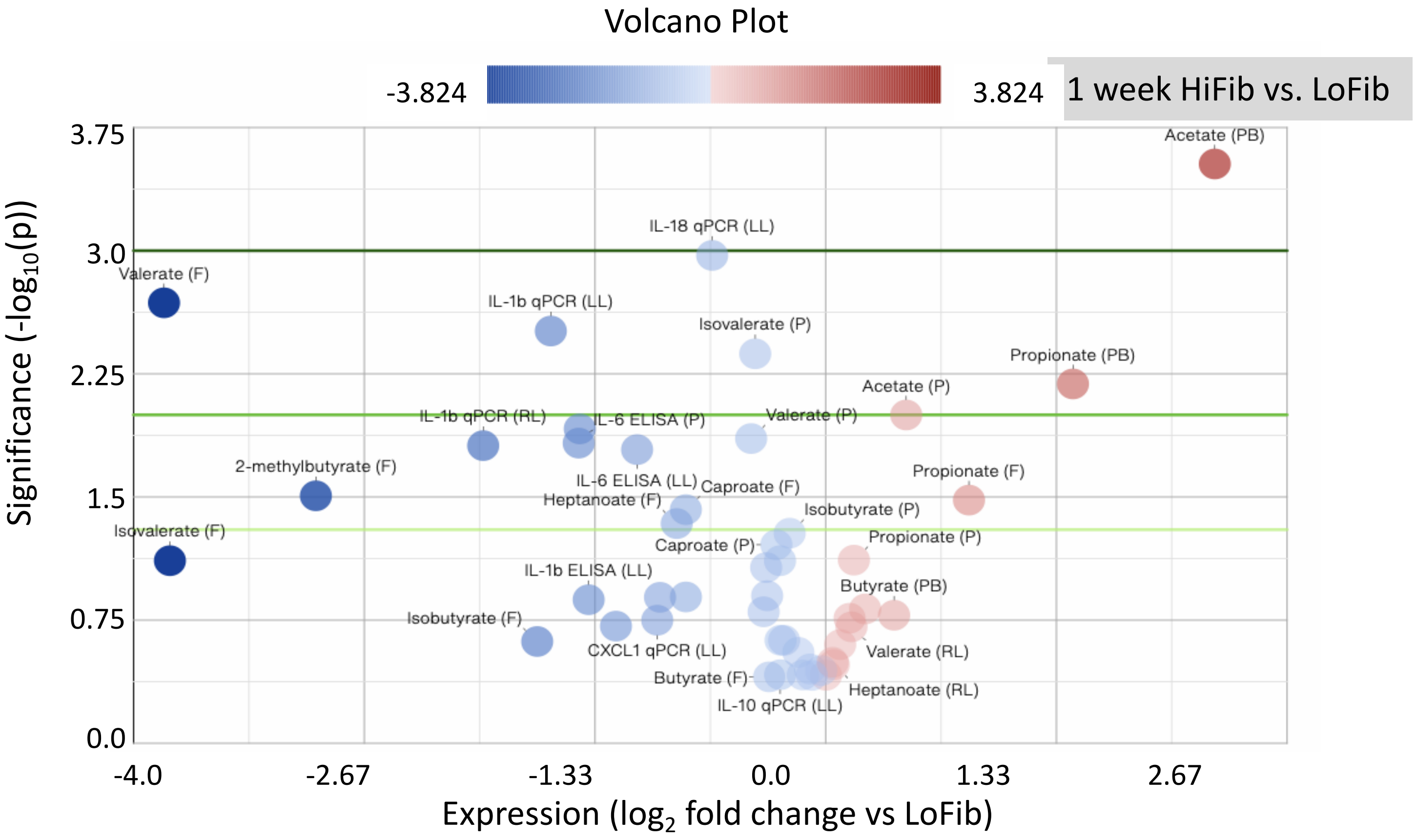

### Supplemental Table 1: Correlation table (all Groups) and including 16S data

Correlation Table (SILVA)

| Symbol | Actinobacteriota | Bacteria_Total | Bacteroidota | Firmicutes | F/B SILVA | OTU_Count | Proteobacteria | Verrucomicrobiota | Caproate (F) | Heptanoate (F) | Propionate (F) | CXCL1 ELISA (LL) | IL-1b ELISA (LL) | IL-1b qPCR (LL) | IL-1B qPCR (RL) | IL-18 qPCR (LL) |
|---|---|---|---|---|---|---|---|---|---|---|---|---|---|---|---|---|
| “Actinobacteriota*” | 0.986*** | −0.379* | −0.694*** | 0.57*** | 0.805*** | | −0.376* | | | | | 0.587** | 0.396* | 0.49* | | 0.475* |
| “Bacteroidota*” | −0.728*** | | 0.901*** | −0.928*** | −0.866*** | | 0.598*** | −0.415* | −0.429* | −0.468** | 0.394* | −0.621*** | −0.442* | −0.646*** | −0.536** | −0.753*** |
| “Firmicutes*” | 0.729*** | | −0.872*** | 0.942*** | 0.95*** | | −0.485** | | 0.375* | 0.451** | −0.358* | 0.612** | 0.441* | 0.518** | 0.462* | 0.716*** |
| “Proteobacteria*” | −0.405* | | 0.636*** | −0.521** | −0.511** | | 0.969*** | −0.523** | −0.412* | | | −0.609** | −0.458* | −0.7*** | | −0.452* |
| “Verrucomicrobiota*” | | 0.371* | | | | 0.479** | −0.521** | 0.987*** | 0.339* | | | | | 0.471* | | |
| FECES | | | | | | | | | | | | | | | | |
| Acetate | | | | | | | | | | | 0.731*** | | | | | |
| Propionate | | | | −0.448** | | −0.418* | | | | −0.496** | 1*** | | −0.4* | | −0.446* | −0.672*** |
| Butyrate | | | | | | | | | | | 0.741*** | | | | | −0.427* |
| Isobutyrate | | | | | | | | | | | 0.339* | | | | | |
| 2-methylbutyrate | 0.377* | | −0.441* | 0.488** | 0.42* | | −0.375* | | | | | 0.404* | | | | |
| Valerate | 0.439* | | −0.467** | 0.384* | 0.408* | | | | 0.349* | | | | | | | |
| Isovalerate | 0.382* | | −0.386* | 0.416* | 0.386* | | −0.345* | | | | | 0.407* | | | | |
| Caproate | | | −0.344* | 0.417* | 0.356* | | | | 1*** | 0.886*** | | | | | 0.395* | |
| Heptanoate | | | −0.34* | 0.502** | 0.411* | | | | 0.886*** | 1*** | −0.496** | | | | 0.549** | 0.573** |
| LEFT LUNG | | | | | | | | | | | | | | | | |
| CXCL1 ELISA | 0.562** | | −0.571** | 0.531** | 0.658*** | | −0.569** | | | | | 1*** | 0.777*** | 0.645*** | | 0.403* |
| CXCL1 qPCR | | | | 0.402* | | | | | | | | | | | 0.584** | |
| CXCL2 ELISA | | | | | | | −0.508** | | | | | 0.75*** | 0.682*** | 0.437* | | |
| CXCL2 qPCR | | | −0.398* | 0.477* | 0.487* | | | | | | | 0.452* | 0.419* | | | 0.526** |
| IL-1b ELISA | 0.413* | | | 0.391* | 0.481* | | −0.418* | | | | −0.4* | 0.777*** | 1*** | 0.593** | | |
| IL-1b qPCR | 0.573** | | −0.607** | 0.537** | 0.497* | | −0.648*** | 0.464* | | | | 0.645*** | 0.593** | | 0.746*** | 0.431* |
| IL-6 ELISA | 0.405* | | −0.461* | 0.558** | 0.445* | | −0.414* | | | | −0.476* | 0.564** | 0.594** | 0.437* | 0.657*** | 0.469* |
| IL-6 qPCR | | | | 0.471* | | | | | | | | 0.416* | 0.428* | | 0.491* | |
| IL-10 qPCR | | | | | | | | | | | | | | | | |
| IL-18 ELISA | | | | | | | | | | | | | | | | |
| IL-18 qPCR | 0.488* | | −0.584** | 0.765*** | 0.68*** | 0.457* | | | | 0.573** | −0.672*** | 0.403* | | 0.431* | 0.398* | 1*** |
| pro-IL-1b ELISA | | | | | | | | | | | | | 0.41* | 0.402* | | |

| Symbol | CXCL1 ELISA (LL) | CXCL1 qPCR (LL) | CXCL2 ELISA (LL) | CXCL2 qPCR (LL) | IL-10 qPCR (LL) | IL-18 ELISA (LL) | IL-18 qPCR (LL) | IL-1b ELISA (LL) | IL-1b qPCR (LL) | IL-1B qPCR (RL) | IL-6 ELISA (LL) | IL-6 qPCR (LL) | pro-IL-1b ELISA (LL) |
|---|---|---|---|---|---|---|---|---|---|---|---|---|---|
| **Actinobacteriota_Enterorhabdus** | | | | | | | 0.508** | | 0.668*** | 0.54** | | | |
| **Bacteroidota_Bacteroides** | −0.565** | | | −0.418* | | | −0.635*** | | −0.634*** | −0.484* | −0.509** | | |
| **Firmicutes_[Eubacterium] nodatum group** | | | | | | | 0.59** | | 0.474* | | | | |
| **Firmicutes_Anaerotruncus** | | 0.473* | | 0.596** | | | 0.542** | | | 0.46* | 0.65*** | 0.485* | |
| **Firmicutes_Clostridium sensu stricto 1** | | | | | | | | | 0.742*** | 0.43* | | | |
| **Firmicutes_Dubosiella** | 0.534** | 0.416* | | 0.47* | | | 0.689*** | 0.431* | | 0.427* | 0.469* | 0.477* | |
| **Firmicutes_Lachnospiraceae NC2004 group** | | | | | | | | | 0.662*** | | | | 0.411* |
| **Firmicutes_Ligilactobacillus** | | 0.77*** | | 0.516* | | | | | | | | 0.611** | |
| **Propionate** | | | | | | | −0.672*** | −0.4* | | −0.446* | −0.476* | | |

### Figure S8A: IL-1$\beta$, IL-18 and Phyla (all Groups)

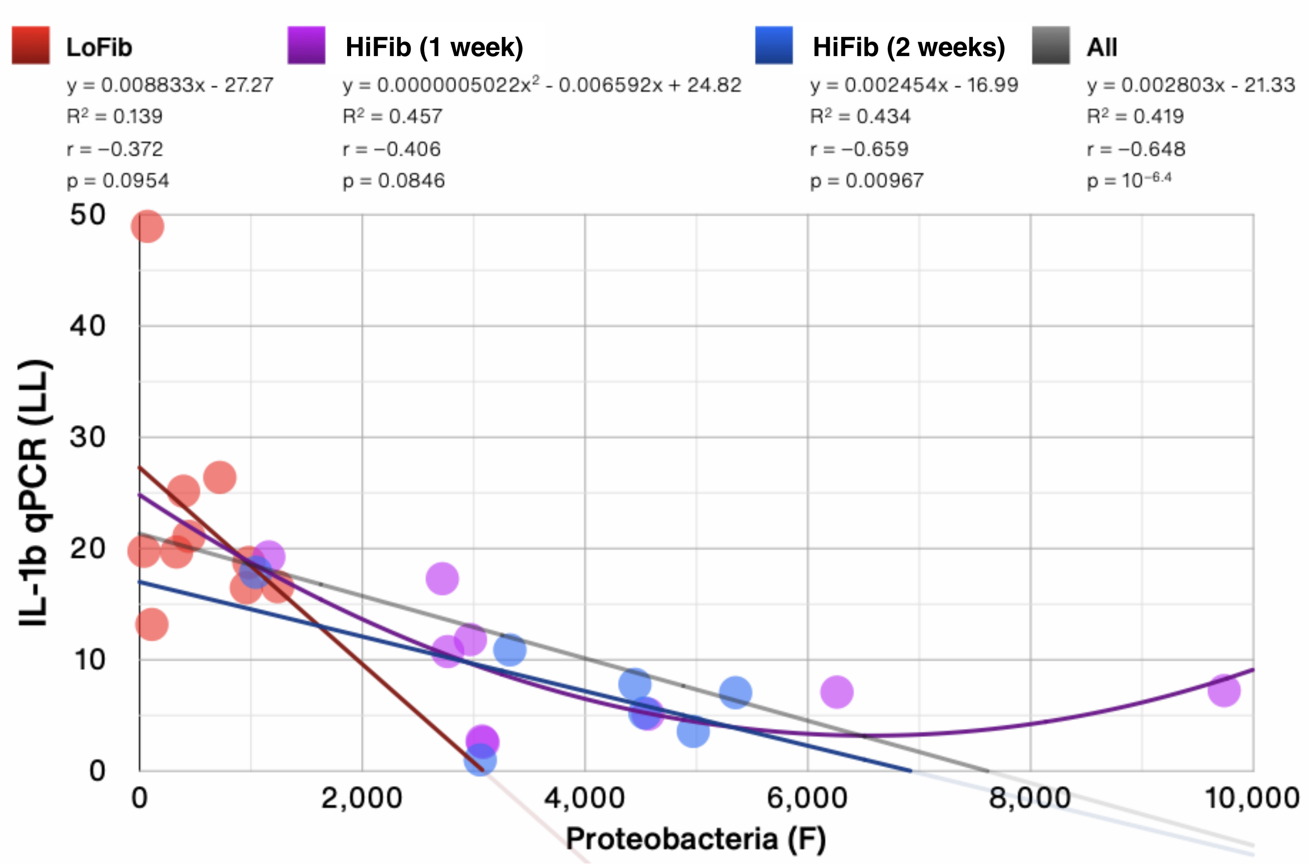


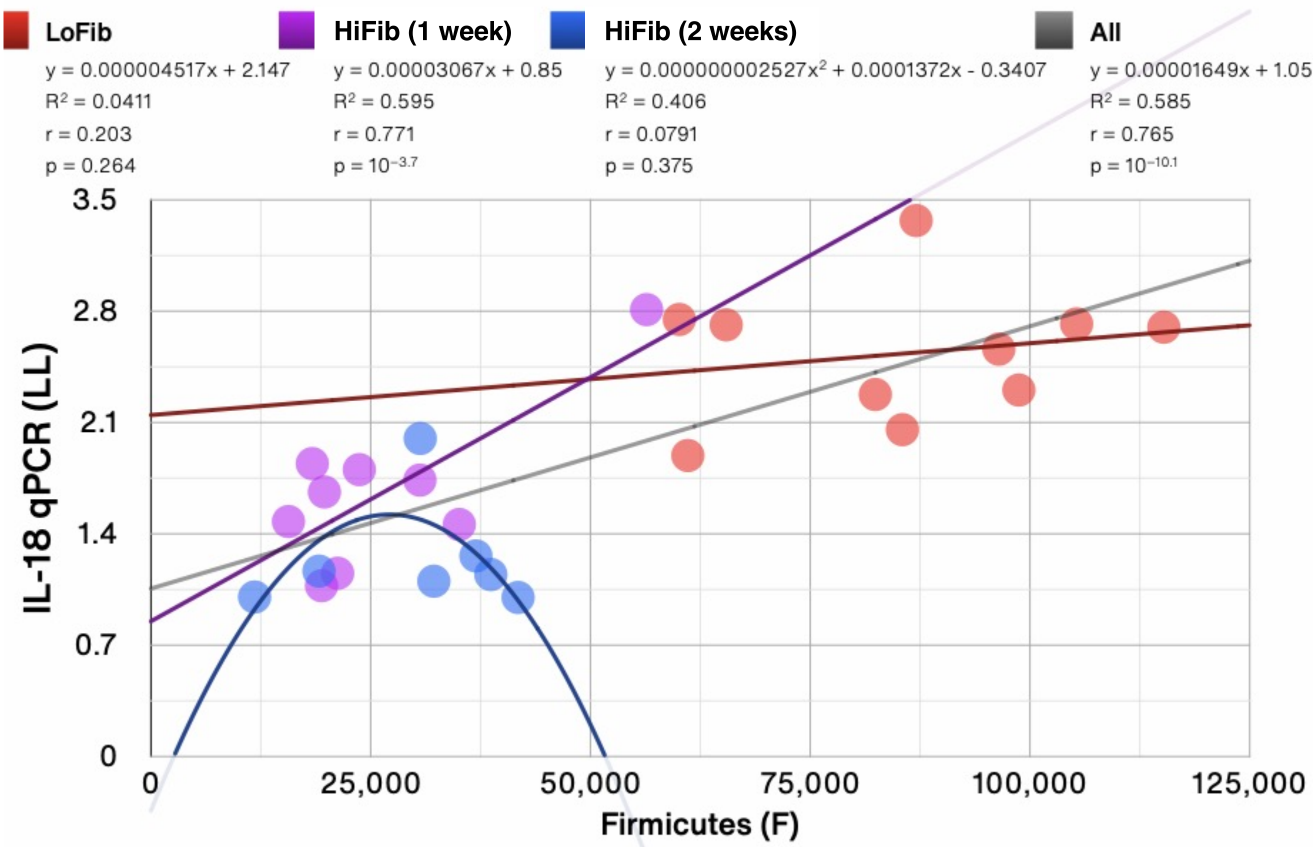

### Figure S8B: More correlations (all Groups)

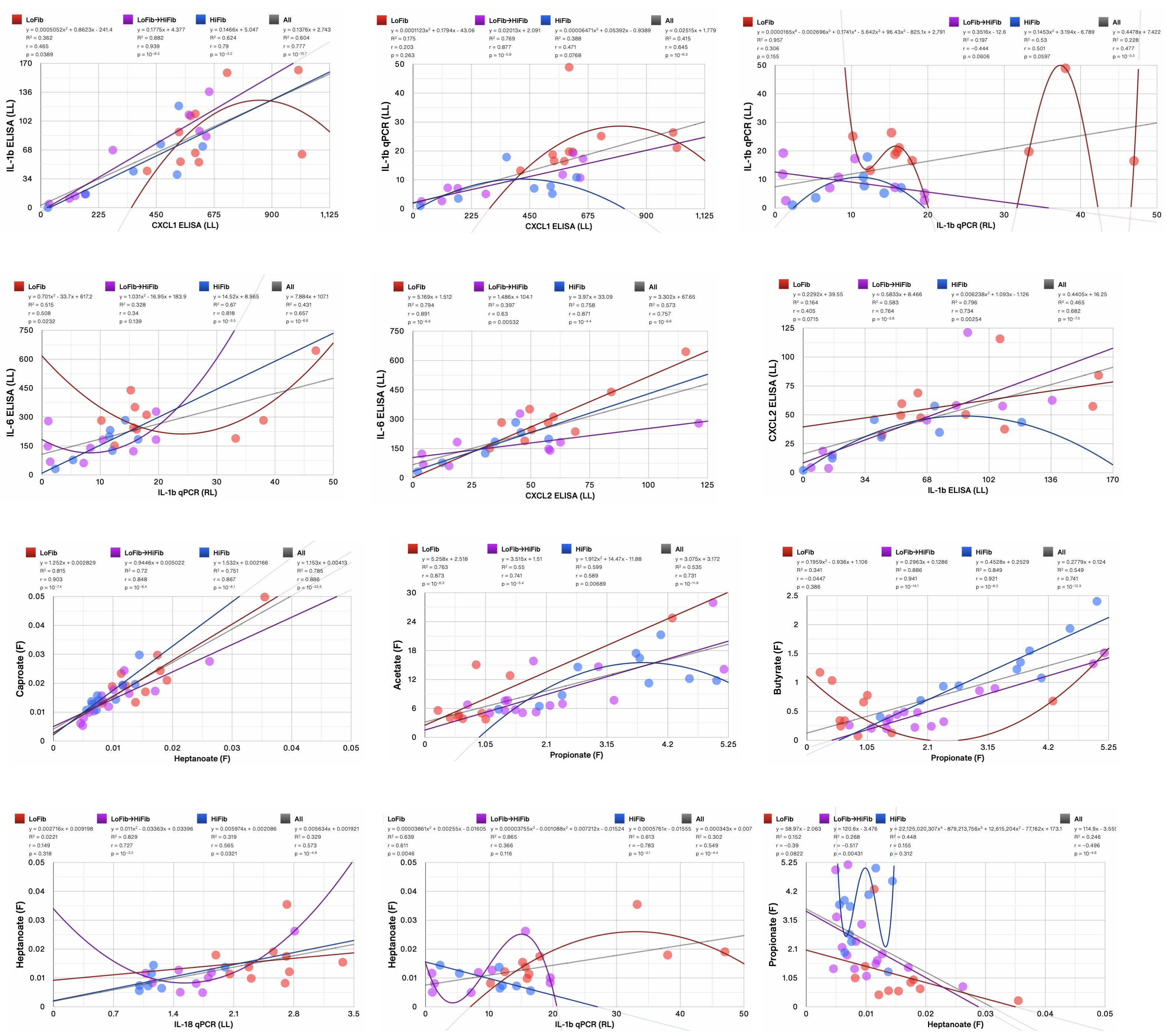

### Supplemental Table 2: ASV level linear correlations with SCFAs and lung immune tone (all groups)

**Anti-inflammatory: Reducing Lung Immune tone ASVs and associated metabolite levels**

| ASV | Fecal C3 | Fecal C7 | Fecal 2Me C4 | Fecal iC5 | Fecal C5 | Portal Blood C2 | Portal Blood C3 | Portal Blood C4 | LL IL-1b mRNA | LL IL-18 mRNA | LL IL-6 mRNA | LL IL-6 ELISA | Plasma IL-6 | P value |
|---|---|---|---|---|---|---|---|---|---|---|---|---|---|---|
| 811f358e4e89eecbd75298ffce878cba_Lachnospiraceae_NA_NA | ▲ (red) | | | | | ▲ (green) | ▲ (red) | ▲ (black) | ▼ (black) | ▼ (red) | | | | 0.001 ▲ (green)<br>0.01 ▲ (red)<br>0.05 ▲ (black) |
| f09f257605135a6049b84 7768387fb33 _ Lachnospiraceae _NA_ NA | ▲ (red) | | | | | ▲ (black) | ▲ (black) | ▲ (black) | | ▼ (red) | | | | 0.01 ▲ (red)<br>0.05 ▲ (black) |
| f54447ae705d557e2c2b3777fbdd56fO_Lachnospiraceae_NA_NA | | | | | | ▲ (red) | ▲ (green) | ▲ (red) | | ▼ (red) | | | | 0.001 ▲ (green)<br>0.01 ▲ (red) |
| 3fb663a5bf98e 163dbeedefe91 c6bae9 _ Lachnospiraceae _NA_ NA | | | | | | ▲ (red) | ▲ (black) | | ▼ (red) | | | | | 0.01 ▲ (red)<br>0.05 ▲ (black) |
| 69a 11 f927915e2a3ba 1 f7b9c84486527 _ Sutterellaceae _Parasutterella _ NA | ▲ (red) | | | | | ▲ (red) | ▲ (red) | | ▼ (black) | ▼ (black) | | | | 0.01 ▲ (red)<br>0.05 ▲ (black) |
| 836cd5ea357 ad392c00a 7 e48d921 be1 d _ Bacteroidaceae _ Bacteroides _ NA | ▲ (red) | | | | | ▲ (red) | ▲ (red) | ▲ (black) | | ▼ (black) | | | | 0.01 ▲ (red)<br>0.05 ▲ (black) |

**Pro-inflammatory: Increasing Lung Immune tone ASVs and associated metabolite levels**

| ASV | Fecal C3 | Fecal C7 | Fecal 2Me C4 | Fecal iC5 | Fecal C5 | Portal Blood C2 | Portal Blood C3 | Portal Blood C4 | LL IL-1b mRNA | LL IL-18 mRNA | RL IL-1b mRNA | LL IL-6 ELISA | Plasma IL-6 | P value ▼ (green) ▼ (red) ▼ (black) |
|---|---|---|---|---|---|---|---|---|---|---|---|---|---|---|
| 22907d450b79c927e4ffe34a2d09d2aa_Lachnospiraceae_Lachnoclostridium_NA | ▼ (red) | ▲ (black) | ▲ (red) | ▲ (red) | | ▼ (red) | ▼ (green) | | ▲ (black) | ▲ (red) | | ▲ (red) | ▲ (black) | 0.001 ▲ (green)<br>0.01 ▲ (red)<br>0.05 ▲ (black) |
| d114fb4c335125128be28401522dd41a_Streptococcaceae_Lactococcus_NA | ▼ (green) | ▲ (red) | ▲ (red) | | | ▼ (red) | ▼ (red) | ▼ (black) | | ▲ (red) | | | | 0.001 ▲ (green)<br>0.01 ▲ (red)<br>0.05 ▲ (black) |
| f3e9d78daeea42d345 0807 48e31 ae3dd_ Oscillospiraceae _ Colidextribacter_NA | | | | | | ▼ (red) | ▼ (green) | ▼ (black) | | ▲ (red) | | | | 0.001 ▲ (green)<br>0.01 ▲ (red)<br>0.05 ▲ (black) |
| 0f2a5acc53e55f2c9fOc7409421dc5a0_Ruminococcaceae_Anaerotruncus_uncultured_bacterium | | | ▲ (black) | ▲ (red) | ▲ (red) | | ▼ (red) | ▼ (red) | | ▲ (red) | ▲ (black) | | | 0.01 ▲ (red)<br>0.05 ▲ (black) |
| 45b 76e49b6500d8d245c2 f27f6fbb 143 _ Erysipelatoclostridiaceae _ Erysipelatoclostridium _ NA | | | | | | | ▼ (black) | ▼ (red) | | ▲ (red) | | | | 0.01 ▲ (red)<br>0.05 ▲ (black) |
| 029ac1 f87 abeae 7 dbac1 ababc489cfec _ Muribaculaceae _ Muribaculaceae _ NA | | | ▲ (green) | ▲ (green) | ▲ (green) | | | | ▲ (black) | | | ▲ (red) | ▲ (red) | 0.001 ▲ (green)<br>0.01 ▲ (red)<br>0.05 ▲ (black) |
| e335f7 4033bc634af43ee6ba a84fa24 7 _Peptostreptococcaceae_Romboutsia_NA | | | | ▲ (black) | | ▼ (black) | ▼ (black) | ▼ (black) | | ▲ (black) | | | | 0.05 ▲ (black) |

### Figure S9: Targeted PICRUST Metabolic Pathway Comparison between Groups 1-3

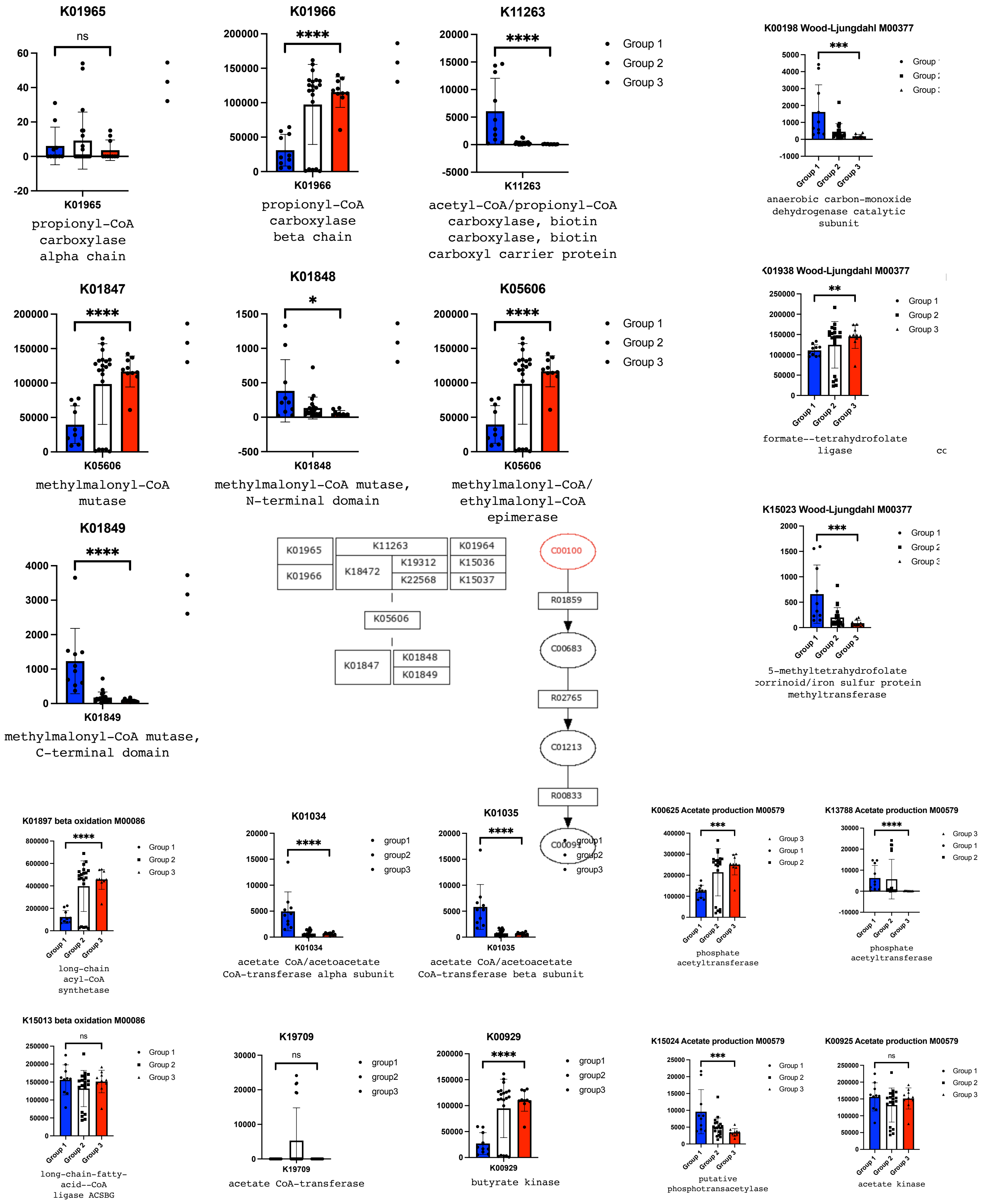

Supplemental Figure S10: Metabolic reprogramming of AM & AT2 cells in vitro after C3 exposure

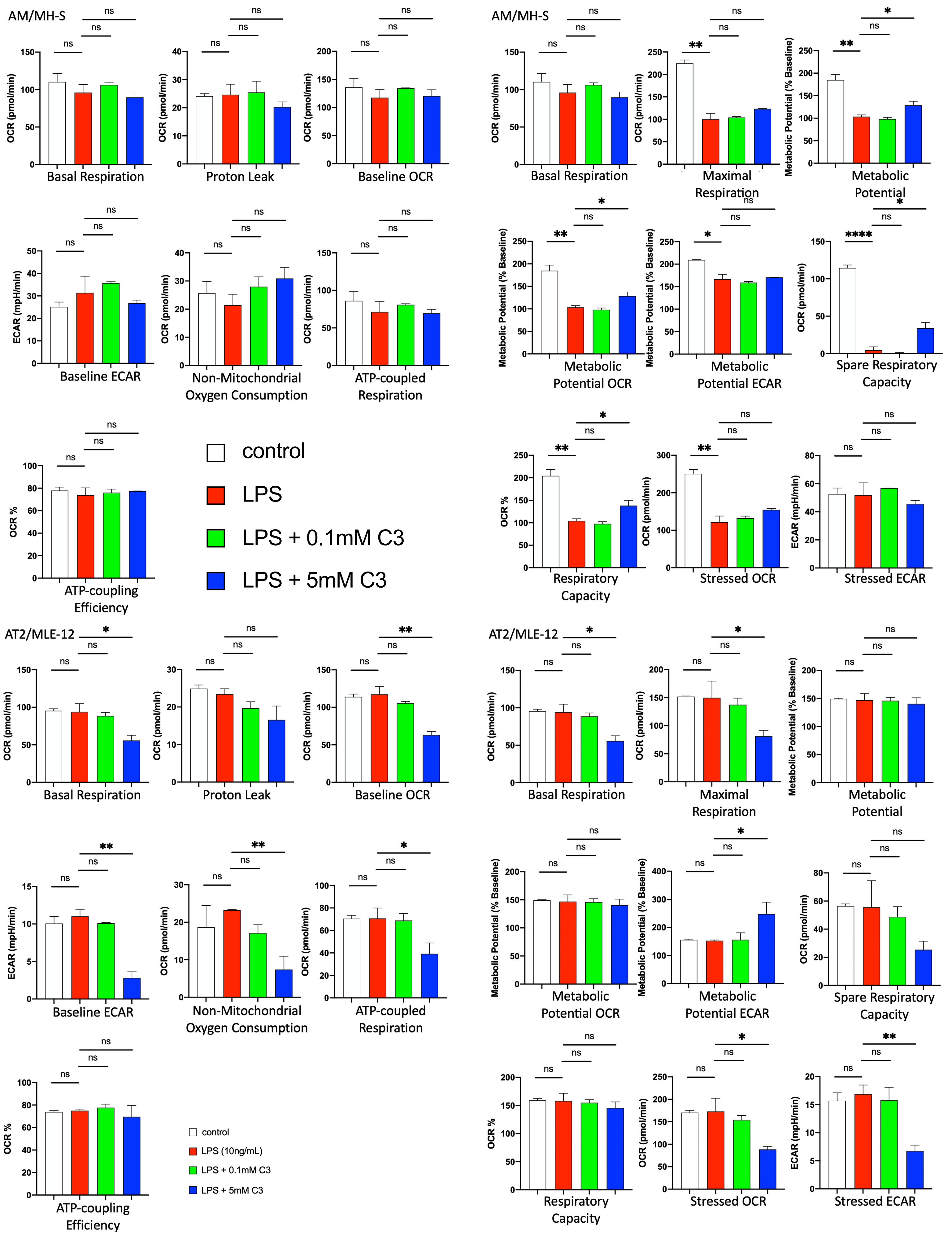

Supplemental Figure S11A: Difference in stool, colon morphology, and weights between LoFib and HiFib groups

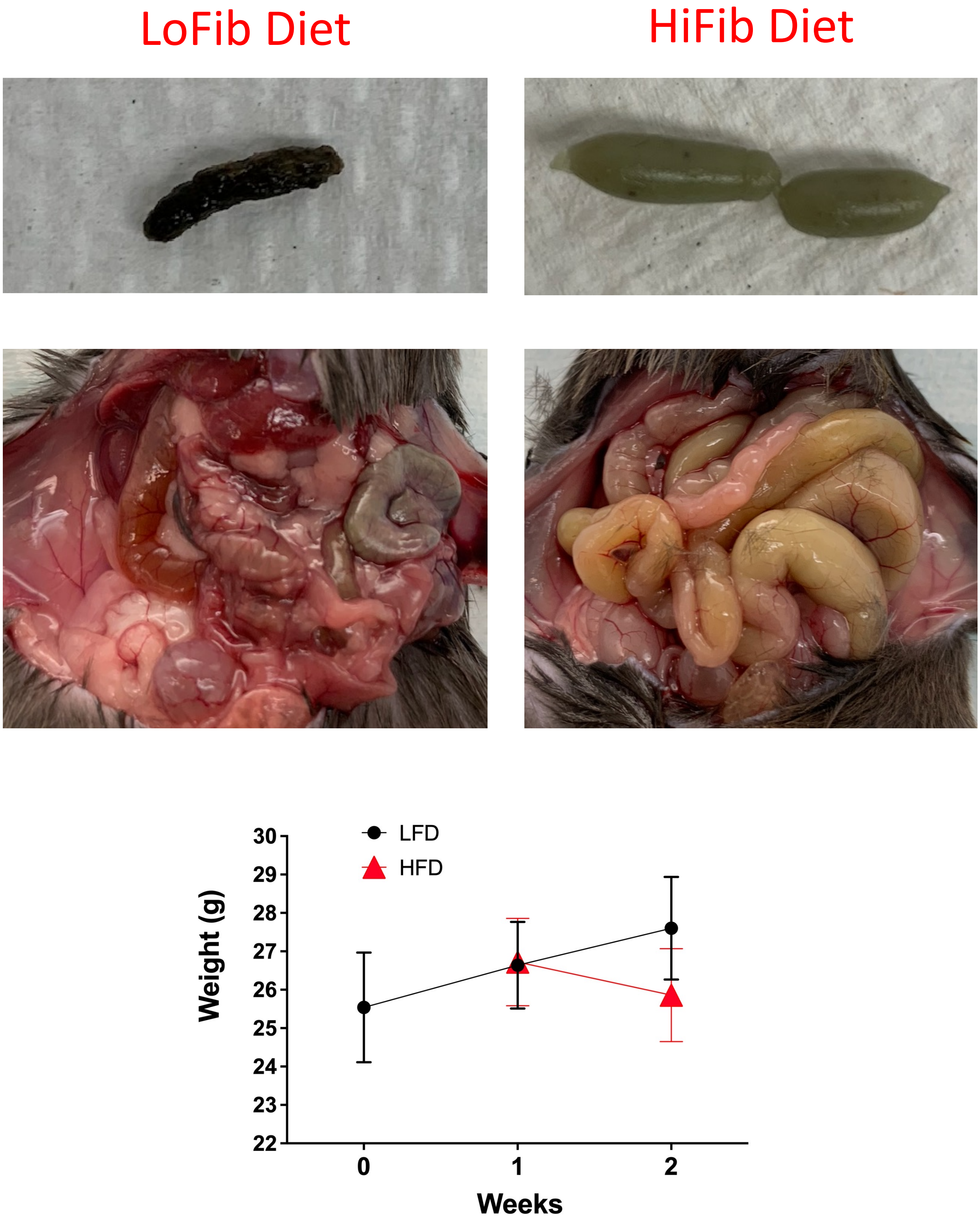

### Supplemental Figure S11B: t=0 starting diet

### LARC diet-5053 4.4% fiber

#### DESCRIPTION

PicoLab® Rodent Diet 20 is a 20% protein diet formulated for rat, hamster and mouse breeding colonies. This diet is a complete life cycle diet formulated using managed formulation, delivering Constant Nutrition®. This is paired with the selection of highest quality ingredients to assure minimal inherent biological variation in long-term studies. Irradiation gives reliable microbial control and eliminates the need for autoclaving. Irradiation treatment and special 3-ply packaging provide virtually bacteria-free dietary control.

##### Features and Benefits

- Managed Formulation delivers Constant Nutrition®
- Precision processing and selection of highest quality ingredients assures Constant Nutrition® quality
- Formulated with 20% protein
- High quality animal protein added to create a superior balance of amino acids for optimum performance
- Recommended for rat breeding colonies and mice not requiring a higher energy diet
- Irradiation gives reliable microbial control and eliminates the need for autoclaving

| Product Forms Available | Catalog # |
|---|---|
| • Oval pellet, (3/8"x5/8"x1"), Irradiated, 30 lb | 3005740-220 |
| • Meal (ground pellets), Irradiated, 30 lb | 3005740-020 |
| **Other Irradiated Versions Available** | **Catalog #** |
| • 5R53: Extruded Particle, Irradiated, 20lb | 3002890-712 |
| • 5R53: Meal, Irradiated, 30lb | 3005839-020 |
| • 5061: Pico-Vac® Lab Rodent Diet, Irradiated 5 lb vacuum sealed, 6 per box (30 lb box) | 0006954 |
| • 5K75: Certified PicoLab® Rodent 20, Irradiated, 30 lb | 3005965-220 |
| • 5LU7: Macro-Pack ™ PicoLab® Rodent Diet 5053 | 0066400 |
| **Other Non-Irradiated Versions Available** | **Catalog #** |
| • 5L0B: Laboratory Rodent Diet 20, 15 kg | 0067097 |
| • 5RA3: Lab Rodent 20 Auto Ext, 25 lb | 3006933-703 |

#### GUARANTEED ANALYSIS

| | |
|---|---|
| Crude protein not less than | 20.00% |
| Crude fat not less than | 4.50% |
| Crude fiber not more than | 6.00% |
| Ash not more than | 7.00% |
| Moisture not more than | 12.00% |

#### INGREDIENTS

Ground Corn, Dehulled Soybean Meal, Wheat Middlings, Ground Wheat, Fish Meal, Wheat Germ, Dried Plain Beet Pulp, Cane Molasses, Brewers Dried Yeast, Ground Oats, Dehydrated Alfalfa Meal, Soybean Oil, Dried Whey, Calcium Carbonate, Salt, DL-Methionine, Menadione Dimethylpyrimidinol Bisulfite (Vitamin K), Choline Chloride, Pyridoxine Hydrochloride, Cholecalciferol (Vitamin D3), Vitamin A Acetate, DL-Alpha-Tocopherol Acetate (Vitamin E), Biotin, Folic Acid, Thiamine Mononitrate, Manganous Oxide, Vitamin B12 Supplement, Zinc Oxide, Ferrous Carbonate, Nicotinic Acid, Riboflavin Supplement, Calcium Pantothenate, Copper Sulfate, Zinc Sulfate, Calcium Iodate, Cobalt Carbonate, Sodium Selenite.

#### CHEMICAL COMPOSITION[1]

| Nutrients[2] | |
|---|---|
| **Protein, %** | **21.0** |
| Arginine, % | 1.29 |
| Cystine, % | 0.36 |
| Glycine, % | 0.98 |
| Histidine, % | 0.53 |
| Isoleucine, % | 0.87 |
| Leucine, % | 1.58 |
| Lysine, % | 1.18 |
| Methionine, % | 0.62 |
| Phenylalanine, % | 0.92 |
| Tyrosine, % | 0.61 |
| Threonine, % | 0.79 |
| Tryptophan, % | 0.24 |
| Valine, % | 0.97 |
| Serine, % | 1.00 |
| Aspartic Acid, % | 2.23 |
| Glutamic Acid, % | 4.26 |
| Alanine, % | 1.20 |
| Proline, % | 1.32 |
| Taurine, % | 0.03 |
| **Fat (ether extract), %** | **5.0** |
| **Fat (acid hydrolysis), %** | **6.3** |
| Cholesterol, ppm | 135 |
| Linoleic Acid, % | 2.32 |
| Linolenic Acid, % | 0.28 |
| Arachidonic Acid, % | 0.02 |
| Omega-3 Fatty Acids, % | 0.42 |
| Total Saturated Fatty Acids, % | 0.77 |
| Total Monounsaturated Fatty Acids, % | 1.00 |
| **Fiber (Crude), %** | **4.4** |
| Neutral Detergent Fiber[3], % | 15.5 |
| Acid Detergent Fiber[4], % | 5.6 |
| **Nitrogen-Free Extract (by difference), %** | **53.5** |
| Starch, % | 28.2 |
| Sucrose, % | 2.71 |
| **Total Digestible Nutrients,%** | **75.1** |
| **Gross Energy, kcal/gm** | **4.11** |
| **Physiological Fuel Value[5], kcal/gm** | **3.43** |
| **Metabolizable Energy, kcal/gm** | **3.02** |

| | |
|---|---|
| Chloride, % | 0.53 |
| Fluorine, ppm | 9.2 |
| Iron, ppm | 184 |
| Zinc, ppm | 79 |
| Manganese, ppm | 82 |
| Copper, ppm | 13 |
| Cobalt, ppm | 0.72 |
| Iodine, ppm | 0.97 |
| Chromium (added), ppm | 0.01 |
| Selenium, ppm | 0.37 |

| Vitamins | |
|---|---|
| Carotene, ppm | 1.5 |
| Vitamin K, ppm | 3.3 |
| Thiamin, ppm | 16 |
| Riboflavin, ppm | 8.1 |
| Niacin, ppm | 84 |
| Pantothenic Acid, ppm | 17 |
| Choline, ppm | 1575 |
| Folic Acid, ppm | 3.0 |
| Pyridoxine, ppm | 9.6 |
| Biotin, ppm | 0.30 |
| $B_{12}$, mcg/kg | 51 |
| Vitamin A, IU/gm | 15 |
| Vitamin $D_3$ (added), IU/gm | 2.3 |
| Vitamin E, IU/kg | 99 |
| Ascorbic Acid, mg/gm | 0.00 |

| Calories provided by: | |
|---|---|
| Protein, % | 24.495 |
| Fat (ether extract), % | 13.122 |
| Carbohydrates, % | 62.382 |

1. Formulation based on calculated values from the latest ingredient analysis information. Since nutrient composition of natural ingredients varies and some nutrient loss will occur due to manufacturing processes, analysis will differ accordingly.
2. Nutrients expressed as percent of ration except where otherwise indicated. Moisture content is assumed to be 10.0% for the purpose of [illegible]

### Supplemental Figure S11C: LoFib diet

### Low fiber diet-5GCX 0% fiber

#### DESCRIPTION

Modification of TestDiet® AIN-93 G Growth Purified Diet, 57W5, with cellulose removed.

Intended for rodents in a laboratory setting.

CAUTION: Contains a new animal drug for investigational use only in laboratory research animals or for tests in vitro. Not for use in humans.

Storage conditions are particularly critical to TestDiet® products, due to the absence of antioxidants or preservative agents. To provide maximum protection against possible changes during storage, store in a dry, cool location. Storage under refrigeration (2° C) is recommended. Maximum shelf life is six months. (If long term studies are involved, storing the diet at -20° C or colder may prolong shelf life.) Be certain to keep in air tight containers.

| ***Product Forms Available**** | ***Catalog #*** |
|---|---|
| 1/2" Pellet, Irradiated | 1819514-203 |

*Other Forms Available On Request*

#### INGREDIENTS (%)

| | |
|---|---|
| Corn Starch | 44.7486 |
| Casein - Vitamin Tested | 20.0000 |
| Maltodextrin | 13.2000 |
| Sucrose | 10.0000 |
| Soybean Oil | 7.0000 |
| AIN 93G Mineral Mix | 3.5000 |
| AIN 93 Vitamin Mix | 1.0000 |
| L-Cystine | 0.3000 |
| Choline Bitartrate | 0.2500 |
| t-Butylhydroquinone | 0.0014 |

#### NUTRITIONAL PROFILE [1]

| ***Protein, %*** | ***18.3*** |
|---|---|
| Arginine, % | 0.70 |
| Histidine, % | 0.52 |
| Isoleucine, % | 0.96 |
| Leucine, % | 1.73 |
| Lysine, % | 1.45 |
| Methionine, % | 0.52 |
| Cystine, % | 0.37 |
| Phenylalanine, % | 0.96 |
| Tyrosine, % | 1.01 |
| Threonine, % | 0.77 |
| Tryptophan, % | 0.22 |
| Valine, % | 1.14 |
| Alanine, % | 0.55 |
| Aspartic Acid, % | 1.29 |
| Glutamic Acid, % | 4.08 |
| Glycine, % | 0.39 |
| Proline, % | 2.36 |
| Serine, % | 1.10 |
| Taurine, % | 0.00 |
| ***Fat, %*** | ***7.1*** |
| Cholesterol, ppm | 0 |
| Linoleic Acid, % | 3.58 |
| Linolenic Acid, % | 0.55 |
| Arachidonic Acid, % | 0.00 |
| Omega-3 Fatty Acids, % | 0.55 |
| Total Saturated Fatty A | 1.05 |
| Total Monounsaturated Fatty Acids, % | 1.54 |
| Polyunsaturated Fatty Acids, % | 3.78 |
| ***Fiber (max), %*** | ***0.0*** |
| ***Carbohydrates, %*** | ***68.2*** |
| ***Energy (kcal/g)*** [2] | ***4.10*** |

| From: | kcal | % |
|---|---|---|
| Protein | 0.733 | 17.9 |
| Fat (ether extract) | 0.638 | 15.6 |
| Carbohydrates | 2.728 | 66.6 |

| ***Minerals*** | |
|---|---|
| Calcium, % | 0.51 |
| Phosphorus, % | 0.32 |
| Potassium, % | 0.36 |
| Magnesium, % | 0.05 |
| Sodium, % | 0.13 |
| Chloride, % | 0.22 |
| Fluorine, ppm | 1.0 |
| Iron, ppm | 39 |
| Zinc, ppm | 35 |
| Manganese, ppm | 11 |
| Copper, ppm | 6.0 |
| Cobalt, ppm | 0.0 |
| Iodine, ppm | 0.21 |
| Chromium (added), ppm | 1.0 |
| Molybdenum, ppm | 0.14 |
| Selenium, ppm | 0.24 |

| ***Vitamins*** | |
|---|---|
| Vitamin A, IU/g | 4.0 |
| Vitamin D-3 (added), IU/g | 1.0 |
| Vitamin E, IU/kg | 81.6 |
| Vitamin K, ppm | 0.75 |
| Thiamin, ppm | 4.8 |
| Riboflavin, ppm | 6.7 |
| Niacin, ppm | 30 |
| Pantothenic Acid, ppm | 16 |
| Folic Acid, ppm | 2.1 |
| Pyridoxine, ppm | 5.8 |
| Biotin, ppm | 0.2 |
| Vitamin B-12, mcg/kg | 28 |
| Choline Chloride, ppm | 1,250 |
| Ascorbic Acid, ppm | 0.0 |

1. Formulation based on calculated values from the latest ingredient analysis information. Since nutrient composition of natural ingredients varies and some nutrient loss will occur due to manufacturing processes, analysis will differ accordingly. Nutrients expressed as percent of ration on an As-Fed basis except where otherwise indicated.

2. Energy (kcal/gm) - Sum of decimal [illegible]

### Supplemental Figure S11D: HiFib diet

### High Fiber diet-5Z6R 35% fiber

#### Mod TestDiet® 57W5 w/ 30% Pectin, Blue — 5Z6R

##### DESCRIPTION

Modification of TestDiet® AIN-93G Semi-Purified Diet, 57W5, with 30% pectin. Dyed blue.

Intended for rodents in a laboratory setting.

CAUTION: Contains a new animal drug for investigational use only in laboratory research animals or for tests in vitro. Not for use in humans.

Storage conditions are particularly critical to TestDiet® products, due to the absence of antioxidants or preservative agents. To provide maximum protection against possible changes during storage, store in a dry, cool location. Storage under refrigeration (2° C) is recommended. Maximum shelf life is six months. (If long term studies are involved, storing the diet at -20° C or colder may prolong shelf life.) Be certain to keep in air tight containers.

| ***Product Forms Available**** | ***Catalog #*** |
|---|---|
| 1/2" Pellet, Irradiated | 1819676-203 |

**Other Forms Available On Request*

##### INGREDIENTS (%)

| | |
|---|---|
| Pectin | 30.0000 |
| Casein - Vitamin Tested | 20.0000 |
| Maltodextrin | 13.2000 |
| Sucrose | 10.0000 |
| Corn Starch | 9.7186 |
| Soybean Oil | 7.0000 |
| Powdered Cellulose | 5.0000 |
| AIN 93G Mineral Mix | 3.5000 |
| AIN 93 Vitamin Mix | 1.0000 |
| L-Cystine | 0.3000 |
| Choline Bitartrate | 0.2500 |
| FD&C Blue No. 2 | 0.0300 |
| t-Butylhydroquinone | 0.0014 |

##### NUTRITIONAL PROFILE [1]

| | |
|---|---|
| ***Protein, %*** | ***18.0*** |
| Arginine, % | 0.70 |
| Histidine, % | 0.52 |
| Isoleucine, % | 0.96 |
| Leucine, % | 1.73 |
| Lysine, % | 1.45 |
| Methionine, % | 0.52 |
| Cystine, % | 0.37 |
| Phenylalanine, % | 0.96 |
| Tyrosine, % | 1.01 |
| Threonine, % | 0.77 |
| Tryptophan, % | 0.22 |
| Valine, % | 1.14 |
| Alanine, % | 0.55 |
| Aspartic Acid, % | 1.29 |
| Glutamic Acid, % | 4.08 |
| Glycine, % | 0.39 |
| Proline, % | 2.36 |
| Serine, % | 1.10 |
| Taurine, % | 0.00 |
| ***Fat, %*** | ***7.0*** |
| Cholesterol, ppm | 0 |
| Linoleic Acid, % | 3.58 |
| Linolenic Acid, % | 0.55 |
| Arachidonic Acid, % | 0.00 |
| Omega-3 Fatty Acids, % | 0.55 |
| Total Saturated Fatty A | 1.05 |
| Total Monounsaturated Fatty Acids, % | 1.54 |
| Polyunsaturated Fatty Acids, % | 3.78 |
| ***Fiber (max), %*** | ***35.0*** |
| ***Carbohydrates, %*** | ***33.2*** |
| ***Energy (kcal/g)*** [2] | ***2.68*** |

| From: | kcal | % |
|---|---|---|
| Protein | 0.719 | 26.9 |
| Fat (ether extract) | 0.632 | 23.6 |
| Carbohydrates | 1.327 | 49.6 |

| | |
|---|---|
| ***Minerals*** | |
| Calcium, % | 0.51 |
| Phosphorus, % | 0.32 |
| Potassium, % | 0.36 |
| Magnesium, % | 0.05 |
| Sodium, % | 0.12 |
| Chloride, % | 0.21 |
| Fluorine, ppm | 1.0 |
| Iron, ppm | 38 |
| Zinc, ppm | 35 |
| Manganese, ppm | 11 |
| Copper, ppm | 6.0 |
| Cobalt, ppm | 0.0 |
| Iodine, ppm | 0.21 |
| Chromium (added), ppm | 1.0 |
| Molybdenum, ppm | 0.14 |
| Selenium, ppm | 0.24 |
| ***Vitamins*** | |
| Vitamin A, IU/g | 4.0 |
| Vitamin D-3 (added), IU/g | 1.0 |
| Vitamin E, IU/kg | 81.6 |
| Vitamin K, ppm | 0.75 |
| Thiamin, ppm | 4.8 |
| Riboflavin, ppm | 6.7 |
| Niacin, ppm | 30 |
| Pantothenic Acid, ppm | 16 |
| Folic Acid, ppm | 2.1 |
| Pyridoxine, ppm | 5.8 |
| Biotin, ppm | 0.2 |
| Vitamin B-12, mcg/kg | 28 |
| Choline Chloride, ppm | 1,250 |
| Ascorbic Acid, ppm | 0.0 |

1. Formulation based on calculated values from the latest ingredient analysis information. Since nutrient composition of natural ingredients varies and some nutrient loss will occur due to manufacturing processes, analysis will differ accordingly. Nutrients expressed as percent of ration on an As-Fed basis except where otherwise indicated.

2. [illegible]
